# Single-molecule identification of full-length proteins with single-amino-acid resolution using nanopores

**DOI:** 10.64898/2026.01.08.698148

**Authors:** Andrea Bonini, Chunzhe Lu, Luca Mantovanelli, Roderick Corstiaan Abraham Versloot, Alwin Jansen, Peter O’Connell Stack, Vasiliki Tsousi, Pascal Knecht, Kathrin Lang, Andrew Heron, Giovanni Maglia

**Author notes:** These authors contributed equally to this work (Andrea Bonini, Chunzhe Lu).

## Abstract

Nanopore-based technologies show promise in single-molecule protein sequencing. By using an unfoldase and a nanopore with enhanced electroosmotic flow, here we show the continuous identification of generic proteins during single nanopore passes. This approach enables the recording of differences in charge and size from single amino acid substitutions compared to reference signals, paving the way for single-molecule protein sequencing and high-throughput proteomics.

## MAIN

Nanopore-based technologies are emerging as a promising, portable, and high-throughput approach for single-molecule sequencing of full-length proteins. This holds the potential to address longstanding challenges in protein analysis, including detecting heterogeneity in post-translational modifications, quantifying low-abundance proteins, and characterizing protein splicing. By tackling these challenges, nanopore technologies will complement mass spectrometry-based methods, accelerating the discovery of insights into the human proteome [^1^].

Previous work showed that enzymes may assist the transport of polypeptides across nanopores [^2– 4^]. More recently, a multi-step method has been described that addressed unstructured or destabilized proteins, showing the potential of nanopores to address single-amino-acid in idealized polypeptide strands and enabling the characterization of post-translational modifications such as phosphorylation [^5^].

Despite these advances, the challenge of dealing with folded proteins remains largely unresolved, as does developing continuous measurements, which are likely required for high-throughput proteomic analysis.

To address these challenges, we developed an approach that uses an unfoldase and a nanopore with an enhanced electroosmotic flow (EOF)[^6^]. The system includes a C-terminal extension on the protein of interest (POI). The latter was designed to ensure an efficient capture, to enable the unfoldase to recognize the POI and to begin unfolding the POI only at the nanopore. Both the unfoldase and the substrate are free to diffuse in the same *cis* solution, enabling continuous measurement.

As for unfoldase, we chose the ATP-dependent motor ClpX for its ability to unravel proteins and move amino acids processively [^7^]. ClpX was also previously used in nanopore experiments [^5,8,9^]. Optimization of the buffer conditions for nanopore experiments (Figure S1) revealed that 1M potassium gluconate (KGlu) allows the enzyme to remain functional while maintaining a high concentration of [K^+^] required for cation selective nanopores (Figure S2).

We introduced a polypeptide tag (sequencing tag, SQtag) at the C-terminus of the POI that included a stalling domain (SD), which was designed to inhibit the unfolding of the POI in solution. The SD included a modified leucine zipper motif [^10^], composed of repeated leucine residues and introduced glutamate and arginine residues across two distinct alpha-helical regions, connected by a flexible linker (Figure 1a). These regions interact via leucine-leucine weak hydrophobic interactions and electrostatic interactions, forming a coiled coil structure [^11^]. The stalling domain was followed by a positively charged region to enhance electrophoretic capture of protein, and a ssrA tag for ClpX recognition (Figure 1a). As an initial test we used a ClpX unfoldase, and maltose-binding protein (MBP) elongated with the C-terminal sequencing tag (MBP-SQtag).

**Figure 1.**
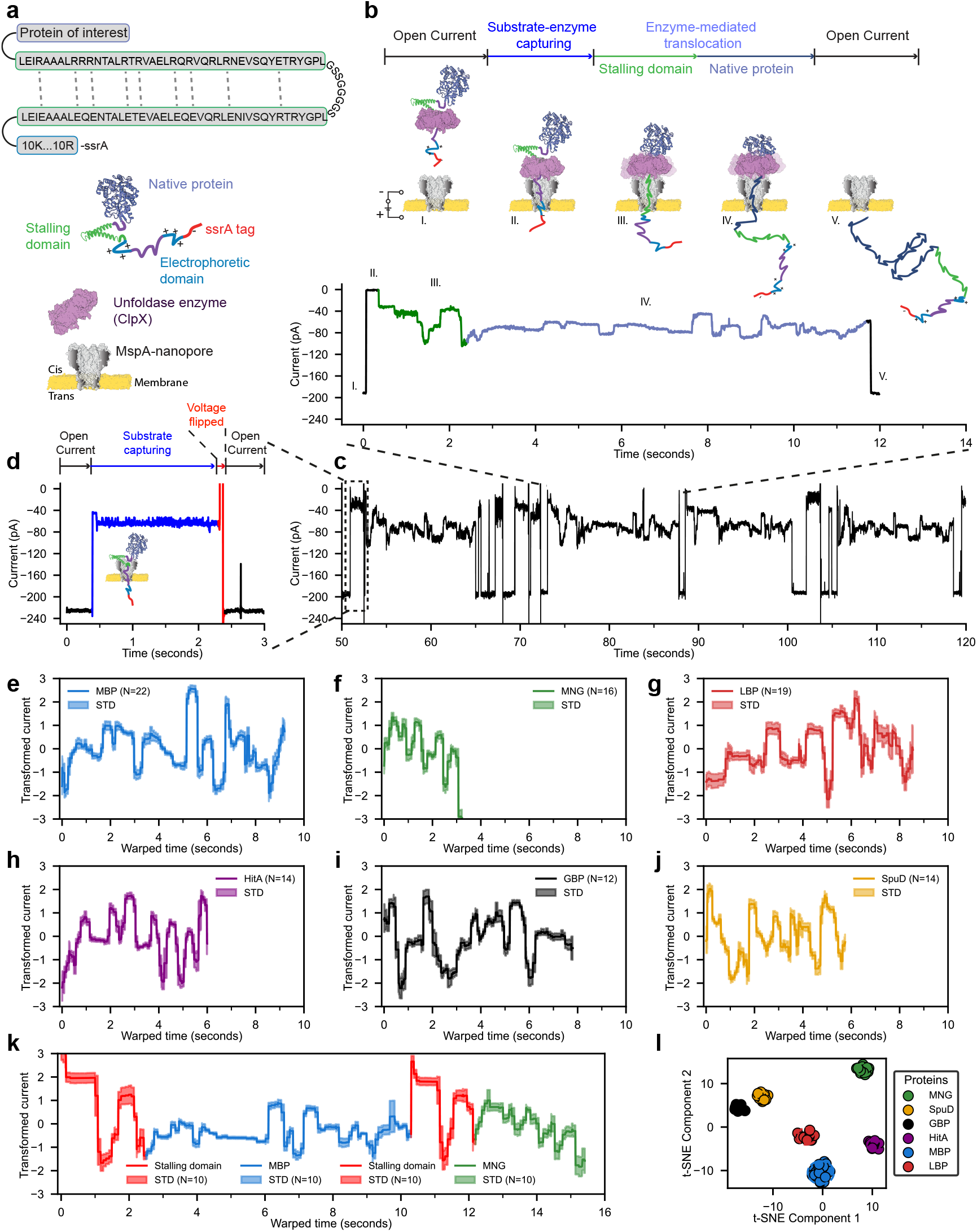
Unfoldase-mediated nanopore protein identification. **(a)** Elements of nanopore-based strategies for protein identification. (**b)** Schematic of cis-to-trans protein translocation mediated by ClpX. (**c)** Extract of real-time trace showing sequential squiggles during protein translocation. (**d)** Single block event from a protein captured without unfoldase. Ensemble aligned traces to the medoid for various proteins, showing only the protein region without the SQtag signal: (**e**) maltose binding protein, MBP-SQtag, **(f)** mNeoGreen, MNG-SQ-tag, **(g) L-**leucine-binding protein, LBP-SQtag, (**h)**, ferric iron-binding periplasmic protein, HitA-SQtag, (**i)** glucose/galactose binding protein, GBP-SQtag **(j)** putrescine-binding periplasmic protein, SpuD-SQtag. (**k)** Ensemble aligned traces to the medoid of a conjugate MNG-SD-MBP-SQtag. (**I)** t-SNE visualization of DTW distances between traces for each protein. All traces were collected at -65 mV at 37 °C in 1 M KGlu, 50 mM HEPES, 10 mM MgCl_2_, pH 7.4, in the presence of 5 nM of POI, 100 nM of ClpX, 2 mM of ATP, 1.6 mM creatine phosphate, 0.4 µM creatine kinase, 1 mM DTT, and 0.5 mM EDTA.

Among the highly cation-selective nanopores capable of translocating unstructured proteins such as CytK-4D, MspA, and Lysenin [^6,12^], we selected the MspA pore for this study. Nanopore experiments were carried out at 37 °C, the optimal working temperature for ClpX, with an applied voltage ranging from -60 to -65 mV (Figure S3 and additional text S1).

Initially, MspA was preferred over other nanopores because of its well defined and narrow sensing region [^12^]. Several enzyme-mediated translocation events, termed as “squiggle”, were observed (Figure 1b), showing the translocation of the substrate proteins from the *cis* to the *trans* side and allowing the nanopore to continuously capture substrates (Figure 1c). By contrast, when the substrate is captured without the unfoldase, a permanent current block is observed (Figure 1d). When MBP was elongated only with a ssrA tag (MBP-ssrA), mainly partial squiggles are observed (Figure S4) confirming the importance of the stalling domain in the sequencing tag (additional text S2).

Each squiggle showed reproducible step-size current features corresponding to the processive feeding of the unfolded polypeptide across the nanopore mediated by the enzyme (additional text S3). Specific current levels could be associated to the capturing of the charged region in the sequencing tag (Figure 1b, (II)), the processing of the stalling domain (Figure 1b, (III)), and the processing of MBP (Figure 1b, (IV), Figure S6).

In CytK-4D we also recorded squiggles similar to MspA. However, the squiggles showed fewer steps and a narrower ionic current range (Figure S7), reflecting the longer transmembrane sensing region of CytK-4D compared to MspA. Interestingly, MspA also provided more informative signals, covering a longer portion of the protein at the N-terminus, compared to CytK-4D (Figure S7).

The ability of the nanopore signal to distinguish between proteins was tested using six natural proteins containing at the C-terminus the sequencing tag (Figure 1). Each protein showed a reproducible signal with several current steps, demonstrating the nanopore’s ability to generate a specific step-based protein profile associated with each protein sequence (Figure 1e-j). Comparison of individual traces using a dynamic time warping algorithm and classification using t-SNE clustering revealed a clear distinction among the different proteins (Figure 1| and S8), indicating that current fingerprints can likely be used to identify proteins during single passes.

A concatemer protein composed by mNeoGreen – stalling domain - maltose binding protein-sequencing tag (MNG-SD-MBP-SQtag) was also measured to elucidate the full MBP and the SD tag sequence, and to demonstrate the potential to process longer proteins (Figure 1k). The ability of ClpX to process and unfold the proteins was concurrently confirmed by a degradation assay and a bulk unfolding assay (Figure S9).

To understand the signal, we first characterized a designed substrate containing a long unstructured polypeptide (178 amino acids) linking the stalling domain to the native MBP protein (additional text S4). Substitutions of multiple amino acids in the unstructured domain revealed that positively charged amino acids (i.e., two or three consecutive arginine) induced large reduction of the ionic current, most likely reflecting the hindered passage of K+ ions in the cation selective MspA (Figure S10b). By contrast, small-volume residues (i.e., two glycines) resulted in significantly increased current signals, most likely reflecting the smaller excluded volume of glycine sidechains (Figure S10a). Substitutions with multiple aromatic (WW or YY) or negatively charged residues (DD) induced intermediate blockades of the ionic currents (Figure S1a), most likely reflecting the increased excluded volume of the amino acid side chains. A similar behavior was also observed with multi-substitutions in the folded MBP domain at different positions (Figure S10 c,d, additional text S4).

Given that substitutions with positively charged amino acids gave the largest signal changes, to map specific positions of the MBP sequence (Figure 2a and S11), we introduced and/or removed single arginine or lysine residues (Figures 2b-f). As shown in Figures 2b, c, d, and e, each mutation resulted in a clear, statistically distinguishable change in the current signal compared to the unmodified protein (reference signal). A clear link between the sequence and the current signal in MBP could then be obtained. The substitution of an arginine in a L-leucine binding protein resulted in the same current reduction observed with MBP (Figure S12), suggesting this behavior is generic, and that proteins might be fingerprinted by mapping their charge distribution.

**Figure 2.**
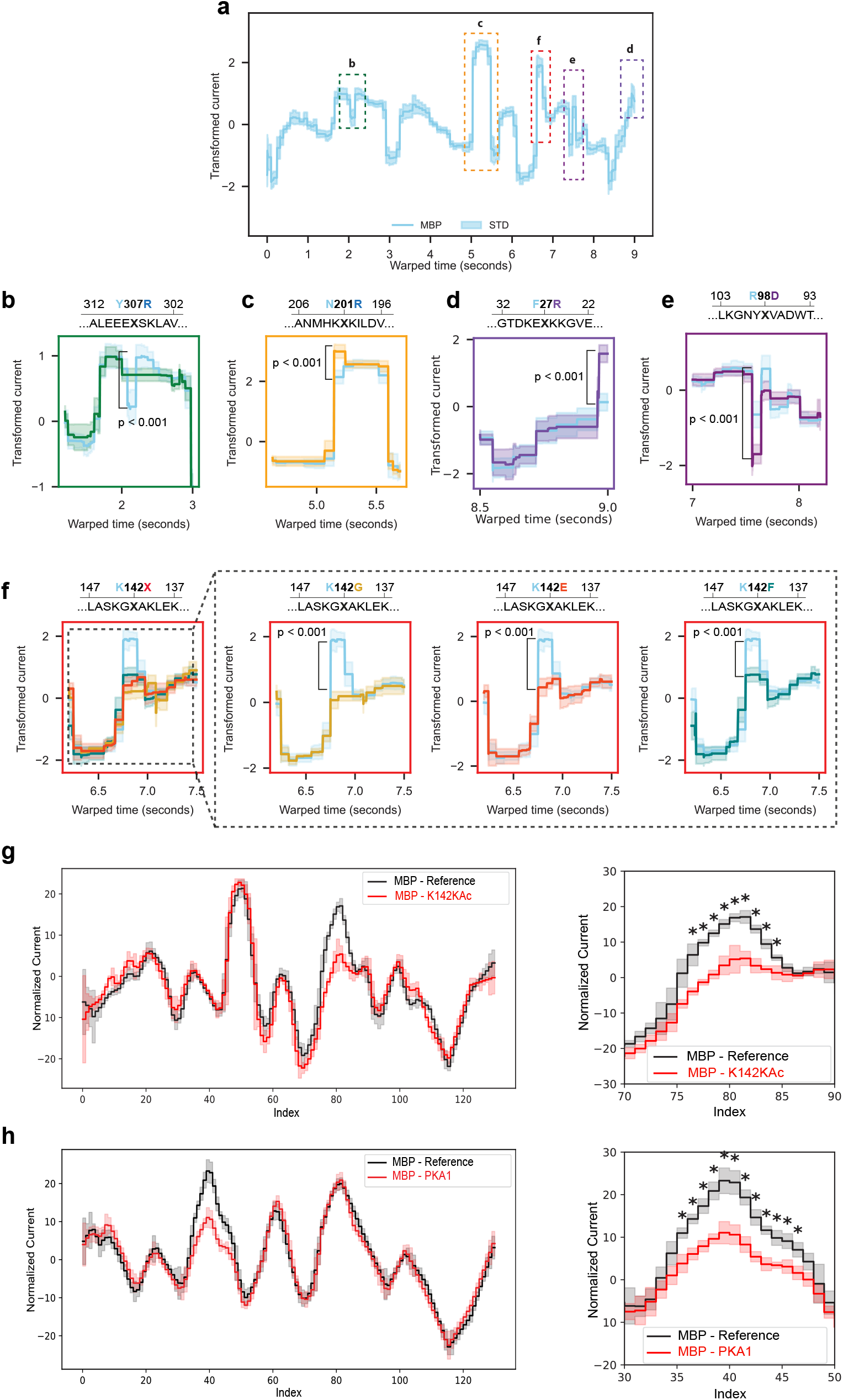
Single amino acid substitution in protein domain. (a) reference trace of MBP showing the positions of single point mutations along the trace, highlighted by rectangles. (b-f) aligned traces displaying only the regions where mutations significantly affected the signal (t-test, point-by-point analysis). The full-length alignment between point mutants and the wild-type protein is illustrated in Fig. S10. Mutations introducing positively charged residues are shown in (b) Y307R (N=17), (c) N201R (N=9), and (d) F27R (N=10). Mutation removing positively charged residues is shown in (e) R98D (N=10). (f) Overlay of all mutations at position K142, including K142G (N=9), MBP_K142E (N=23), and MBP_K142F (N=15)). (g) Alignment between MBP reference (N = 29) and acetylated MBP-K142KAc (N = 21). (h) Alignment between MBP reference (N = 60) and phosphorylated MBP-PKA1 (N = 61). All traces were collected at -60 to -65 mV at 37°C in 1 M KGlu, 50 mM HEPES, 10 mM MgCl_2_, pH 7.4, in the presence of 5-20 nM of POI, 100 nM of ClpX, 2 mM of ATP, 1.6 mM creatine phosphate, 0.4 µM creatine kinase, 1 mM DTT, and 0.5 mM EDTA.

Notably, the MBP-F27R mutation (Figure 2d) showed a change at the very end of the MBP signal, suggesting that after the nanopore reads this residue, ClpX releases MBP atop of the nanopore. Hence, this method can read a window of amino acids from the C-terminus of the POI to near the end of the N-terminus sequence, missing the last ∼50 residues (additional text S5). Based on this information, we divided the number of amino acids read by the number of steps observed in the squiggles, yielding an average of approximately 4 ± 1 amino acid per enzyme step across all six protein domains measured (Table S3, additional text S6). This value is consistent with step sizes measured by other techniques, such as single-molecule optical tweezers [^13^], and differs from a recently reported 2–amino-acid step size derived from experiments on unstructured synthetic polypeptides [5]. Overall, this largely depends on the specific conditions of the system (bufferconditions, sensing region of the pore, protein structure, etc.) as well as the method used to identify steps in the traces (additional text S6).

Next, we investigated the effect of single substitutions of different amino acids within the same position (Figure 2f) and two of the most common post translational modifications (Figure 2 g, and h). At position 142 of MBP, we substituted a lysine residue with smaller-volume (glycine), negatively charged (aspartate), and aromatic (phenylalanine) residues. The current levels varied accordingly, from high (lysine) to low blockage (glycine) with intermediate levels for aspartate and phenylalanine, reflecting the effects of residue volume and charge on the nanopore signal. The system’s ability to detect PTMs was tested by using genetic code expansion to introduce acetyllysine site-specifically at position 142 of MBP (MBP-^K142KAc^), Figure 2g and Figure S13).

Comparison of the time-warped ionic current traces (Figure 2g), indicated that acetyllysine increased the ionic current, reflecting the removal of the positive charge otherwise present in lysine. The effect of phosphorylation was tested by producing a MBP variant (MBP^PKA1^) that contained a site for phosphorylation by cAMP dependent protein kinase (PKA). Phosphorylated MBP^PKA1^ showed an increased signal compared to unphosphorylated MBP^PKA1^ (Figure 2h and Figure S14), suggesting that the additional negative charge further increased the ion selectivity of the nanopore during the transient occupancy of the phosphate group at the constriction of the nanopore.

It is worth noting that both acetylation and phosphorylation increased the ionic current despite these modifications increasing the size of the amino acid side chain. This suggests that alterations of the charge of the side chain have a larger impact on the ionic current signal than a small (<100 Da) modification of their side chain.

In summary, we present a novel nanopore-based approach capable of continuously measuring native proteins at the single-molecule level. This technique provides reproducible step-size signals from individual proteins. Protein squiggles showed unique and reproducible signals that were affected by both the charge and size of individual amino acids and allowed for the identification of phosphorylation and acetylation in single amino acids. Proteins could be distinguished after a single pass, indicating this method could already be utilized for protein identification and PTM characterization in complex media using prerecorded signals. As our understanding of these signals improves, particularly through the integration of AI-driven algorithms, we anticipate a transition from reference-based identification to *de novo* sequencing applications.

## Acknowledgements

The authors A.B., C.L. and G.M., disclose support for the research of this work from Funder NWO-VICI [grant number 192.068] and Funder NIH [grant 1R01HG012554].

## METHODS

### Chemicals, enzymes, and reagents

The chemicals and corresponding suppliers are listed as follows: DNA oligos and genes used were ordered from Integrated DNA Technologies. Phusion U DNA polymerase, dNTPs, GeneRuler 1kb Plus DNA Ladder, Ethidium Bromide Dye, PageRuler™ Prestained Protein Ladder (10-180 kDa), GeneJET PCR Purification Kit, GeneJET Gel Extraction Kit, GeneJET Plasmid Miniprep Kit, potassium trifluoroacetate (≥98%), and protease inhibitors (Pierce Protease Inhibitor Mini Tablets, EDTA-free) were purchased from Thermo Fisher Scientific. USER enzyme was purchased from New England Biolabs. Mix2Seq Sequencing Kit was purchased from Eurofins. Agar, agarose, arabinose, glucose, LB medium, 2×YT medium, magnesium chloride, ethylenediaminetetraacetic acid (EDTA), potassium gluconate (≥99%), potassium chloride (≥99%), potassium sulfate (≥99%), HEPES (≥99%), n-dodecyl-beta-maltoside (DDM), imidazole (≥99%), NaCl (≥99.5%), isopropyl-β-d-thiogalactoside (≥99.0%) were purchased from Roth. Creatine phosphate disodium salt, creatine kinase (from rabbit muscle), chloramphenicol (≥98%), potassium acetate (99%), potassium nitrate (99%), n-pentane, adenosine 5^′^-triphosphate (ATP) disodium salt hydrate and Adenosine 5^′^-[γ-thio]triphosphate tetralithium salt (ATP-γ-S) were purchased from Sigma-Aldrich. Ampicillin sodium salt was purchased from Fisher Bio Reagents. Ni-NTA agarose was ordered from Qiagen. Desthiobiotin and Strep-Tactin® beads were ordered from IBA Lifesciences. Quick Start Bradford 1× Dye Reagent and Bio-Spin Disposable Chromatography Columns were purchased from Bio-Rad. DPhPC was purchased from Avanti Polar Lipids. n-Hexadecane (99%) were purchased from Acros Organics. Pentane (99%) from ThermoFisher Scientific. Block polymers including PBD_22_PEO_14_ (Poly(1,2-butadiene)-b-(ethylene oxide), Mn (g/mol): 1200-b-600, P41745-BdEO), and PBD_11_PEO_8_ (Poly(1,2-butadiene)-b-(ethylene oxide), with a purity grade of PBD-PEO samples contained >85% 1,2-butadiene (thus <15% 1,4-butadiene) were purchased from Polymer Source, Inc.

### Expression and purification of MspA and CytK-4D nanopores

BL21(DE3) electrical competence cells were transformed with the plasmid pT7sc1-MspA using a Bio-Rad Micropulser (bacterial setting) and selected on LB agar plates containing 100 μg mL-1 ampicillin. The next day, a single colony was inoculated in 10 mL LB medium with 100 μg mL-1 ampicillin and cultivated overnight at 37 °C and 200 rpm. The day after, this overnight culture was used to inoculate 200 mL of LB medium at an initial OD600 of 0.1 and cultivated at 37 °C and 200 rpm. Once the optical density reached 0.6-0.8, the culture was placed on ice for 5 min before 0.5 mM IPTG was added to induce protein expression. After 18 h expression at 20 °C, 200 rpm, the cells were harvested (8,000 rpm, 10 min). Cell pellets were frozen at -80 °C for 2 h and then resuspend in 20 mL of ice-cold lysis buffer (50 mM Tris-HCl pH 7.5, 150 mM NaCl, 0.02% DDM, and half tablet of cocktail protease inhibitors). The cell resuspension was first sonicated (Branson 450 Sonifier) at 20% duty cycle and 3 output control for 5 min and then spun down at 8,000 rpm for 30 min. A 300 μL of Strep-Tactin® slurry (50% suspension) that was prewashed three times with lysis buffer was added into the supernatant and incubated for 20 min. The mixture was loaded into columns (1 mL bio-spin chromatography, Bio-Rad), which allow the follow-through pass by gravity. Afterwards, the beads bed was prewashed with a 5-10 mL wash buffer (50 mM Tris-HCl pH 7.5, 150 mM NaCl, 0.02% DDM). MspA proteins were eluted with 300 μL of elution buffer (50 mM Tris-HCl pH 7.5, 150 mM NaCl, 0.02% DDM, 2.5 mM desthiobiotin) in two fractions. MspA oligomers were confirmed by SDS-PAGE without heating step during sample preparation.

CytK-4D nanopore was expressed and purified according to the published protocol [^6^].

### Cloning and production of substrate proteins and mutants

The DNA sequences of mNeoGreen (mNG), maltose-binding protein (MBP), glucose/galactose-binding protein (GBP), leucine-binding protein (LBP), ferric iron-binding periplasmic protein (HitA), putrescine-binding periplasmic protein (SpuD), and stalling domain (consisting of an engineered leucine-zipper + linker region + electrophoretic tag) were ordered from Integrated DNA technologies (IDT, USA). To simplify the protein purification procedure, the periplasmic signal peptide sequences of these proteins were predicted by using the SignalP-6.0 server (https://services.healthtech.dtu.dk/services/SignalP-6.0/) and removed during plasmid construction using USER cloning. Uracil-containing primers were ordered from Integrated DNA technologies (IDT, USA) and used to amplify substrate protein genes and pT7sc1 vector backbone using Phusion U Hot Start DNA polymerase (ThermoFisher Scientific). PCR reactions were run according to the manufacturer’s instructions. The PCR products were cleaned up by using either GeneJET PCR Purification Kit or GeneJET Gel Extraction Kit (ThermoFisher Scientific). The gene fragments and the linearized pT7sc1 vector were added in a molar ratio of 3:1 and follow the standard USER reaction and program [^14^]. The USER reaction mixtures were transformed into chemically competent *E. coli c*ells using the heat shock method. The cells were selected on an LB agar plate with 100 μg mL-1 ampicillin. Single colonies were picked up and inoculated in 10 mL LB medium for plasmids extraction. The successful plasmid constructions were confirmed by Sanger sequencing (Eurofins). The verified plasmids were transformed into the SG1146a competence cells for substrate protein production. Due to the deletion of ClpP protease in SG1146a expression strain, degradation of ssrA-containing protein was diminished [^15^]. Protein purification was performed as the method described for MspA nanopore with minor modifications. Lysis and wash buffers (50 mM HEPES, 150 mM NaCl, 1 mM DTT, pH7.5) were used. 5 mM Desthiobiothin was added into the wash buffer, resulting in an elution buffer. The presence of proteins was confirmed by SDS-PAGE. Proteins were desalted and concentrated using Amicon® Ultra Centrifugal Filter (10 kDa MWCO) and flash frozen in storage buffer (50 mM HEPES, pH 7.5, 150 mM NaCl, 1 mM DTT, 10% glycerol) and stored at -20 °C. Point mutants were introduced into MBP and LBP protein by USER cloning as stated previously. Protein purification of the mutants was performed the same as wild type proteins.

### Protein sequences of substrates and sequencing tag

#### Sequencing tag

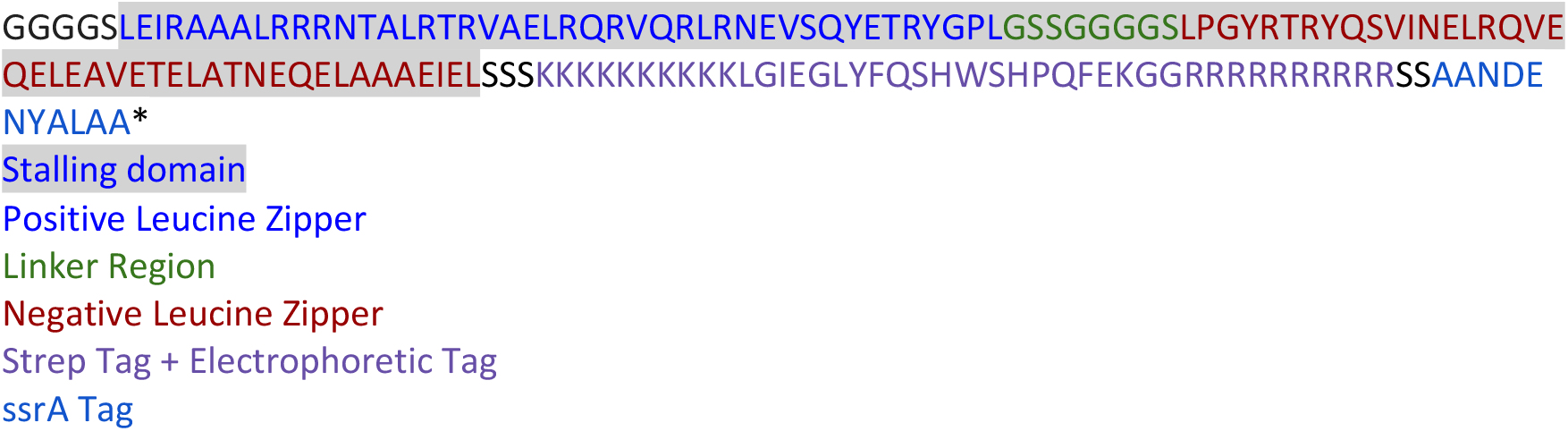

#### mNG

VSKGEEDNMASLPATHELHIFGSINGVDFDMVGQGTGNPNDGYEELNLKSTKGDLQFSPWILVPHIGYGFHQYLPYPDGMSPFQAAMVDGSGYQVHRTMQFEDGASLTVNYRYTYEGSHIKGEAQVKGTGFPADGPVMTNSLTAADWCRSKKTYPNDKTIISTFKWSYTTGNGKRYRSTARTTYTFAKPMAANYLKNQPMYVFRKTELKHSKTELNFKEWQKAFTDVMGMDELYK

#### MBP

KIEEGKLVIWINGDKGYNGLAEVGKKFEKDTGIKVTVEHPDKLEEKFPQVAATGDGPDIIFWAHDRFGGYAQSGLLAEITPDKAFQDKLYPFTWDAVRYNGKLIAYPIAVEALSLIYNKDLLPNPPKTWEEIPALDKELKAKGKSALMFNLQEPYFTWPLIAADGGYAFKYENGKYDIKDVGVDNAGAKAGLTFLVDLIKNKHMNADTDYSIAEAAFNKGETAMTINGPWAWSNIDTSKVNYGVTVLPTFKGQPSKPFVGVLSAGINAASPNKELAKEFLENYLLTDEGLEAVNKDKPLGAVALKSYEEELAKDPRIAATMENAQKGEIMPNIPQMSAFWYAVRTAVINAASGRQTVDEALKDAQT

#### GBP

ADTRIGVTIYKYDDNFMSVVRKAIEQDAKAAPDVQLLMNDSQNDQSKQNDQIDVLLAKGVKALAINLVDPAAAGTVIEKARGQNVPVVFFNKEPSRKALDSYDKAYYVGTDSKESGIIQGDLIAKHWAANQGWDLNKDGQIQFVLLKGEPGHPDAEARTTYVIKELNDKGIKTEQLQLDTAMWDTAQAKDKMDAWLSGPNANKIEVVIANNDAMAMGAVEALKAHNKSSIPVFGVDALPEALALVKSGALAGTVLNDANNQAKATFDLAKNLADGKGAADGTNWKIDNKVVRVPYVGVDKDNLAEFSKK

#### LBP

DDIKVAVVGAMSGPIAQWGDMEFNGARQAIKDINAKGGIKGDKLVGVEYDDACDPKQAVAVANKIVNDGIKYVIGHLCSSSTQPASDIYEDEGILMISPGATNPELTQRGYQHIMRTAGLDSSQGPTAAKYILETVKPQRIAIIHDKQQYGEGLARSVQDGLKAANANVVFFDGITAGEKDFSALIARLKKENIDFVYYGGYYPEMGQMLRQARSVGLKTQFMGPEGVGNASLSNIAGDAAEGMLVTMPKRYDQDPANQGIVDALKADKKDPSGPYVWITYAAVQSLATALERTGSDEPLALVKDLKANGANTVIGPLNWDEKGDLKGFDFGVFQWHADGSSTAAK

#### HitA

AALEVLFQGPGYQDPNSMDPVTLTLYNGQHAATGIAIAKAFQDKTGIQVKIRKGGDGQLASQITEEGARSPADVLYTEESPPLIRLASAGLLAKLEPETLALVEPEHAGGNGDWIGITARTRVLAYNPKKIDEKDLPKSLMDLSDPSWSGRFGFVPTSGAFLEQVAAVIKLKGQEEAEDWLTGLKAFGSIYTNNVTAMKAVENGEVDMALINNYYWYTLKKEKGELNSRLHYFGNQDPGALVTVSGAAVLKSSKHPREAQQFVAFMLSEEGQKAILSQSAEYPMRKGMQADPALKPFAELDPPKLTPADLGEASEALSLERDVGLN

#### SpuD

ADNKVLHVYNWSDYIAPDTLEKFTKETGIKVVYDVYDSNEVLEAKLLAGKSGYDVVVPSNSFLAKQIKAGVYQKLDKSKLPNWKNLNKDLMHTLEVSDPGNEHAIPYMWGTIGIGYNPDKVKAAFGDNAPVDSWDLVFKPENIQKLKQCGVSFLDSPTEILPAALHYLGYKPDTDNPKELKAAEELFLKIRPYVTYFHSSKYISDLANGNICVAIGYSGDIYQAKSRAEEAKNKVTVKYNIPKEGAGSFFDMVAIPKDAENTEGALAFVNFLMKPEIMAEITDVVQFPNGNAAATPLVSEAIRNDPGIYPSEEVMKKLYTFPDLPAKTQRAMTRSWTKIKSGK

#### mNG-SD-MBP

VSKGEEDNMASLPATHELHIFGSINGVDFDMVGQGTGNPNDGYEELNLKSTKGDLQFSPWILVPHIGYGFHQYLPYPDGMSPFQAAMVDGSGYQVHRTMQFEDGASLTVNYRYTYEGSHIKGEAQVKGTGFPADGPVMTNSLTAADWCRSKKTYPNDKTIISTFKWSYTTGNGKRYRSTARTTYTFAKPMAANYLKNQPMYVFRKTELKHSKTELNFKEWQKAFTDVMGMDELYKNGGGGSLEIRAAALRRRNTALRTRVAELRQRVQRLRNEVSQYETRYGPLGSSGGGGSLPGYRTRYQSVINELRQVEQELEAVETELATNEQELAAAEIELSSSKKKKKKKKKKLGIEGLYFQSHGGKIEEGKLVIWINGDKGYNGLAEVGKKFEKDTGIKVTVEHPDKLEEKFPQVAATGDGPDIIFWAHDRFGGYAQSGLLAEITPDKAFQDKLYPFTWDAVRYNGKLIAYPIAVEALSLIYNKDLLPNPPKTWEEIPALDKELKAKGKSALMFNLQEPYFTWPLIAADGGYAFKYENGKYDIKDVGVDNAGAKAGLTFLVDLIKNKHMNADTDYSIAEAAFNKGETAMTINGPWAWSNIDTSKVNYGVTVLPTFKGQPSKPFVGVLSAGINAASPNKELAKEFLENYLLTDEGLEAVNKDKPLGAVALKSYEEELAKDPRIAATMENAQKGEIMPNIPQMSAFWYAVRTAVINAASGRQTVDEALKDAQTN

### Cloning and production of MBP-^K142KAc^

The site-specifically acetylated MBP was achieved using a genetic code expansion approach. Briefly, a premature amber (TAG) codon was introduced into MBP-K142 using site-directed mutagenesis by PCR-driven overlap extension, resulting in pBAD_MBP-K142TAG. Chemically competent *E. coli* K12ΔclpP cells were transformed with pBAD_MBP-K142TAG and pEVOL_AcKRS3, a plasmid containing *M. barkeri* PylRS mutant [^16^] known to incorporate acetyllysine (AcK) under a constitutive GlnS promoter and an arabinose inducible promoter with pyrrolysyl-tRNA (PylT) under a constitutive promoter. After recovery with 1 mL SOC medium for 1 hour at 37 °C, the cells were cultured overnight in 5 mL 2xYT medium containing both 100 μg mL-1 ampicillin and 50 μg mL-1 chloramphenicol at 37 °C, 200 rpm. The overnight culture was diluted to an OD600 of 0.05 in 50 mL of fresh 2xYT medium containing the half concentration of corresponding antibiotics. For acetyllysine incorporation, the expression culture was further supplemented with acetyllysine (10 mM) and nicotinamide (NAM, 20 mM). The culture was incubated at 37 °C, 200 rpm, until OD600 reached 0.6, at which point protein expression was induced by addition of 0.05% arabinose (w/v). The culture was further incubated at 37 °C, 200 rpm, for 16 hours. The cells were harvested by centrifugation (4000xg, 20 min, 4 °C) and resuspended in 20 mL lysis buffer (20 mM Tris pH 7.5 at 4 °C, 150 mM NaCl) containing cOmplete™ protease inhibitor (Roche, one tablet per 50 mL). The lysis buffer further contained 20 mM NAM for the acetyllysine containing protein. The cells were lysed by sonication with cooling in an ice-water bath. The lysed cells were centrifuged (14000xg, 20 min, 4 °C) and the supernatant was incubated with Strep beads (Strep-Tactin® MacroPrep® resin from IBA Lifesciences) for 4 hours at 4 °C with agitation. After incubation, the Strep beads were collected on a gravity column and washed with 7 column volumes (CV) of wash buffer (50 mM Tris pH 7.5 at 4 °C, 150 mM NaCl). The protein was eluted with a wash buffer supplemented with 2.5 mM desthiobiotin. The elution was rebuffered into a storage buffer (20 mM Tris pH 7.5 at 4 °C, 100 mM NaCl) by dialysis using a Pur-A-Lyzer™ Maxi 6000 dialysis unit. The protein was concentrated using an Amicon® centrifugal filter unit 30 kDa MWCO (Millipore). Protein concentrations were determined using a NanoPhotometer® N60 (IMPLEN). Extinction coefficients were calculated with ProtParam (https://web.expasy.org/protparam/). The purified protein was flash frozen in aliquots using liquid nitrogen and stored at -80 °C until further use. Acetylated MBP was verified by Liquid Chromatography-Mass Spectrometry (LC-MS) analysis. This was performed on a Bruker-Compact (Q-TOF MS) with chromatographic separation on an Acquity UPLC® Protein BEH C4 column (2.1 x 100 mm, 1.7 µm).

### Purification of MBP^PKA1^

For our phosphorylation study, we designed MBP^PKA1^, a mutant of MBP where we introduced an extended loop with an {RRAS} motive between positions 199 and 201 of MBP. The {RRAS} motive can be recognised by cAMP dependent protein kinase A (PKA) to phosphorylate the serine residue. 50 ng of pET28a plasmid containing the gene for MBP^PKA1^ was transformed into E. coli BL21 (DE3) chemically competent cells. The transformed cells were grown into single colonies in lysogeny broth (LB) agar plates with 50 ug/ml kanamycin. A single colony was used to inoculate a 10 mL LB starter culture that grew at 37 °C for 20 hours under constant shaking at 180 rpm. 5 mL of the starter culture was used to inoculate a 400 mL LB culture. When the OD600 reached 0.7, 0.1 mM of IPTG was added to induce protein production and the cells were grown at 25 °C for 20 hours under constant shaking at 180 rpm. Cells were harvested by centrifugation at 1900 g for 30 min at 4 °C and resuspended in 50 mM Tris-HCl, 150 mM NaCl, 1 mM EDTA, 20 mM imidazole, pH 7.5 (lysis buffer) supplemented with 1 tablet of protease inhibitors and 4 units of DNase I. The cells were lysed by sonication on ice for 12 min at 18% intensity (7 min on/ 7 min off) using a Branson Sonifier SFX 550 and afterwards centrifuged at 7500 g for 30 min at 4 °C. The supernatant was filtered with syringe filters with a pore size of 0.45 µm and incubated with 1 mL of pre-washed Ni-NTA agarose beads. The beads were washed with 10 mL lysis buffer, and the protein was eluted with 500 mM imidazole. Imidazole was later removed by a buffer exchange to 1x PBS (a 10x PBS solution from VWR was diluted 10 times with MilliQ) using PD10 columns (Cytiva). The protein was stored at -20 °C in the presence of 10% glycerol until use.

### Phosphorylation of MBP^PKA1^

5 µM of MBP^PKA1^ was mixed with 1x NE Buffer for Protein Kinases (PK) from New England Biolabs and 200 µM ATP in a final volume of 100 µL. The mixture was incubated for 30 minutes at 30°C, under constant shaking at 600 rpm. 7500 units of cAMP-dependent Protein Kinase (PKA), catalytic subunit (New England Biolabs) was added, and the mixture was incubated for 2 hours at 30°C, under constant shaking at 600 rpm. Afterwards, a buffer exchange to 1x PBS (a 10x PBS solution from VWR was diluted 10 times with MilliQ) was performed using a PD10 column (Cytiva) and protein was stored at -20°C until use.

Phosphorylation of MBP^PKA1^ was confirmed via Mass Spectrometry analysis (LC-MS/MS). 25 µl of protein sample was mixed with 20 ng of Lys-C and incubated overnight at 37°C. The reaction was afterwards quenched by the addition of 5% formic acid. The samples were analyzed on a LC-MS/MS consisting of a Shimadzu LC20-XR system (Shimadzu Benelux, Den Bosch, The Netherlands) interfaced with a Q-Exactive plus mass spectrometer (Thermo Fisher Scientific, USA).

### Purification of single point mutants of 20 amino acids in MBP

Single point mutations of MBP proteins were produced by GenScript. The proteins were designed to cover all 20 amino acids at position 100 and 201 of MBP. The proteins carry an N-terminal strep-tag and a C-terminal leader domain. Proteins are purified by a single strep-tag purification followed by a buffer exchange to 50 mM HEPES, 20% glycerol, pH 7.4.

### Cloning, expression and purification of ClpX and ClpP

The genes of ClpXdN (residue 62-424) and ClpP were amplified from the genome of *E. coli* K12. As described previously, USER cloning was used to construct pBAD-ClpXdN_Strep and pACYC-ClpP_Strep. The successful constructs were verified by Sanger sequencing (Eurofins). Afterwards, the correct plasmids were transferred into *E. coli* BL21AI expression cells. ClpXdN expression was conducted the same as the previous procedure for substrate protein, except that 1% arabinose was used as an inducer instead of IPTG. For ClpP expression, both 1% arabinose and 0.5 mM IPTG were added. In the case of ClpXdN and ClpP purification, pellets from 100 mL culture were resuspended into a 10 mL lysis buffer (50 mM HEPES, 200 mM KCl, pH 7.4), followed by sonication and purification steps which were same as substrate protein Strep-tag purification. After the elution step, ClpXdN and ClpP proteins were concentrated using Amicon® Ultra Centrifugal Filter (30 kDa MWCO) and flash frozen in storage buffer (50 mM HEPES, pH 7.5, 200 mM KCl, 2 mM DTT, 10% glycerol) and stored at -80 °C.

### Protein sequence of ClpXdN and ClpP

#### ClpXdN

MKHHHHHHGSGENLYFQSRSALPTPHEIRNHLDDYVIGQEQAKKVLAVAVYNHYKRLRNGDTSNGVELGKSNILLIGPTGSGKTLLAETLARLLDVPFTMADATTLTEAGYVGEDVENIIQKLLQKCDYDVQKAQRGIVYIDEIDKISRKSDNPSITRDVSGEGVQQALLKLIEGTVAAVPPQGGRKHPQQEFLQVDTSKILFICGGAFAGLDKVISHRVETGSGIGFGATVKAKSDKASEGELLAQVEPEDLIKFGLIPEFIGRLPVVATLNELSEEALIQILKEPKNALTKQYQALFNLEGVDLEFRDEALDAIAKKAMARKTGARGLRSIVEAALLDTMYDLPSMEDVEKVVIDESVIDGQSKPLLIYGKPEAQQASGEGSSWSHPQFEK*

#### ClpP

MSYSGERDNFAPHMALVPMVIEQTSRGERSFDIYSRLLKERVIFLTGQVEDHMANLIVAQMLFLEAENPEKDIYLYINSPGGVITAGMSIYDTMQFIKPDVSTISMGQAASMGAFLLTAGAKGKRFSLPNSRVMIHQPLGGYQGQATDIEIHAREILKVKGRMNELMALHTGQSLEQIERDTERDRFLSAPEAVEYGLVDSILTHRNGSSWSHPQFEK*

### ClpX unfoldase assay

Fluorescence-based assays were used to test ClpXdN unfoldase activity. 150 µL reactions were prepared by adding 33 nM mNeoGreen containing a ssrA tag at the C-terminus, 250 nM ClpXdN, 250 nM ClpP, and 1xATP regeneration buffer (a final concentration of 50 mM HEPES, 5 mM MgCl2, 16 mM creatine phosphate, 4 µM creatine kinase, 2 mM DTT, 5 mM EDTA). Lastly, different potassium salts at various concentrations (0.2-1 M) were added into the wells. The reaction plate was loaded into a CLARIOstar Plus plate reader (BMG Labtech). Fluorescence emission at 588 nm (±20 nm, low pass filter of 561.2 nm) after excitation at 537 nm (±15 nm) was monitored for 10 min at 37 °C. Prior to the start of monitor, 1 mM ATP was injected into the well to initiate the unfolding. Data was collected using the instrument’s Installation Package V5.60 software before exporting via the MARS V3.10 software.

### Substrate degradation assay

A degradation assay was conducted for substrates containing a stalling domain to verify the presence of ssrA tag and its capabilities of being recognized by ClpXdN. The degradation assays were performed under the following conditions: 5 μg substrate, 1.3 μM ClpXdN, 1.3 μM ClpP, 150 mM NaCl, 1xATP regeneration system, 1 mM ATP, 2 mM DTT. The mixture was incubated at 37°C for 3 hours. The reaction was stopped by adding a 1x loading dye buffer and heated at 95 °C for 10 minutes. As a control group (t=0h), loading dye and heating step was immediately implemented after the assays were prepared. Both samples (t=0h and 3h) were loaded and checked by SDS-PAGE.

### Electrophysiology set-up and membrane formation

Recordings in planar lipid bilayers were conducted using a custom-designed chamber comprising two compartments, separated by a 25 μm thick Teflon membrane with an aperture of approximately 100 μm, as previously described [^17^]. A droplet containing n-hexadecane dissolved in n-pentane (4% v/v) was applied to the Teflon membrane, with the volume being approximately half that contained in a 10 μL glass capillary. After the evaporation of n-pentane, 450 μL of buffer was added to each compartment, followed by two droplets of a DPhPC:PDB_22_PEO_14_ solution (molar ratio 3:1) prepared by mixing separate 10 mg/mL solutions of DPhPC and PDB_22_PEO_14_ in n-pentane for MspA nanopore experiments. For CytK-4D experiments, a DPhPC:PDB_11_PEO_8_ solution (molar ratio 1:1) was prepared by mixing separate 10 mg/mL solutions of DPhPC and PDB_11_PEO_8_ in n-pentane[^18^]. Ag/AgCl electrodes were connected to the two compartments via agarose bridges (2.5% agarose, 3 M KCl solution), with the cis compartment grounded. Measurements were recorded using an Axon™ Digidata® 1550B digitizer and an Axopatch 200B amplifier (Molecular Devices), with data acquisition performed using Clampex 11.1 software.

### Enzymatic mediated protein translocation measurements

After isolating a single pore in the measurement buffer (1 M potassium gluconate, 50 mM HEPES, 10 mM MgCl_2_, pH 7.4), an I/V curve was performed to determine the orientation of the pore. Subsequently, different components were added to the chamber: the substrate (protein of interest) at a final concentration of 5 nM, the enzyme ClpX at a final concentration of 100 nM, ATP at 2 mM and the regeneration system containing a final concentration of 1.6 mM creatine phosphate, 0.4 µM creatine kinase, 1 mM DTT, and 0.5 mM EDTA. All experiments were conducted at 37 °C, with temperature controlled by a Peltier heating system.

### Data acquisition and analysis

All traces were collected using a sampling frequency of 10 kHz and a Bessel low-pass filter set to 2 kHz. The traces were first visualized using Clampex 11.1 software, and individual events (squiggles) were manually segmented. A consistent current signal associated with the specific sequence “GGGGS,” present between the SQ tag and the protein of interest (POI), was used to identify and segment the SQtag from the POI. Partial squiggles or squiggles containing interruption in the current levels (Figure S15) were excluded from the analysis. Each segmented squiggle was then analyzed using a custom Python script.

#### Preprocessing data

Each trace was first filtered by applying a low-pass Bessel filter of 50 Hz (N = 5, Wn = 0.01) using the function from signal processing scipy.signal Python package. The signal is then transformed into a step-like signal by identifying the change points with the Linearly Penalized Segmentation (PELT) function (model= l2, min_size=80, pen = 1) (additional text S6 and figure S16 and figure S17)[^19^]. Next, the average current value is calculated between two adjacent discontinuity points, generating a segmented signal with the average current between the identified points. Finally, the signal is normalized to the current values using Z-score normalization (transformed current) to highlight the signal variations from the signal’s average (Figure S17).

#### Alignment

To generate a reference signal for each specific protein, all the traces from one protein are aligned using the Dynamic Time Warping (DTW) function [^20^]. The pairwise distance between each trace is calculated, and the medoid is found, which is then used as the reference to align all the traces. Once all the traces are aligned to the medoid, the average and standard deviation of the current are calculated.

#### Protein and single-amino-acid substitution identification

For protein identification all the traces (Figure S18-23 for raw squiggles) belonging to the measured proteins are preprocessed, and the DTW distance between them is calculated. The resulting DTW distance matrix is used to create a hierarchical clustering heatmap and a t-distributed stochastic neighbor embedding (t-SNE) plot[^21^] .For single amino acid substitution identification, a specific DTW alignment between the reference signal of the wild-type protein and the mutated version is performed, followed by a point-by-point t-test to identify where the signal shows statistically significant differences (Figure S24-35, raw squiggles).

For PTMs, traces were pretreated to reduce the sampling space by pooling one out of every ten data points, reducing the sampling rate from 10 kHz to 1 kHz. Spike-shaped noise and blockades were removed using a median filter (*scipy*.*signal, medfilt*, kernel size = 3), followed by sliding-window spike detection. Spikes were identified when the sliding-window median differed by more than three standard deviations from adjacent windows. Identified spikes were replaced using a median value and Gaussian noise estimated from neighboring windows. The signal was then denoised using total variation denoising (*condat-tv*, TVDCondat2013, strength = 100) and segmented with the PELT algorithm (model = *l2*, penalty = log(len(signal))). Traces from the same substrate were averaged using shapeDWA (DTW window = 10) [^22^], after discarding the first 30 segments corresponding to the noisy leader domain. The number of segments in the averaged trace was set to the mean number across individual traces. Equal segment numbers in unmodified and modified substrates enabled elastic functional principal component analysis (*fdasrsf*). Average traces were warped for segment-by-segment comparison, and Cohen’s distance was calculated between corresponding segments; values with an absolute magnitude greater than three were considered significant.

