## Supplementary information for "Single-molecule identification of full-length proteins with single-amino-acid resolution using nanopores"

### Additional Text.

S1. Optimization of nanopore signals

The enzymatic activity of ClpX decreased with increasing KCl concentration (Figures S1, c). Since cation-selective nanopores require high [K+] for optimal ionic signals, we assessed the activity of ClpX in the presence of various potassium salts, including potassium acetate, potassium gluconate, potassium trifluoromethanesulfonate(KTF), $KNO_3$, and $K_2SO_4$. Our findings indicate that ClpX remains active in potassium acetate and potassium gluconate across a concentration range from 0.3 M to 1 M (Figure S1 d,e), whereas no activity was observed with KTF, $KNO_3$, or $K_2SO_4$ (Figure S1b). Potassium gluconate was identified as a salt that produced higher enzymatic activity at a concentration of 1M, and was therefore selected for the electrophysiology experiments. Compared to 1M KCl, the presence of potassium gluconate resulted in a similar I-V profile, with a decrease in total ionic current of approximately 21% at -65 mV and 37 $^{o}$C (figure S2 a), the conditions at which most of the experiments were conducted.

We studied the capture of the substrate (MBP-SQtag) at different voltages and observed almost no capture at -35 mV, with an increasing capture rate as the voltage increased from -50 mV to -65 mV and up to -80 mV (Figure S3, a,b ,c and d). We also investigated how the bias affected enzyme-mediated translocation, identifying an optimal squiggle frequency at -65 mV. Notably, low squiggle frequencies were observed at both -50 mV and -80 mV (Figure S3 e,f and g). This behavior can be attributed to two factors: at low voltage, capture is infrequent, while at high voltage, the nanopore is predominantly occupied by free substrate, possibly because the enzyme did not withstand the force induced by the applied potential. Additionally, at -80 mV across all proteins tested, we observed that many squiggles exhibited interruptions of the signal (the ionic current was nearly zero). This could be due to several factors. For example, at higher bias larger forces might be generated by the applied electric field (EF) and electroosmotic flow (EOF) may pull on the substrate inside the nanopore, potentially blocking the enzyme unfolding process.

S2. The effect of stalling domain on substrate unfolding by ClpX

To confirm that the presence of stalling domains (SD) slows down the unfolding of protein of interest (POI) by ClpX in solution, we constructed fusion proteins mNG-MBP containing different amounts of stalling domains, resulting in mNG-MBP, mNG-SD-MBP, mNG-SD-MBP-SQ. In the last construct, SQtag was introduced because it was placed in the C-terminus (therefore a ssrA tag and electrophoretic tag are required) and it also includes a stalling domain. As shown in Figure S9c, the fluorescence signal of each substrate alone remains stable over time, while the fluorescence signal of each substrate mixed with ClpX drops upon incubation. The unfolding rate by ClpX was negatively correlated with the number of stalling domains: substrates containing more stalling domains exhibited progressively reduced unfolding speeds. As expected, mNG–MBP was unfolded at the fastest rate, followed by mNG–SD–MBP, whereas mNG–SD–MBP–SQ was unfolded at the slowest rate. In summary, these results demonstrate that introducing SDs into the POI slows its unfolding by ClpX in solution.

S3. Protein translocation mediated by ClpX

Each recorded squiggle exhibits a stepwise translocation pattern, strongly suggesting that protein movement through the nanopore is regulated by enzymatic activity. To further confirm this is an enzyme-mediated translocation, rather than a passive translocation where the enzyme acts merely as a physical barrier, we performed control experiments in which enzyme kinetics were modulated by adding a slow-hydrolyzable ATP analog. Specifically, ATP-γS was added at concentrations of 1 or 2 mM in the presence of 2 mM ATP. Translocation times of SQ–MBP were measured with ClpX at –65 mV under these conditions. The mean translocation time increased from 8.2 ± 1.2 s (N = 9) in the presence of ATP alone to 17.1 ± 3.3 s (N=14) with 1 mM ATP-γS and to 27.5 ± 5.2 s (N=8) with 2 mM ATP-γS (Example of trace for each condition was illustrated in Figure S4). The increase in translocation time demonstrates that translocation is actively controlled by the enzyme rather than arising from passive EOF-driven transport.

S4. Amino acid substitution

We investigated the nanopore's ability to discriminate against amino acid substitutions in a polypeptide's unstructured region. Initially, we used the same sequence used by Montone et al.[[1]], which might allow comparison between the signal of MspA and CsgG nanopores. The sequence consists of repeats of glycine (G), serine (S), glutamic acid (E), and aspartic acid (D) flanked by two tyrosines (Y). This unstructured sequence was rich in negatively charged residues, which under our conditions required translocation against an applied potential. The Motone's sequence was slightly modified for our study by incorporating an additional tyrosine, creating a three-tyrosine (YYY) repeat. A double mutation was introduced at the center of the sequence, generating two variants: WW-GG and RR-DD. These sequences were inserted between the SQtag and the MBP protein domain (MBP-SAA(WW,GG)-SQtag and MBP-SAA(RR,DD)-SQtag).

The resulting squiggle traces (Figure S9 a,b, S20) show that the unstructured polypeptide region exhibits higher current relative to the SQtag and MBP domains. This is most likely due to the high content of neutral and small amino acids (G, S) and negatively charged residues (E, D), which are expected to allow a high amount of $K^+$ ions across ion selective MspA nanopores. Additionally, the figure demonstrates how the YYY repeats produce a specific and reproducible blocking signal in three distinct regions of the polypeptide. Furthermore, the high content of negatively charged residues might stretch the polypeptides under the negatively applied potentials used. Between the repeated regions, introduction of a GG mutation increased the current, while WW induced a higher current blockage. DD substitutions cause minimal variation, and the RR mutation results in a substantial current blockage.

Building on these findings, we next introduced mutations directly in the MBP protein region, selecting G and R, which exhibited the highest variability in the unstructured polypeptide region. We generated two MBP mutations with GGG and RRR amino acid substitutions, resulting in MBP-GGG-SQtag (Raw squiggles Figure S21) and MBP-RRR-SQtag (Raw squiggles Figure S22), respectively. Figure S9c clearly demonstrates variant signals, with the GGG mutation resulting in an increase in the current signal, while the RRR mutation caused a significant current blockage (Figure S9d). These results suggested that larger or positively charged side chains induce larger current blockades.

S5. Amino acid reading window

The number of amino acids addressed by this method starts at the C-terminus of the protein of interest (POI) and terminates when the enzymes release the last amino acid at the N-terminus of the POI. Since there is a distance between the *cis* entry of the nanopore and the recognition region near the *trans* entry, a portion of the POI is not enzymatically ratcheted through the nanopore and rapidly translocates in the nanopore, resulting in the loss of the characteristic stepwise motion observed during enzyme-mediated translocation.

The combined length of the nanopore and enzyme is approximately 15 nm, corresponding to ~50 amino acids [[2]], that could not be resolved by the system. Each POI analyzed in this study (Table S2) has been extended at the N-terminal with 21 amino acids containing a polypeptide linker and a histidine tag to aid protein purification. Considering MBP amino acid substitution, we observed substitution with arginine at position 27 (MBP-F27R) induced a strong reduction of the current signal at the very end of the squiggle, indicating that ~ 50 amino acids are not enzymatically threaded across the nanopore.

S6. Step size analysis and change-point detection

To convert raw current traces into step-like signals, we applied the Linearly Penalized Segmentation (PELT) algorithm with the L2 model, which detects changes in the mean of the data, setting the penalty to 1 to achieve maximum sensitivity.
The *min_size* parameter, which represents the minimum time interval required to define a step, was seated as 80 ms and selected based on values reported in the literature on ClpX kinetic steps and adapted to our system.
According to two different studies, the translocation rate expressed in AA/step (the number of amino acids processed by ClpX per ATP hydrolysis cycle) was reported to be 2 [1] and 6 [3]. Considering these findings, and given that in our system we measured an average translocation velocity of approximately 39 AA/s across the six proteins analyzed, the estimated duration of a single translocation step is about 51 ms in the case of 2 AA/step and about 154 ms in the case of 6 AA/step.
Testing these values in the PELT algorithm, a *min_size* of 51 ms led to overfitting, whereas 154 ms caused underfitting. Values of 80–100 ms provided the best agreement with the observed traces, and 80 ms was ultimately chosen (Figure Sx). Importantly, the number of detected steps remained robust across increased noise levels (50 Hz to 5 kHz), indicating that this parameter reliably identifies enzyme-mediated protein translocation events in our setup.

Table S1. Plasmids used in this study for protein production.

| Plasmids Name | Description | Reference |
|---|---|---|
| pT7sc1-MspA | pT7sc1 vector containing *MspA* gene with a Strep-Tag; $Amp^R$ | This work |
| pBAD-ClpXdN | pBAD vector containing *ClpXdN* (residue 62-424) gene with a Strep-Tag; $Amp^R$ | This work |
| pACYC-ClpP | pACYC vector containing *ClpP* gene with a Strep-Tag; $Cm^R$ | This work |
| pBAD-mNG | pBAD vector containing *mNG* gene with His-Tag and ssrA Tag; $Amp^R$ | This work |
| pT7sc1-MBP-10R | pT7sc1 vector containing *MBP-10R* gene with a His-Tag; $Amp^R$ | This work |
| pT7sc1-MBP-SQtag | pT7sc1 vector containing *MBP* gene, stalling domain consisting of an engineered leucine-zipper, slippery region, Strep Tag, electrophoretic tag, and a C-terminal ssrA Tag; $Amp^R$ | This work |
| pT7sc1-GBP-SQtag | pT7sc1 vector containing *GBP* gene, stalling domain consisting of an engineered leucine-zipper, slippery region, Strep Tag, and electrophoretic tag, and a C-terminal ssrA Tag; $Amp^R$ | This work |
| pT7sc1-LBP-SQtag | pT7sc1 vector containing *LBP* gene, stalling domain consisting of an engineered leucine-zipper, slippery region, Strep Tag, and electrophoretic tag, and a C-terminal ssrA Tag; $Amp^R$ | This work |
| pT7sc1-SpuD-SQtag | pT7sc1 vector containing *LBP* gene, stalling domain consisting of an engineered leucine-zipper, slippery region, Strep Tag, and electrophoretic tag, and a C-terminal ssrA Tag; $Amp^R$ | This work |
| pT7sc1-HitA-SQtag | pT7sc1 vector containing *HitA* gene, stalling domain consisting of an engineered leucine-zipper, slippery region, Strep Tag, and electrophoretic tag, and a C-terminal ssrA Tag; $Amp^R$ | This work |
| pT7sc1-MNG-SQtag | pT7sc1 vector containing *mNG* gene, stalling domain consisting of an engineered leucine-zipper, slippery region, Strep Tag, and electrophoretic tag, and a C-terminal ssrA Tag; $Amp^R$ | This work |
| pT7sc1-MNG-SD_MBP-SQtag | pT7sc1 vector containing *mNG_MBP* gene, stalling domain consisting of an engineered leucine-zipper, slippery region, Strep Tag, and electrophoretic tag, and a C-terminal ssrA Tag; $Amp^R$ | This work |
| pT7sc1-MBP-GGG-SQtag | pT7sc1 vector containing *MBP* gene with GGG substitution (240-242), stalling domain consisting of an engineered leucine-zipper, slippery region, Strep Tag, electrophoretic tag, and a C-terminal ssrA Tag; $Amp^R$ | This work |
| pT7sc1-MBP-RRR-SQtag | pT7sc1 vector containing *MBP* gene with RRR substitution (307-309), stalling domain consisting of an engineered leucine-zipper, slippery region, Strep Tag, electrophoretic tag, and a C-terminal ssrA Tag; $Amp^R$ | This work |
| pT7sc1-MBP-SAA (WW,GG)-SQtag | pT7sc1 vector containing *MBP* gene, SAA (WW,GG) fragment, stalling domain consisting of an engineered leucine-zipper, slippery region, Strep Tag, electrophoretic tag, and a C-terminal ssrA Tag; $Amp^R$ | This work |

| | | |
|---|---|---|
| pT7sc1-MBP-SAA (RR,DD)-SQtag | pT7sc1 vector containing *MBP* gene, SAA (RR,DD) fragment, stalling domain consisting of an engineered leucine-zipper, slippery region, Strep Tag, electrophoretic tag, and a C-terminal ssrA Tag; $Amp^R$ | This work |
| pT7sc1-MBP F27R-SQtag | pT7sc1 vector containing *MBP F27R* gene, stalling domain consisting of an engineered leucine-zipper, slippery region, Strep Tag, electrophoretic tag, and a C-terminal ssrA Tag; $Amp^R$ | This work |
| pT7sc1-MBP R98D-SQtag | pT7sc1 vector containing *MBP R98D* gene, stalling domain consisting of an engineered leucine-zipper, slippery region, Strep Tag, electrophoretic tag, and a C-terminal ssrA Tag; $Amp^R$ | This work |
| pT7sc1-MBP K142G-SQtag | pT7sc1 vector containing *MBP K142G* gene, stalling domain consisting of an engineered leucine-zipper, slippery region, Strep Tag, electrophoretic tag, and a C-terminal ssrA Tag; $Amp^R$ | This work |
| pT7sc1-MBP K142E-SQtag | pT7sc1 vector containing *MBP K142E* gene, stalling domain consisting of an engineered leucine-zipper, slippery region, Strep Tag, electrophoretic tag, and a C-terminal ssrA Tag; $Amp^R$ | This work |
| pT7sc1-MBP K142F-SQtag | pT7sc1 vector containing *MBP K142F* gene, stalling domain consisting of an engineered leucine-zipper, slippery region, Strep Tag, electrophoretic tag, and a C-terminal ssrA Tag; $Amp^R$ | This work |
| pT7sc1-MBP N201R-SQtag | pT7sc1 vector containing *MBP N201R* gene, stalling domain consisting of an engineered leucine-zipper, slippery region, Strep Tag, electrophoretic tag, and a C-terminal ssrA Tag; $Amp^R$ | This work |
| pT7sc1-MBP Y307R-SQtag | pT7sc1 vector containing *MBP Y307R* gene, stalling domain consisting of an engineered leucine-zipper, slippery region, Strep Tag, electrophoretic tag, and a C-terminal ssrA Tag; $Amp^R$ | This work |
| pT7sc1-LBP D268R-SQtag | pT7sc1 vector containing *LBP D268R* gene, stalling domain consisting of an engineered leucine-zipper, slippery region, Strep Tag, and electrophoretic tag, and a C-terminal ssrA Tag; $Amp^R$ | This work |
| pBAD-MBP-K163TAG | pBAD vector containing *MBP-K163TAG* gene with a N-terminal His6-Tag; $Amp^R$ | This work |
| pEVOL_AcKRS3 | pEVOL vector containing two copies of a *M. barkeri* PylRS mutant known to incorporate AcK under a constitutive GlnS promoter and an arabinose inducible promoter with a pyrrolysyl-tRNA (PylT) copy under a constitutive promoter, $Cm^R$. | This work |

SQtag: sequencing tag.

Table S2. The complete protein sequence of the substrate constructs in this study. SD: stalling domain. SQtag: sequencing tag.

| |
|---|
| **MBP-10R** |
| MGSSHHHHHHSSGLVPRGSHNKIEEGKLVIWINGDKGYNGLAEVGKKFEKDTGIKVTVEHPDKLEEKFPQVAAT<br>GDGPDIIFWAHDRFGGYAQSGLLAEITPDKAFQDKLYPFTWDAVRYNGKLIAYPIAVEALSLIYNKDLLPNPPKTW<br>EEIPALDKELKAKGKSALMFNLQEPYFTWPLIAADGGYAFKYENGKYDIKDVGVDNAGAKAGLTFLVDLIKNKHM<br>NADTDYSIAEAAFNKGETAMTINGPWAWSNIDTSKVNYGVTVLPTFKGQPSKPFVGVLSAGINAASPNKELAKE<br>FLENYLLTDEGLEAVNKDKPLGAVALKSYEEELAKDPRIAATMENAQKGEIMPNIPQMSAFWYAVRTAVINAAS<br>GRQTVDEALKDAQTNSSS**RRRRRRRRRR**LGIEGLYFQSHSSAANDENYALAA* |
| **MBP**-SQtag |
| MGSSHHHHHHSSGLVPRGSHNKIEEGKLVIWINGDKGYNGLAEVGKKFEKDTGIKVTVEHPDKLEEKFPQVAAT<br>GDGPDIIFWAHDRFGGYAQSGLLAEITPDKAFQDKLYPFTWDAVRYNGKLIAYPIAVEALSLIYNKDLLPNPPKTW<br>EEIPALDKELKAKGKSALMFNLQEPYFTWPLIAADGGYAFKYENGKYDIKDVGVDNAGAKAGLTFLVDLIKNKHM<br>NADTDYSIAEAAFNKGETAMTINGPWAWSNIDTSKVNYGVTVLPTFKGQPSKPFVGVLSAGINAASPNKELAKE<br>FLENYLLTDEGLEAVNKDKPLGAVALKSYEEELAKDPRIAATMENAQKGEIMPNIPQMSAFWYAVRTAVINAAS<br>GRQTVDEALKDAQTNGGGGSLEIRAAALRRRNTALRTRVAELRQRVQRLRNEVSQYETRYGPLGSSGGGGSLPG<br>YRTRYQSVINELRQVEQELEAVETELATNEQELAAAEIELSSSKKKKKKKKKKLGIEGLYFQSHWSHPQFEKGGRRR<br>RRRRRRRSSAANDENYALAA* |
| **SpuD**-SQtag |
| MGSSHHHHHHSSGLVPRGSHNADNKVLHVYNWSDYIAPDTLEKFTKETGIKVVYDVYDSNEVLEAKLLAGKSGY<br>DVVVPSNSFLAKQIKAGVYQKLDKSKLPNWKNLNKDLMHTLEVSDPGNEHAIPYMWGTIGIGYNPDKVKAAFG<br>DNAPVDSWDLVFKPENIQKLKQCGVSFLDSPTEILPAALHYLGYKPDTDNPKELKAAEELFLKIRPYVTYFHSSKYIS<br>DLANGNICVAIGYSGDIYQAKSRAEEAKNKVTVKYNIPKEGAGSFFDMVAIPKDAENTEGALAFVNFLMKPEIMA<br>EITDVVQFPNGNAAATPLVSEAIRNDPGIYPSEEVMKKLYTFPDLPAKTQRAMTRSWTKIKSGKNGGGGSLEIRA<br>AALRRRNTALRTRVAELRQRVQRLRNEVSQYETRYGPLGSSGGGGSLPGYRTRYQSVINELRQVEQELEAVETEL<br>ATNEQELAAAEIELSSSKKKKKKKKKKLGIEGLYFQSHWSHPQFEKGGRRRRRRRRRRSSAANDENYALAA* |
| **LBP**-SQtag |
| MGSSHHHHHHSSGLVPRGSHNDDIKVAVVGAMSGPIAQWGDMEFNGARQAIKDINAKGGIKGDKLVGVEYD<br>DACDPKQAVAVANKIVNDGIKYVIGHLCSSSTQPASDIYEDEGILMISPGATNPELTQRGYQHIMRTAGLDSSQG<br>PTAAKYILETVKPQRIAIIHDKQQYGEGLARSVQDGLKAANANVVFFDGITAGEKDFSALIARLKKENIDFVYYGGY<br>YPEMGQMLRQARSVGLKTQFMGPEGVGNASLSNIAGDAAEGMLVTMPKRYDQDPANQGIVDALKADKKDPS<br>GPYVWITYAAVQSLATALERTGSDEPLALVKDLKANGANTVIGPLNWDEKGDLKGFDFGVFQWHADGSSTAAK<br>NGGGGSLEIRAAALRRRNTALRTRVAELRQRVQRLRNEVSQYETRYGPLGSSGGGGSLPGYRTRYQSVINELRQV<br>EQELEAVETELATNEQELAAAEIELSSSKKKKKKKKKKLGIEGLYFQSHWSHPQFEKGGRRRRRRRRRRSSAANDE<br>NYALAA* |
| **HitA**-SQtag |
| MGSSHHHHHHSSGLVPRGSHNAALEVLFQGPGYQDPNSMDPVTLTLYNGQHAATGIAIAKAFQDKTGIQVKIR<br>KGGDGQLASQITEEGARSPADVLYTEESPPLIRLASAGLLAKLEPETLALVEPEHAGGNGDWIGITARTRVLAYNPK<br>KIDEKDLPKSLMDLSDPSWSGRFGFVPTSGAFLEQVAAVIKLKGQEEAEDWLTGLKAFGSIYTNNVTAMKAVEN<br>GEVDMALINNYYWYTLKKEKGELNSRLHYFGNQDPGALVTVSGAAVLKSSKHPREAQQFVAFMLSEEGQKAILS<br>QSAEYPMRKGMQADPALKPFAELDPPKLTPADLGEASEALSLERDVGLNNGGGGSLEIRAAALRRRNTALRTRV<br>AELRQRVQRLRNEVSQYETRYGPLGSSGGGGSLPGYRTRYQSVINELRQVEQELEAVETELATNEQELAAAEIELS<br>SSKKKKKKKKKKLGIEGLYFQSHWSHPQFEKGGRRRRRRRRRRSSAANDENYALAA* |
| **TbpA**-SQtag |
| MGSSHHHHHHSSGLVPRGSHNLVPRGSHMKPVLTVYTYDSFAADWGPGPVVKKAFEADCNCELKLVALEDGV<br>SLLNRLRMEGKNSKADVVLGLDNNLLDAASKTGLFAKSGVAADAVNVPGGWNNDTFVPFDYGYFAFVYDKNKL<br>KNPPQSLKELVESDQNWRVIYQDPRTSTPGLGLLLWMQKVYGDDAPQAWQKLAKKTVTVTKGWSEAYGLFLK<br>GESDLVLSYTTSPAYHILEEKKDNYAAANFSEGHYLQVEVAARTAASKQPELAQKFLQFMVSPAFQNAIPTGNW<br>MYPVANVTLPAGFEKLTKPATTLEFTPAEVAAQRQAWISEWQRAVSRNGGGGSLEIRAAALRRRNTALRTRVAE<br>LRQRVQRLRNEVSQYETRYGPLGSSGGGGSLPGYRTRYQSVINELRQVEQELEAVETELATNEQELAAAEIELSSS<br>KKKKKKKKKKLGIEGLYFQSHWSHPQFEKGGRRRRRRRRRRSSAANDENYALAA* |
| **GBP**-SQtag |
| MGSSHHHHHHSSGLVPRGSHNADTRIGVTIYKYDDNFMSVVRKAIEQDAKAAPDVQLLMNDSQNDQSKQND |

| |
|---|
| QIDVLLAKGVKALAINLVDPAAAGTVIEKARGQNVPVVFFNKEPSRKALDSYDKAYYVGTDSKESGIIQGDLIAKH<br>WAANQGWDLNKDGQIQFVLLKGEPGHPDAEARTTYVIKELNDKGIKTEQLQLDTAMWDTAQAKDKMDAWL<br>SGPNANKIEVVIANNDAMAMGAVEALKAHNKSSIPVFGVDALPEALALVKSGALAGTVLNDANNQAKATFDLA<br>KNLADGKGAADGTNWKIDNKVVRVPYVGVDKDNLAEFSKKNGGGGSLEIRAAALRRRNTALRTRVAELRQRVQ<br>RLRNEVSQYETRYGPLGSSGGGGSLPGYRTRYQSVINELRQVEQELEAVETELATNEQELAAAEIELSSSKKKKKKK<br>KKKLGIEGLYFQSHWSHPQFEKGGRRRRRRRRRRSSAANDENYALAA* |
| **MNG**-SQtag |
| MGSSHHHHHHSSGLVPRGSHNVSKGEEDNMASLPATHELHIFGSINGVDFDMVGQGTGNPNDGYEELNLKST<br>KGDLQFSPWILVPHIGYGFHQYLPYPDGMSPFQAAMVDGSGYQVHRTMQFEDGASLTVNYRYTYEGSHIKGEA<br>QVKGTGFPADGPVMTNSLTAADWCRSKKTYPNDKTIISTFKWSYTTGNGKRYRSTARTTYTFAKPMAANYLKN<br>QPMYVFRKTELKHSKTELNFKEWQKAFTDVMGMDELYKNGGGGSLEIRAAALRRRNTALRTRVAELRQRVQRL<br>RNEVSQYETRYGPLGSSGGGGSLPGYRTRYQSVINELRQVEQELEAVETELATNEQELAAAEIELSSSKKKKKKKKK<br>KLGIEGLYFQSHGGWSHPQFEKGGRRRRRRRRRRSSAANDENYALAA* |
| **MNG**-**MBP** |
| MGSSHHHHHHSSGLVPRGSHNVSKGEEDNMASLPATHELHIFGSINGVDFDMVGQGTGNPNDGYEELNLKST<br>KGDLQFSPWILVPHIGYGFHQYLPYPDGMSPFQAAMVDGSGYQVHRTMQFEDGASLTVNYRYTYEGSHIKGEA<br>QVKGTGFPADGPVMTNSLTAADWCRSKKTYPNDKTIISTFKWSYTTGNGKRYRSTARTTYTFAKPMAANYLKN<br>QPMYVFRKTELKHSKTELNFKEWQKAFTDVMGMDELYKNGSHGGKIEEGKLVIWINGDKGYNGLAEVGKKFEK<br>DTGIKVTVEHPDKLEEKFPQVAATGDGPDIIFWAHDRFGGYAQSGLLAEITPDKAFQDKLYPFTWDAVRYNGKLI<br>AYPIAVEALSLIYNKDLLPNPPKTWEEIPALDKELKAKGKSALMFNLQEPYFTWPLIAADGGYAFKYENGKYDIKDV<br>GVDNAGAKAGLTFLVDLIKNKHMNADTDYSIAEAAFNKGETAMTINGPWAWSNIDTSKVNYGVTVLPTFKGQP<br>SKPFVGVLSAGINAASPNKELAKEFLENYLLTDEGLEAVNKDKPLGAVALKSYEEELAKDPRIAATMENAQKGEIM<br>PNIPQMSAFWYAVRTAVINAASGRQTVDEALKDAQTNGGSSAANDENYALAA* |
| **MNG**-SD-**MBP** |
| MGSSHHHHHHSSGLVPRGSHNVSKGEEDNMASLPATHELHIFGSINGVDFDMVGQGTGNPNDGYEELNLKST<br>KGDLQFSPWILVPHIGYGFHQYLPYPDGMSPFQAAMVDGSGYQVHRTMQFEDGASLTVNYRYTYEGSHIKGEA<br>QVKGTGFPADGPVMTNSLTAADWCRSKKTYPNDKTIISTFKWSYTTGNGKRYRSTARTTYTFAKPMAANYLKN<br>QPMYVFRKTELKHSKTELNFKEWQKAFTDVMGMDELYKNGGGGSLEIRAAALRRRNTALRTRVAELRQRVQRL<br>RNEVSQYETRYGPLGSSGGGGSLPGYRTRYQSVINELRQVEQELEAVETELATNEQELAAAEIELSSSKKKKKKKKK<br>KLGIEGLYFQSHGGKIEEGKLVIWINGDKGYNGLAEVGKKFEKDTGIKVTVEHPDKLEEKFPQVAATGDGPDIIFW<br>AHDRFGGYAQSGLLAEITPDKAFQDKLYPFTWDAVRYNGKLIAYPIAVEALSLIYNKDLLPNPPKTWEEIPALDKEL<br>KAKGKSALMFNLQEPYFTWPLIAADGGYAFKYENGKYDIKDVGVDNAGAKAGLTFLVDLIKNKHMNADTDYSIA<br>EAAFNKGETAMTINGPWAWSNIDTSKVNYGVTVLPTFKGQPSKPFVGVLSAGINAASPNKELAKEFLENYLLTDE<br>GLEAVNKDKPLGAVALKSYEEELAKDPRIAATMENAQKGEIMPNIPQMSAFWYAVRTAVINAASGRQTVDEALK<br>DAQTNGGSSAANDENYALAA* |
| **MNG**-SD-**MBP**-SQtag |
| MGSSHHHHHHSSGLVPRGSHNVSKGEEDNMASLPATHELHIFGSINGVDFDMVGQGTGNPNDGYEELNLKST<br>KGDLQFSPWILVPHIGYGFHQYLPYPDGMSPFQAAMVDGSGYQVHRTMQFEDGASLTVNYRYTYEGSHIKGEA<br>QVKGTGFPADGPVMTNSLTAADWCRSKKTYPNDKTIISTFKWSYTTGNGKRYRSTARTTYTFAKPMAANYLKN<br>QPMYVFRKTELKHSKTELNFKEWQKAFTDVMGMDELYKNGGGGSLEIRAAALRRRNTALRTRVAELRQRVQRL<br>RNEVSQYETRYGPLGSSGGGGSLPGYRTRYQSVINELRQVEQELEAVETELATNEQELAAAEIELSSSKKKKKKKKK<br>KLGIEGLYFQSHGGKIEEGKLVIWINGDKGYNGLAEVGKKFEKDTGIKVTVEHPDKLEEKFPQVAATGDGPDIIFW<br>AHDRFGGYAQSGLLAEITPDKAFQDKLYPFTWDAVRYNGKLIAYPIAVEALSLIYNKDLLPNPPKTWEEIPALDKEL<br>KAKGKSALMFNLQEPYFTWPLIAADGGYAFKYENGKYDIKDVGVDNAGAKAGLTFLVDLIKNKHMNADTDYSIA<br>EAAFNKGETAMTINGPWAWSNIDTSKVNYGVTVLPTFKGQPSKPFVGVLSAGINAASPNKELAKEFLENYLLTDE<br>GLEAVNKDKPLGAVALKSYEEELAKDPRIAATMENAQKGEIMPNIPQMSAFWYAVRTAVINAASGRQTVDEALK<br>DAQTNGGGGSLEIRAAALRRRNTALRTRVAELRQRVQRLRNEVSQYETRYGPLGSSGGGGSLPGYRTRYQSVINE<br>LRQVEQELEAVETELATNEQELAAAEIELSSSKKKKKKKKKKLGIEGLYFQSHGGWSHPQFEKGGRRRRRRRRRRS<br>SAANDENYALAA* |
| **MBP-SAA(WW,GG)**-SQtag |
| MGSSHHHHHHSSGLVPRGSHNKIEEGKLVIWINGDKGYNGLAEVGKKFEKDTGIKVTVEHPDKLEEKFPQVAAT<br>GDGPDIIFWAHDRFGGYAQSGLLAEITPDKAFQDKLYPFTWDAVRYNGKLIAYPIAVEALSLIYNKDLLPNPPKTW<br>EEIPALDKELKAKGKSALMFNLQEPYFTWPLIAADGGYAFKYENGKYDIKDVGVDNAGAKAGLTFLVDLIKNKHM<br>NADTDYSIAEAAFNKGETAMTINGPWAWSNIDTSKVNYGVTVLPTFKGQPSKPFVGVLSAGINAASPNKELAKE<br>FLENYLLTDEGLEAVNKDKPLGAVALKSYEEELAKDPRIAATMENAQKGEIMPNIPQMSAFWYAVRTAVINAAS<br>GRQTVDEALKDAQTNGGGGSSGSGSSDSSGGYYYGGSSDSSGSGSSESGSESSGSGSSDSSGG**WW**GGSSDSSG |

| |
|---|
| SGSSESGSESSGSGSSDSSGGYYYGGSSDSSGSGSSESGSESSGSGSSDSSGG**GG**GGSSDSSGSGSSESGSESSGSG<br>SSDSSGGYYYGGSSDSSGSGSSESGSESSGSGSSDSSGGSGENLYFQSLEIRAAALRRRNTALRTRVAELRQRVQRL<br>RNEVSQYETRYGPLGSSGGGGSLPGYRTRYQSVINELRQVEQELEAVETELATNEQELAAAEIELSSSKKKKKKKKK<br>KLGIEGLYFQSHWSHPQFEKGGRRRRRRRRRRSSAANDENYALAA* |
| **MBP-SAA(RR,DD)**-SQtag |
| MGSSHHHHHHSSGLVPRGSHNKIEEGKLVIWINGDKGYNGLAEVGKKFEKDTGIKVTVEHPDKLEEKFPQVAAT<br>GDGPDIIFWAHDRFGGYAQSGLLAEITPDKAFQDKLYPFTWDAVRYNGKLIAYPIAVEALSLIYNKDLLPNPPKTW<br>EEIPALDKELKAKGKSALMFNLQEPYFTWPLIAADGGYAFKYENGKYDIKDVGVDNAGAKAGLTFLVDLIKNKHM<br>NADTDYSIAEAAFNKGETAMTINGPWAWSNIDTSKVNYGVTVLPTFKGQPSKPFVGVLSAGINAASPNKELAKE<br>FLENYLLTDEGLEAVNKDKPLGAVALKSYEEELAKDPRIAATMENAQKGEIMPNIPQMSAFWYAVRTAVINAAS<br>GRQTVDEALKDAQTNGGGGSSGSGSSDSSGGYYYGGSSDSSGSGSSESGSESSGSGSSDSSGG**RR**GGSSDSSGS<br>GSSESGSESSGSGSSDSSGGYYYGGSSDSSGSGSSESGSESSGSGSSDSSGG**DD**GGSSDSSGSGSSESGSESSGSGS<br>SDSSGGYYYGGSSDSSGSGSSESGSESSGSGSSDSSGGSGENLYFQSLEIRAAALRRRNTALRTRVAELRQRVQRL<br>RNEVSQYETRYGPLGSSGGGGSLPGYRTRYQSVINELRQVEQELEAVETELATNEQELAAAEIELSSSKKKKKKKKK<br>KLGIEGLYFQSHWSHPQFEKGGRRRRRRRRRRSSAANDENYALAA* |
| **MBP**-GGG-SQtag |
| MGSSHHHHHHSSGLVPRGSHNKIEEGKLVIWINGDKGYNGLAEVGKKFEKDTGIKVTVEHPDKLEEKFPQVAAT<br>GDGPDIIFWAHDRFGGYAQSGLLAEITPDKAFQDKLYPFTWDAVRYNGKLIAYPIAVEALSLIYNKDLLPNPPKTW<br>EEIPALDKELKAKGKSALMFNLQEPYFTWPLIAADGGYAFKYENGKYDIKDVGVDNAGAKAGLTFLVDLIKNKHM<br>NADTDYSIAEAAFNKGETAMTINGPWAWSNIDTSK**GGG**GVTVLPTFKGQPSKPFVGVLSAGINAASPNKELAKE<br>FLENYLLTDEGLEAVNKDKPLGAVALKSYEEELAKDPRIAATMENAQKGEIMPNIPQMSAFWYAVRTAVINAAS<br>GRQTVDEALKDAQTNGGGGSLEIRAAALRRRNTALRTRVAELRQRVQRLRNEVSQYETRYGPLGSSGGGGSLPG<br>YRTRYQSVINELRQVEQELEAVETELATNEQELAAAEIELSSSKKKKKKKKKKLGIEGLYFQSHWSHPQFEKGGRRR<br>RRRRRRRRSSAANDENYALAA* |
| **MBP**-RRR-SQtag |
| MGSSHHHHHHSSGLVPRGSHNKIEEGKLVIWINGDKGYNGLAEVGKKFEKDTGIKVTVEHPDKLEEKFPQVAAT<br>GDGPDIIFWAHDRFGGYAQSGLLAEITPDKAFQDKLYPFTWDAVRYNGKLIAYPIAVEALSLIYNKDLLPNPPKTW<br>EEIPALDKELKAKGKSALMFNLQEPYFTWPLIAADGGYAFKYENGKYDIKDVGVDNAGAKAGLTFLVDLIKNKHM<br>NADTDYSIAEAAFNKGETAMTINGPWAWSNIDTSKVNYGVTVLPTFKGQPSKPFVGVLSAGINAASPNKELAKE<br>FLENYLLTDEGLEAVNKDKPLGAVALKS**RRR**ELAKDPRIAATMENAQKGEIMPNIPQMSAFWYAVRTAVINAAS<br>GRQTVDEALKDAQTNGGGGSLEIRAAALRRRNTALRTRVAELRQRVQRLRNEVSQYETRYGPLGSSGGGGSLPG<br>YRTRYQSVINELRQVEQELEAVETELATNEQELAAAEIELSSSKKKKKKKKKKLGIEGLYFQSHWSHPQFEKGGRRR<br>RRRRRRRRSSAANDENYALAA* |
| MBP-**K163TAG** |
| MGSSHHHHHHSSGLVPRGSHNKIEEGKLVIWINGDKGYNGLAEVGKKFEKDTGIKVTVEHPDKLEEKFPQVAAT<br>GDGPDIIFWAHDRFGGYAQSGLLAEITPDKAFQDKLYPFTWDAVRYNGKLIAYPIAVEALSLIYNKDLLPNPPKTW<br>EEIPALDKELKA*GKSALMFNLQEPYFTWPLIAADGGYAFKYENGKYDIKDVGVDNAGAKAGLTFLVDLIKNKHM<br>NADTDYSIAEAAFNKGETAMTINGPWAWSNIDTSKVNYGVTVLPTFKGQPSKPFVGVLSAGINAASPNKELAKE<br>FLENYLLTDEGLEAVNKDKPLGAVALKSYEEELAKDPRIAATMENAQKGEIMPNIPQMSAFWYAVRTAVINAAS<br>GRQTVDEALKDAQTNGGGGSLEIRAAALRRRNTALRTRVAELRQRVQRLRNEVSQYETRYGPLGSSGGGGSLPG<br>YRTRYQSVINELRQVEQELEAVETELATNEQELAAAEIELSSSKKKKKKKKKKLGIEGLYFQSHWSHPQFEKGGRRR<br>RRRRRRRRSSAANDENYALAA* |
| **AcKRS3** |
| MDKKPLDVLISATGLWMSRTGTLHKIKHHEVSRSKIYIEMACGDHLVVNNSRSCRTARAFRHHKYRKTCKRCRVS<br>DEDINNFLTRSTESKNSVKVRVVSAPKVKKAMPKSVSRAPKPLENSVSAKASTNTSRSVPSPAKSTPNSSVPASAP<br>APSLTRSQLDRVEALLSPEDKISLNMAKPFRELEPELVTRRKNDFQRLYTNDREDYLGKLERDITKFFVDRGFLEIKS<br>PILIPAEYVERMGINNDTELSKQIFRVDKNLCLRPMMAPTIFNYARKLDRILPGPIKIFEVGPCYRKESDGKEHLEEF<br>TMVNFFQMGSGCTRENLEALIKEFLDYLEIDFEIVGDSCMVYGDTLDIMHGDLELSSAVVGPVSLDREWGIDKP<br>WIGAGFGLERLLKVMHGFKNIKRASRSESYYNGISTNL* |

Table S3. Analysis of the features extracted from protein translocation traces, including translocation duration and the number of observed steps. The calculation of the number of amino acids for each protein is based on the total number of amino acids starting from the C-terminus of the protein and non-considering the SQtag sequence, subtracting the last 50, which are not read by the nanopore. The averages are derived from a selected number of traces for each protein: MBP-SQtag (N=22), LBP-SQtag (N=19), GBP-SQtag (N=12), SpuD-SQtag (N=14), MNG-SQtag (N=16), and HitA-SQtag (N=14).

| Protein | Number of Amino acids read by MspA | Average time (s) ± STD | Average of Amino acids/seconds | Number of steps found in the trace ± STD | Average of the number of Amino acids for single step |
|---|---|---|---|---|---|
| MBP | 340 | 10 ± 2 | 34 | 88 ± 14 | 4 |
| LBP | 320 | 10 ± 3 | 32 | 82 ± 21 | 4 |
| GBP | 284 | 9.7 ± 2.5 | 29 | 81 ± 18 | 4 |
| SpuD | 318 | 7 ± 1 | 45 | 62 ± 7 | 5 |
| MNG | 210 | 3.7 ± 0.6 | 57 | 36 ± 5 | 6 |
| HitA | 301 | 7.5 ± 1.7 | 40 | 66 ± 16 | 5 |

### Supplementary figures

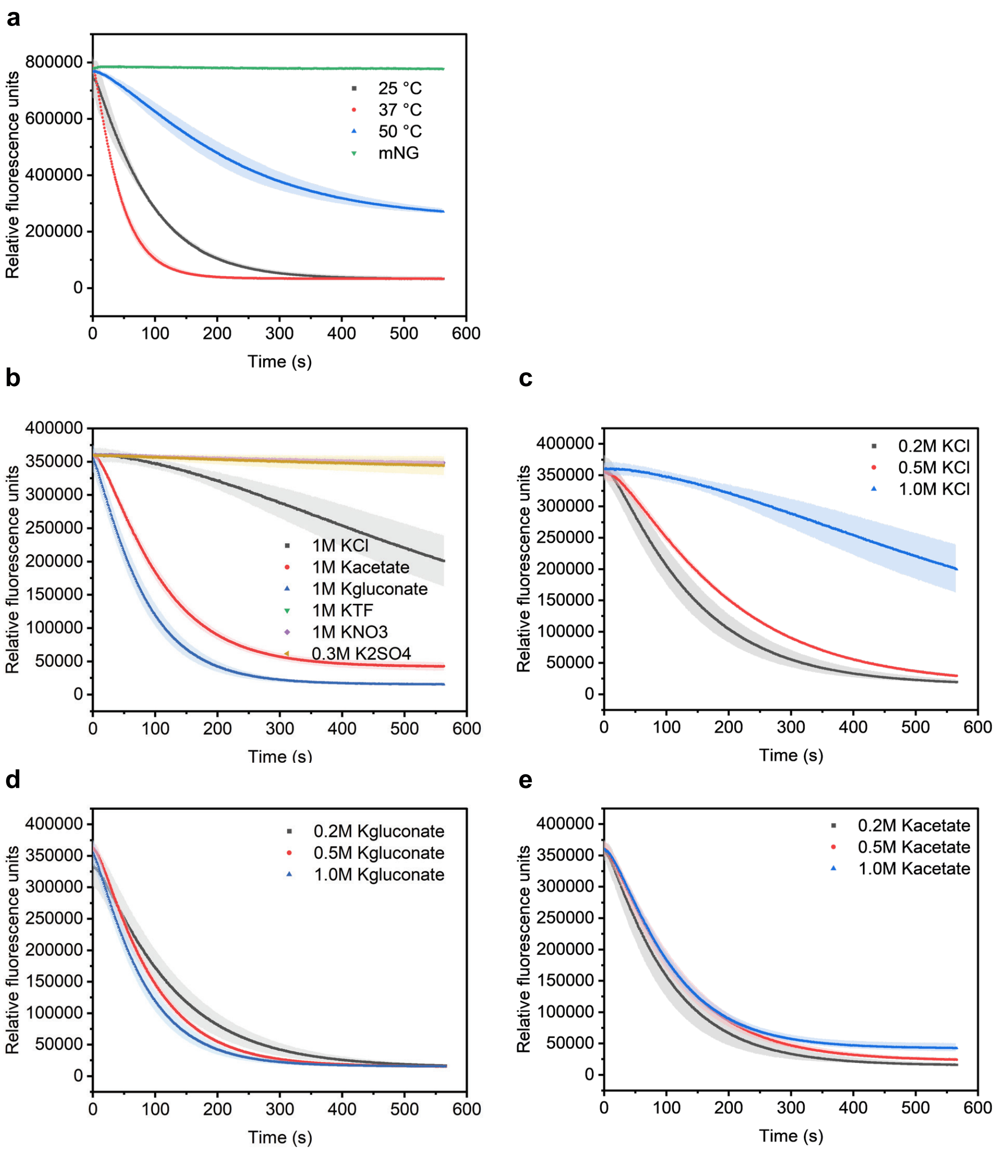


**Figure S1**. **Optimization of ClpX-ClpP enzyme activity under various conditions.** (a) Degradation assay performed at different temperatures. 37°C is the optimal condition used for subsequent experiments. Reactions were conducted in a 150 µL mixture containing 50 mM HEPES (pH 7.5), 5 mM $MgCl_2$, 200 mM KCl, 4 mM ATP, an ATP regeneration system, 1 mM DTT, 500 nM ClpX and ClpP, and 66 nM mNG-ssrA. (b) Assays conducted with various salts. ClpX-ClpP activity is diminished at 1 M KCl, 1 M $KNO_3$, 1 M KTF (potassium trifluoromethanesulfonate), and 0.3 M $K_2SO_4$, whereas higher activity is observed in the presence of 1 M K-gluconate or 1 M K-acetate. Reactions were conducted in a 150 µL mixture containing 50 mM HEPES (pH 7.5), 5 mM $MgCl_2$, corresponding potassium salts, 4 mM ATP, an ATP regeneration system, 1 mM DTT, 250 nM ClpX and ClpP, and 33 nM mNG-ssrA. Panels (c), (d), and (e) compare enzyme activity at different concentrations of KCl, K-gluconate, and K-acetate, respectively.

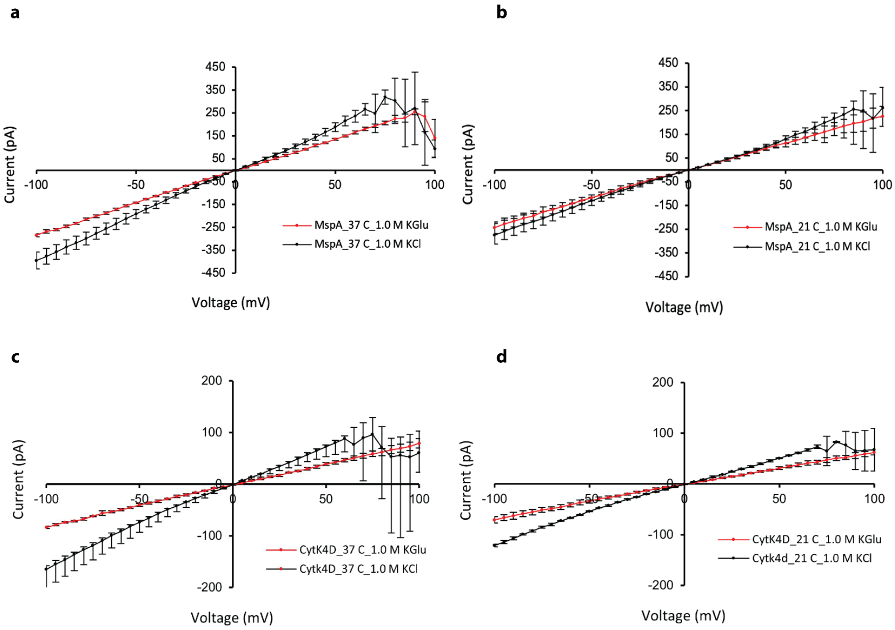


**Figure S2. Example I-V curves for MspA and CytK-4D nanopores in K-gluconate and 1 M KCl**. (a) Comparison of MspA (N=5) in 1 M K-gluconate (red line) and 1 M KCl (black line) at 37°C. (b) Comparison of CytK-4D (N=5) in 1 M K-gluconate (red line) and 1 M KCl (black line) at 21°C. Conditions: 50 mM HEPES, 10 mM $MgCl_2$, pH 7.4.

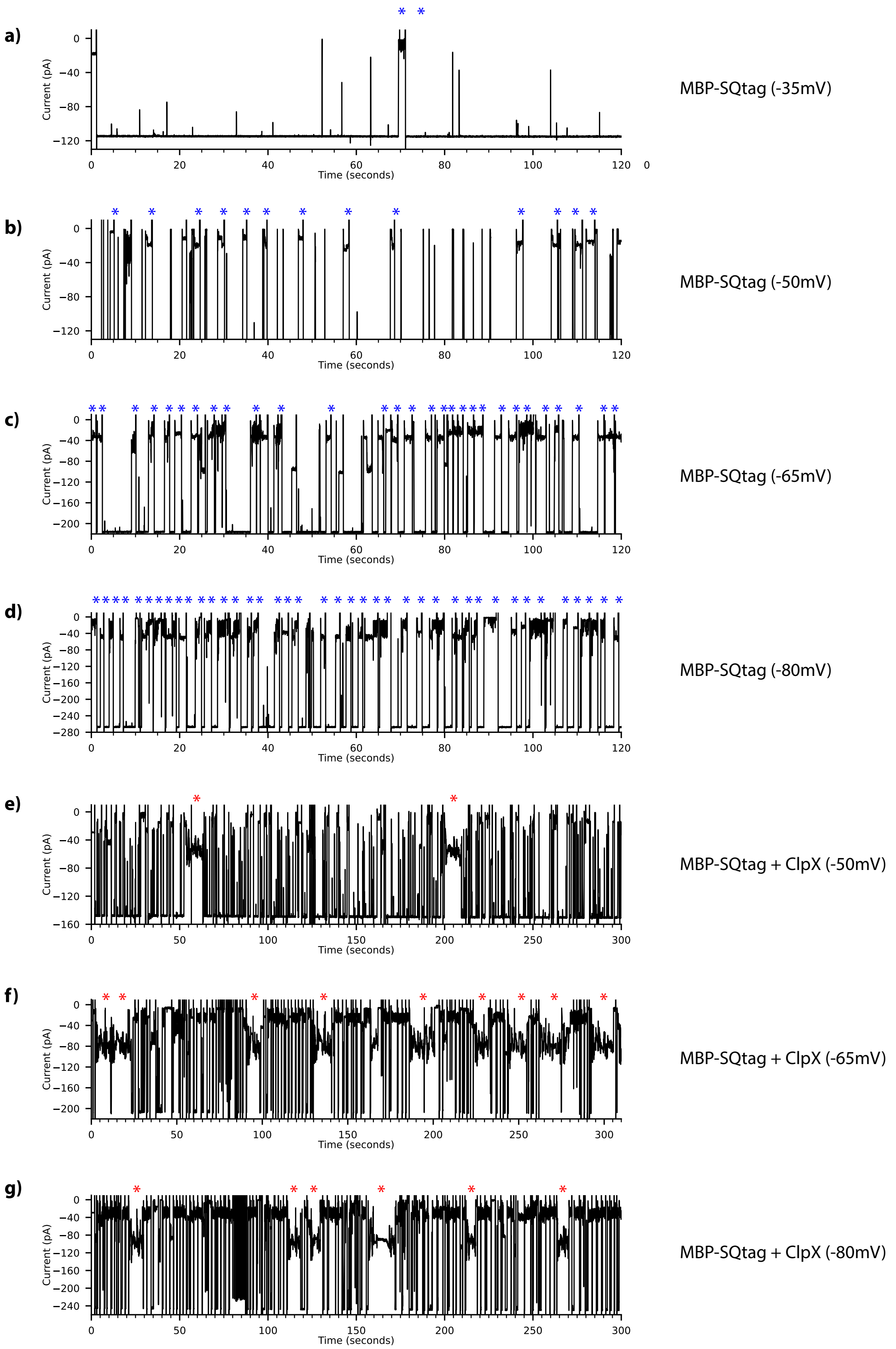


**Figure S3. Example traces recorded at different voltages for MBP-SQtag in the presence and absence of ClpX.** (a–d) Two-minute traces of MBP-SQtag capture at -30 mV, -50 mV, -65 mV, and -80 mV. (e–g) Five-minute traces in the presence of MBP-SQtag and ClpX at -50 mV,

-65 mV, and -80 mV. Blue asterisks refer to substrate capture, and the red asterisk refers to enzymatic-mediated translocation events. All recordings were made using MspA at 37°C in 1 M KGlu, 50 mM HEPES, 10 mM $MgCl_2$, pH 7.4, in the presence of 5 nM of POI, 100 nM of ClpX, 2 mM of ATP, 1.6 mM creatine phosphate, 0.4 µM creatine kinase, 1 mM DTT, and 0.5 mM EDTA.

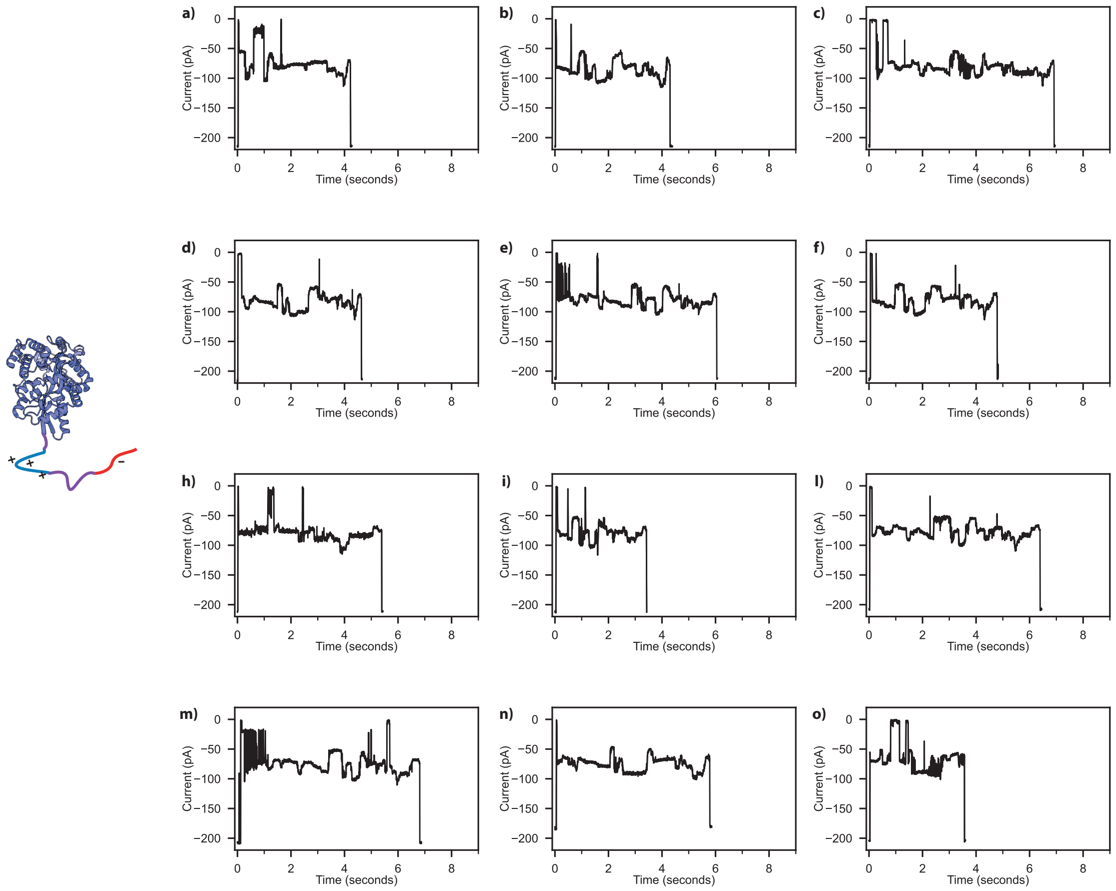


**Figure S4. Representative blocked or partial squiggles of the MBP-10R-ssrA substrate.** (a-o) Each squiggle represents a portion of the full signal, indicated by the different starting points for each trace. The MBP-10R-ssrA substrate contains only the ssrA tag without the stalling domain. Raw data were acquired using MspA in the presence of 100 nM of ClpX, 5 nM of POI, with a voltage of -65 mV applied at 37°C in 1 M K-gluconate, 50 mM HEPES, 10 mM $MgCl_2$, pH 7.4, 2 mM of ATP, 1.6 mM creatine phosphate, 0.4 µM creatine kinase, 1 mM DTT, and 0.5 mM EDTA. Traces were filtered using a low-pass Bessel filter (500Hz) for enhanced visualization.

**a)**

**b)**

**c)**

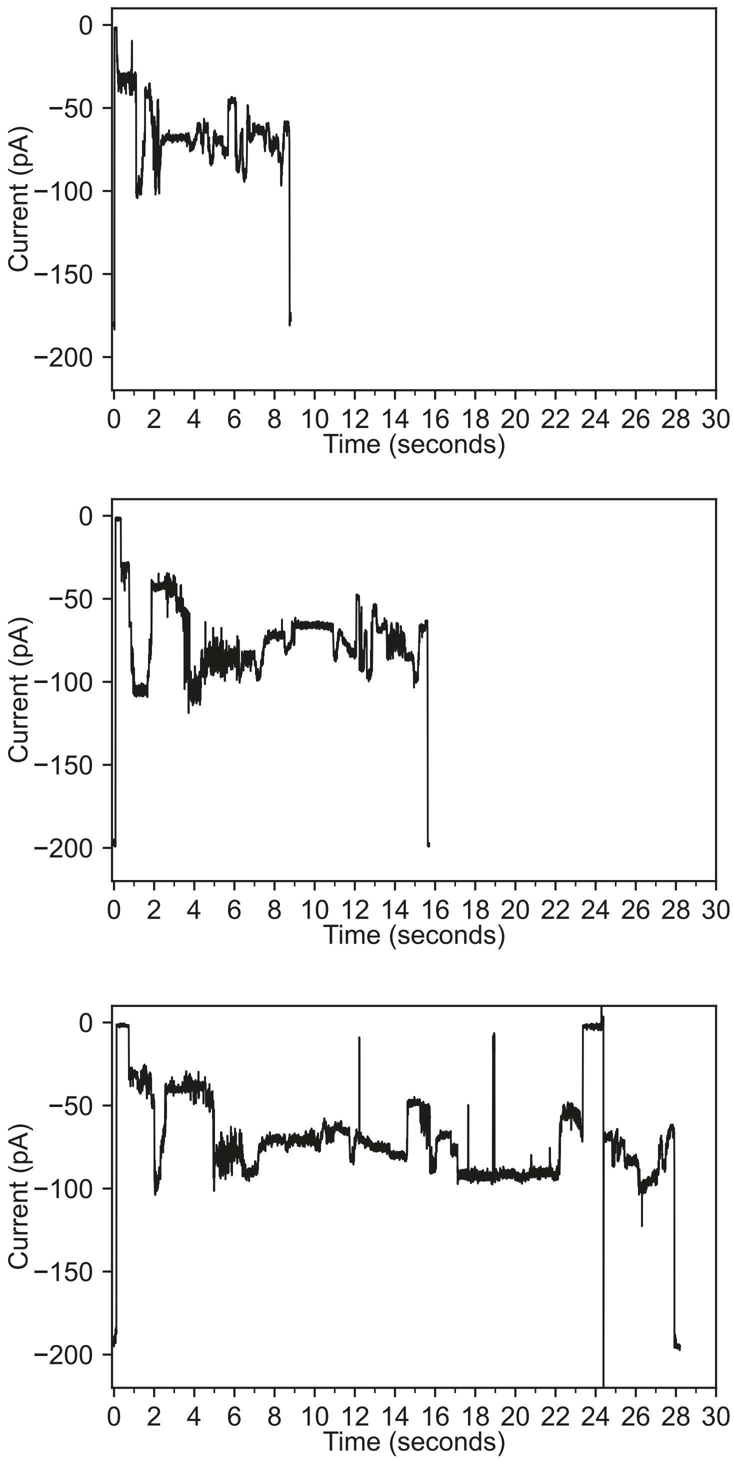


**Figure S5. Example traces recorded at different concentrations of ATP-γS for MBP-SQtag in the presence of 2 mM of ATP.** a) no ATP-γS, b) 1 mM of ATP-γS, c) 2 mM of ATP-γS. Raw data were acquired using MspA in the presence of 100 nM of ClpX, 5 nM of POI, with a voltage of -65 mV applied at 37°C in 1 M K-gluconate, 50 mM HEPES, 10 mM $MgCl_2$, pH 7.4, 2 mM of ATP, 1.6 mM creatine phosphate, 0.4 µM creatine kinase, 1 mM DTT, and 0.5 mM EDTA. Traces were filtered using a low-pass Bessel filter (500Hz) for enhanced visualization.

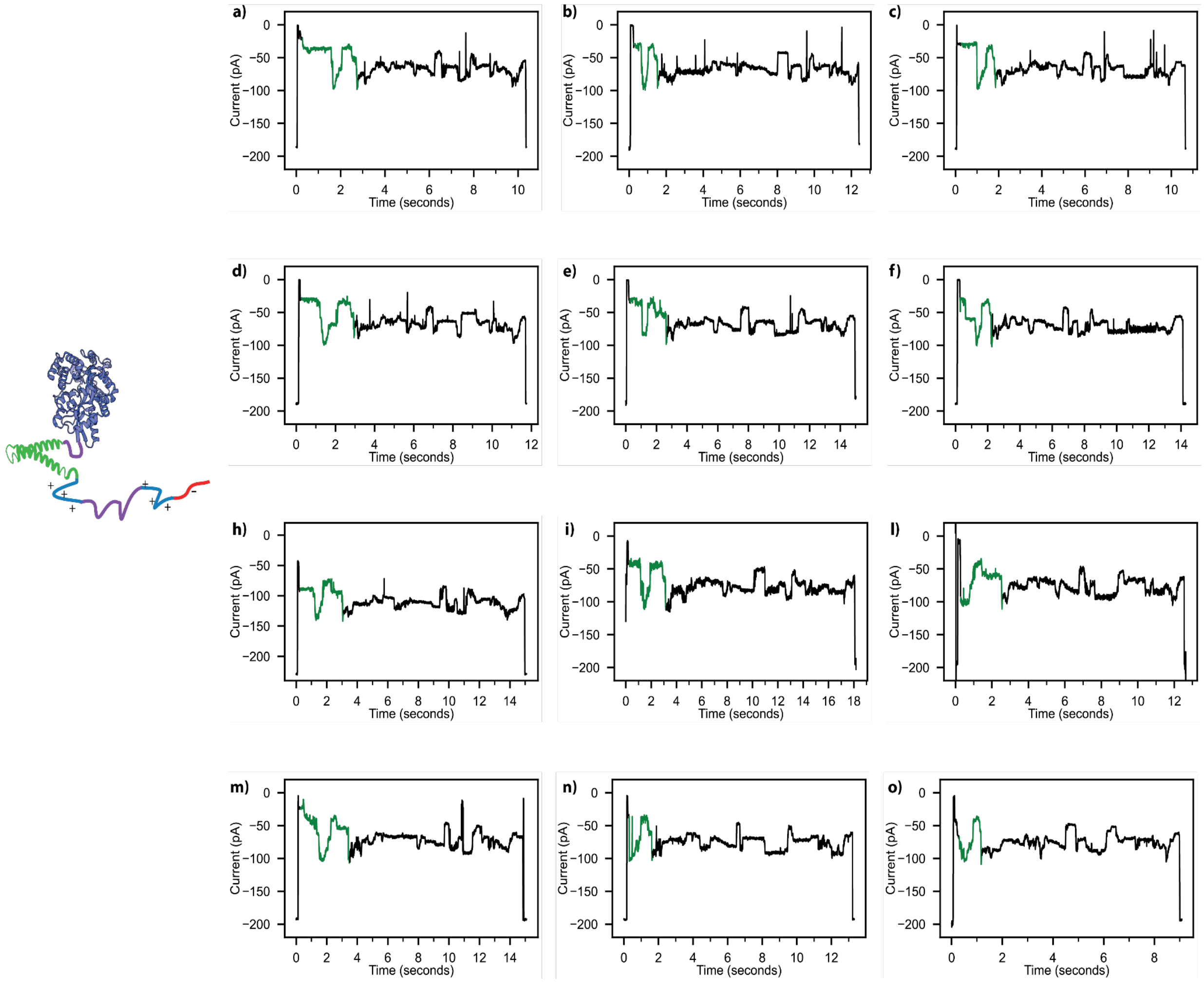


**Figure S6. Representative squiggles of the MBP-SQtag substrate**. (a-o) Traces showing two distinct regions: the stalling domain (green) and the protein domain starting from the C-terminus (black). Raw data were acquired using MspA in the presence of 100 nM of ClpX, 5 nM of POI, with a voltage of -65 mV applied at 37°C in 1 M K-gluconate, 50 mM HEPES, 10 mM $MgCl_2$, pH 7.4, 2 mM of ATP, 1.6 mM creatine phosphate, 0.4 µM creatine kinase, 1 mM DTT, and 0.5 mM EDTA. (a,o) Traces were filtered using a low-pass Bessel filter (500Hz) for enhanced visualization.

.

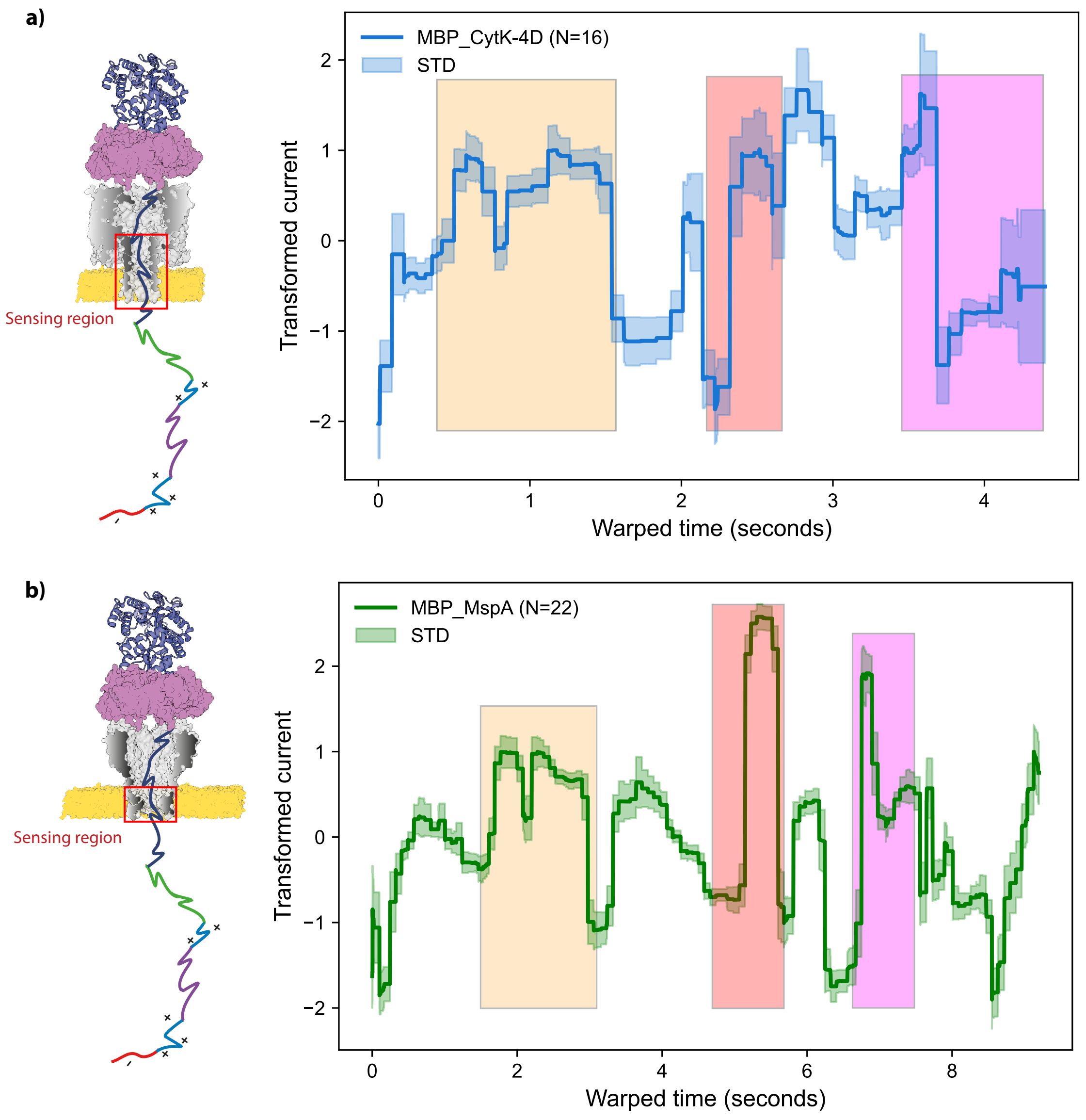


**Figure S7. Comparison of MBP-SQtag squiggle through CytK-4D and MspA nanopores.** Colored rectangles highlight portions of the trace representing the same part of the protein sequence. (a) Compiled CytK-4D squiggles, (b) Compiled MspA squiggles. MspA shows more detailed steps due to its shorter sensing region, allowing for a higher-resolution reading of the protein. In contrast, the CytK-4D trace, with a longer sensing region, provides less detailed information and covers a smaller reading window in the protein sequence. a) Trace collected at -80 mV at 37°C in 1 M KGlu, 50 mM HEPES, 10 mM $MgCl_2$, pH 7.4. b) Trace collected at -65 mV at 37°C in 1 M KGlu, 50 mM HEPES, 10 mM $MgCl_2$, pH 7.4. The traces were collected at the optimal voltage for each nanopore. CytK-4D required higher applied voltages to produce a good squiggle frequency compared to MspA.

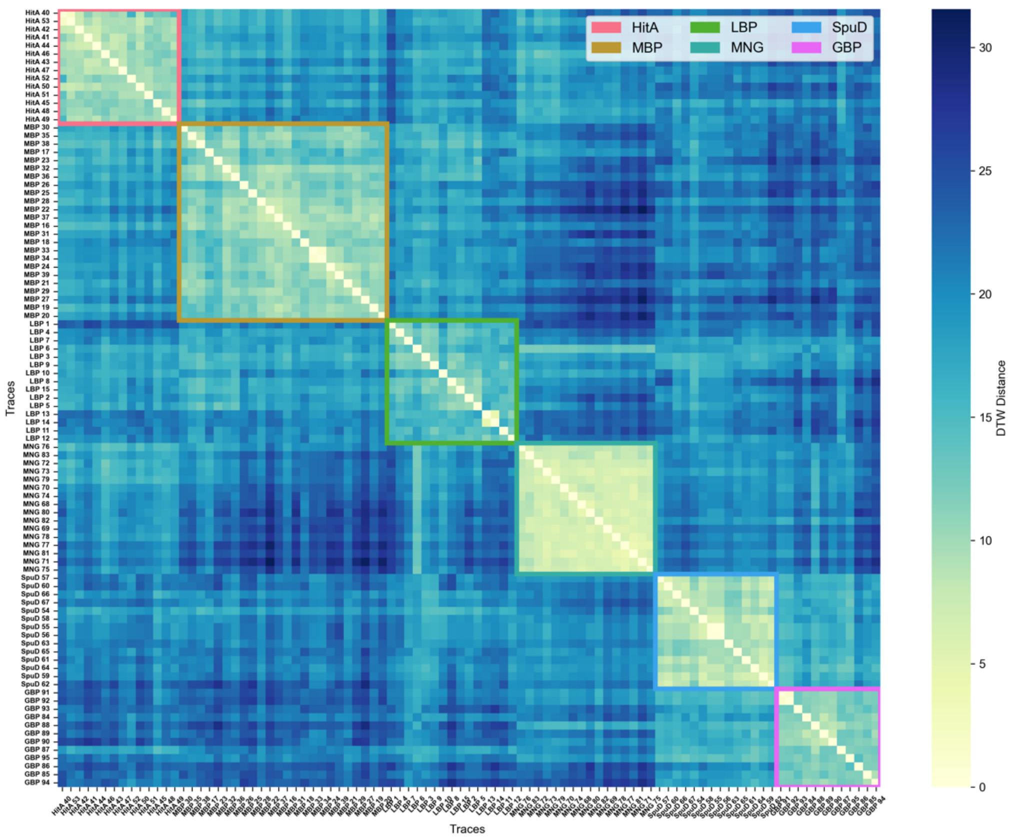


**Figure S8. Hierarchical clustering heatmap based on the dynamic time warping (DTW) distance matrix calculated between all the traces for protein identification**. The color intensity of the squares indicates similarity, with lighter colors representing higher similarity. The hierarchical clustering resulted in groups of similar traces, each corresponding to a different protein, and these groups are delineated by specific-colored lines. Regions outside these groups, with darker intensity colors, highlight differences between traces of different proteins.

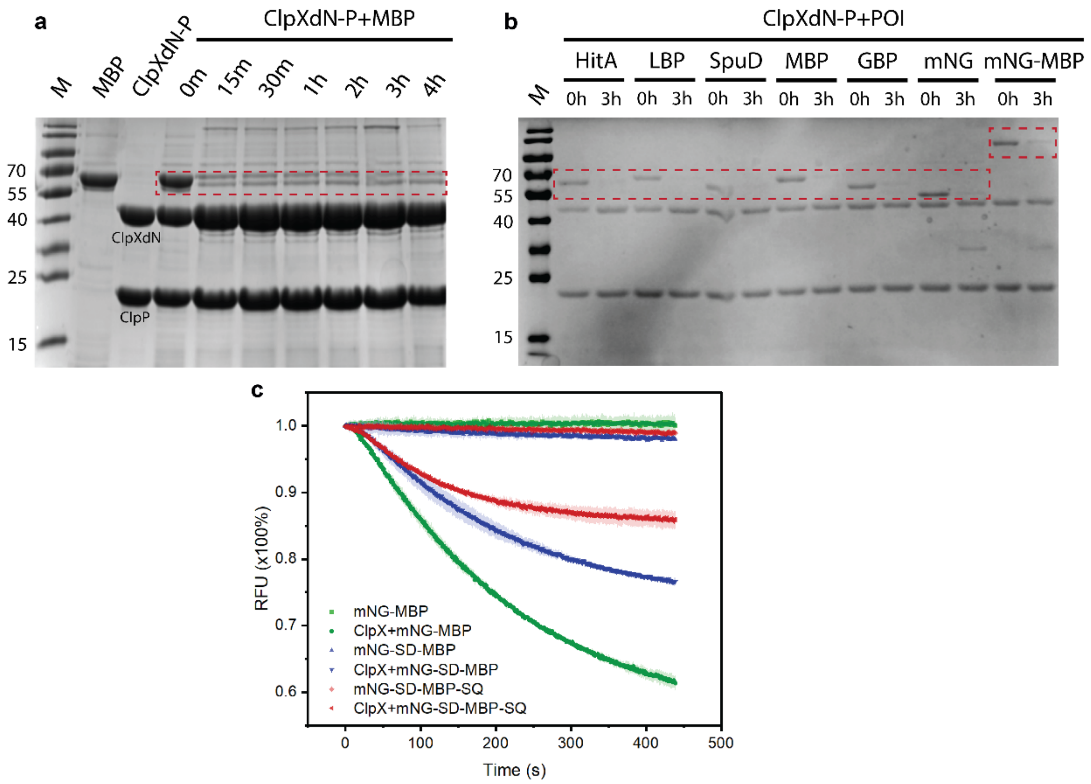


**Figure S9. Analysis of the degradation and unfolding of designed substrates by ClpXdN or together with ClpP**. (a) Degradation of the MBP-SD substrate by ClpXdN and ClpP over time. Reactions were carried out in a 40 µL mixture containing 50 mM HEPES (pH 7.5), 5 mM $MgCl_2$, 200 mM NaCl, 4 mM ATP, an ATP regeneration system, 1 mM DTT, 500 nM ClpX and ClpP, and 66 nM MBP, incubated at 37°C. After the reaction, the samples were mixed with 4x loading dye and boiled at 95°C for 10 minutes. (b) Degradation assay of seven substrates (HitA-SQtag, LBP-SQtag, SpuD-SQtag, MBP-SQtag, GBP-SQtag, mNG-SQtag, mNG-SD-MBP-SQtag) used in protein identification (Figure 1). The same reaction conditions were applied as in (a), except for the substrates. (c) Bulking unfolding assay of mNG-MBP-ssrA (mNG-MBP), mNG-SD-MBP-ssrA (mNG-SD-MBP), and mNG-SD-MBP-SQtag (mNG-SD-MBP-SQ) by ClpXdN alone. Reactions were conducted at 37°C in a 150 µL mixture containing 50 mM HEPES (pH 7.5), 5 mM $MgCl_2$, corresponding potassium salts, 4 mM ATP, an ATP regeneration system, 1 mM DTT, 250 nM ClpX, and 50 nM substrates. Each unfolding assay was performed in triplicate.

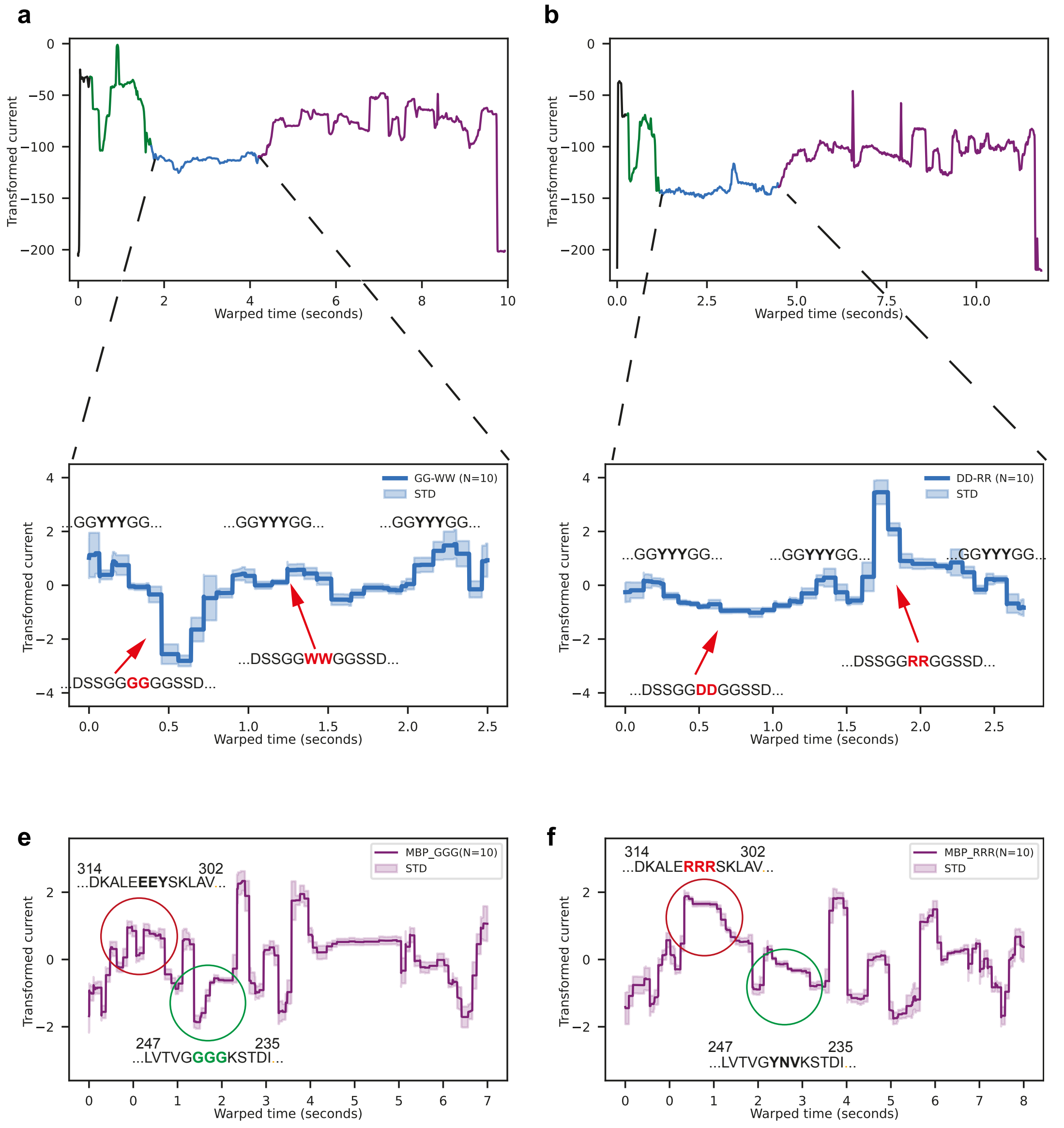


**Figure S10. Multiple amino acid substitutions in an unstructured synthetic polypeptide (SAA) and in MBP**. (a, b) Representative traces of the MBP-SAA(WW,GG)-SQtag and MBP-SAA(RR,DD)-SQtag substrates, highlighting three distinct regions: the stalling domain (green), unstructured polypeptide (blue), and protein domain (purple). (a) Introduction of two glycine (GG) and tryptophan (WW) substitutions in the unstructured polypeptide (between YYY) resulting in a current enhancement for GG and a larger current blockade for WW. (b) DD and RR substitutions in the same region cause a current block for RR and no significant change is observed with DD. (e) Introduction of a triple GGG substitution in the MBP native domain (MBP-GGG-SD) results in an increase in current associated with the small size of the amino acids. (f) Introduction of a triple RRR mutation in the MBP protein domain (MBP-RRR-SD) leads to a significant current blockade associated with the interaction between the positively charged amino acids and the negatively charged residues in the sensing region of the pore. All traces were obtained using Msp in the presence of 100 nM of ClpX, 5 nM of POI, with a voltage of -65 mV applied at 37°C in 1 M K-gluconate, 50 mM HEPES, 10 mM $MgCl_2$, pH 7.4, 2 mM of ATP, 1.6 mM creatine phosphate, 0.4 µM creatine kinase, 1 mM DTT, and 0.5 mM EDTA.

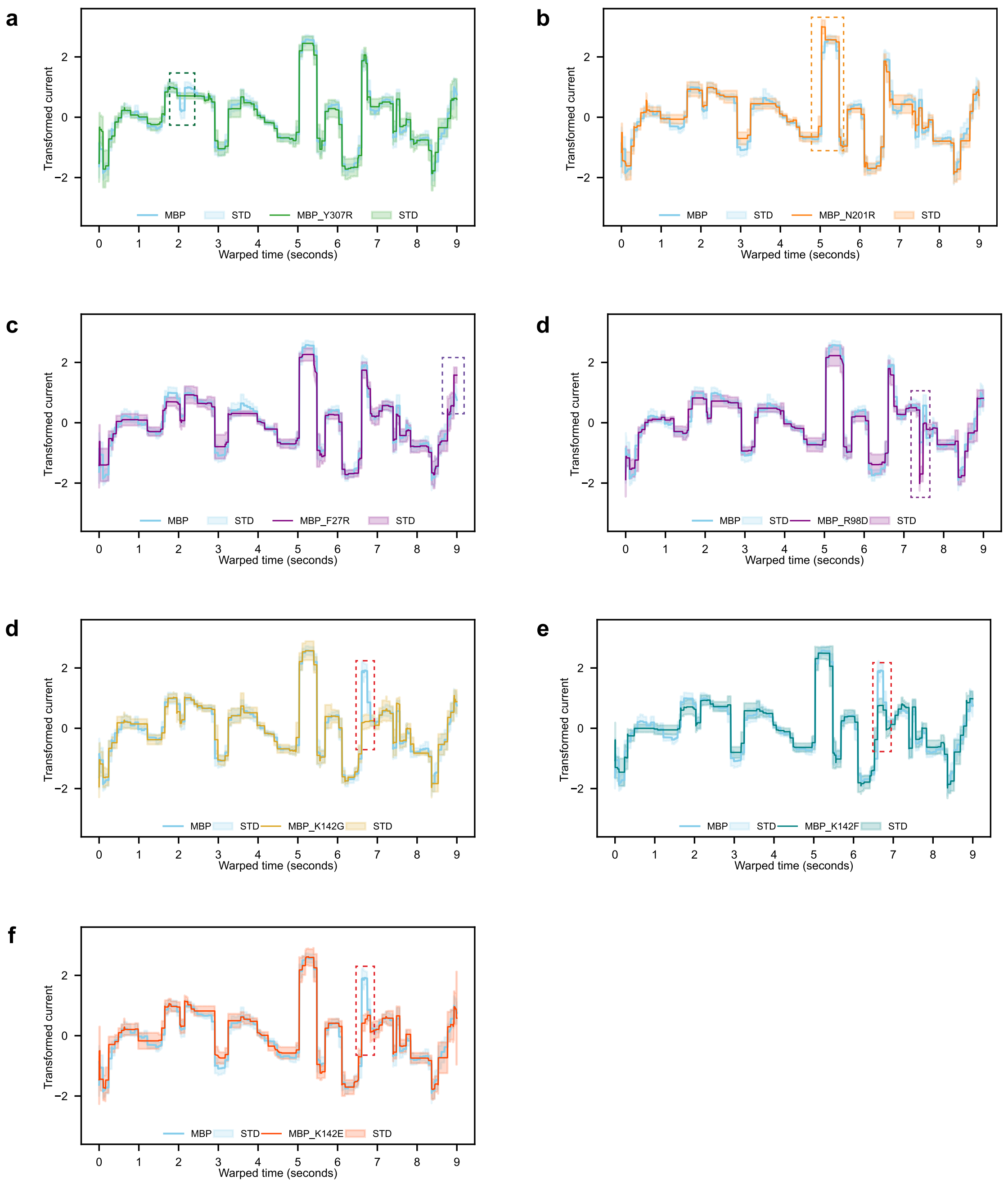


**Figure S11**. **Single point mutation in MBP protein**. Aligned full traces of MBP single point mutations compared to the MBP-SQtag (light blue, reference). Rectangles highlight the signal variation induced by the mutation. (a) MBP-Y307R (N=17), (b) MBP-N201R (N=9), (c) MBP-F27R (N=10), (d) MBP-R98D (N=10), (e) MBP-K142G (N=9), (f) MBP-K142F (N=15), (g) MBP-K142E (N=23). All traces were obtained using MspA in the presence of 100 nM of ClpX, 5 nM of POI, with a voltage of -65 mV applied at 37°C in 1 M K-gluconate, 50 mM HEPES, 10 mM $MgCl_2$, pH 7.4, 2 mM of ATP, 1.6 mM creatine phosphate, 0.4 µM creatine kinase, 1 mM DTT, and 0.5 mM EDTA.

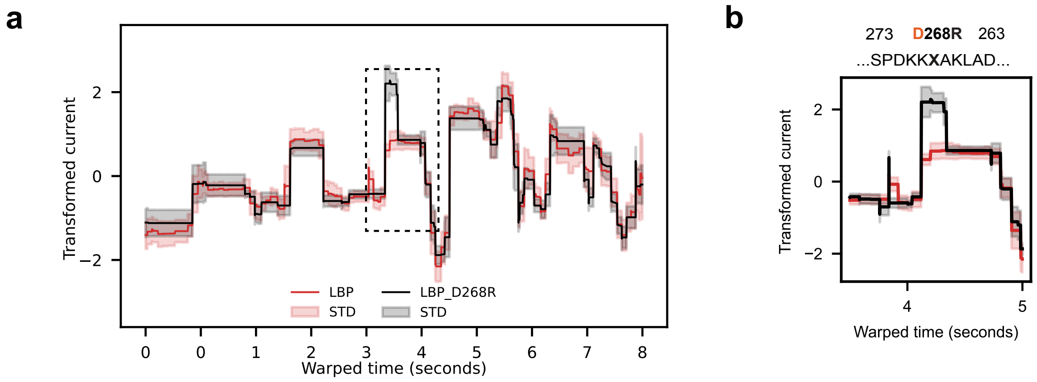


**Figure S12. Single arginine substitution in LBP protein. (**a) Aligned full traces of LBP-D268R-SQtag (N=12) to the reference LBP-SQtag (N=15). (b) Highlighted region where the mutation significantly affects the trace. All traces were obtained using MspA in the presence of 100 nM of ClpX, 5 nM of POI, with a voltage of -65 mV applied at 37°C in 1 M K-gluconate, 50 mM HEPES, 10 mM $MgCl_2$, pH 7.4, 2 mM of ATP, 1.6 mM creatine phosphate, 0.4 µM creatine kinase, 1 mM DTT, and 0.5 mM EDTA

.

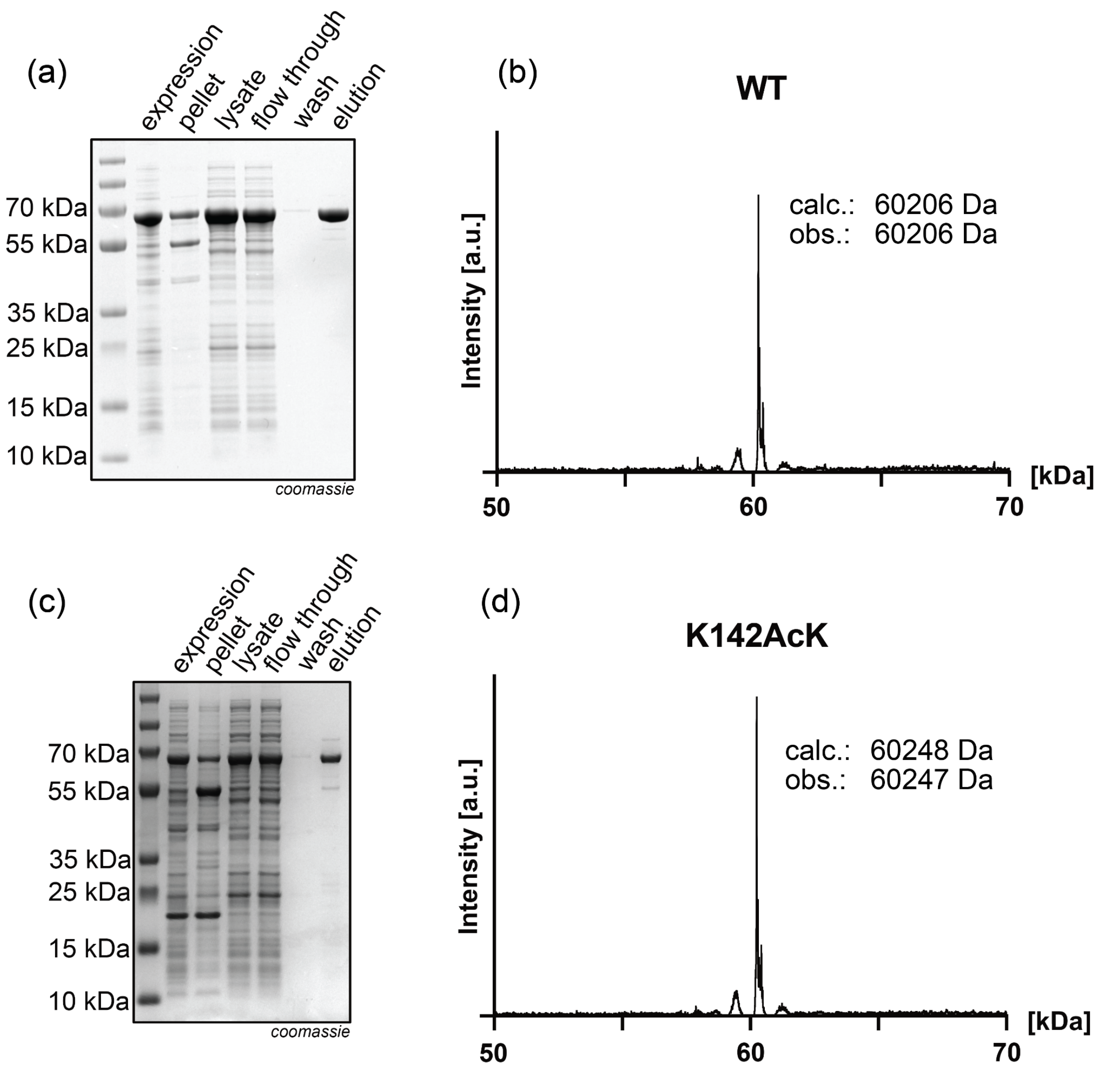


**Figure S13. SDS-PAGE and mass spectrometry analysis of MBP-SQtag and MBP-K142AcK-SQtag proteins.** 12% SDS analysis of (a) MBP-SQtag and (c) MBP-K142AcK-SQtag. Mass spectra of (b) MBP-SQtag and (d) MBP-K142AcK-SQtag.

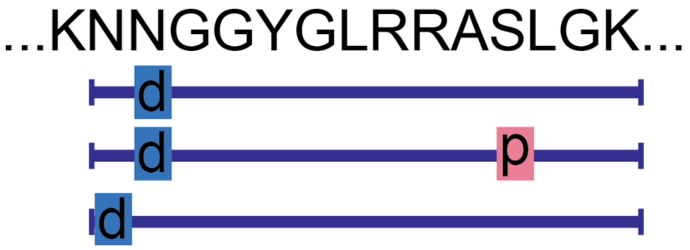


| Peptide | Length | m/z | RT | Peak Area | |
|---|---|---|---|---|---|
| | | | | MBP-PKA1_unmod | MBP-PKA1_phos |
| NN(+0.98)GGYGLRRASLGK | 1462.8 | 366.70 | 3.37 | 2.13E+07 | not detected |
| NN(+0.98)GGYGLRRAS(+79.97)LGK | 1542.7 | 386.69 | 3.31 | not detected | 2.77E+06 |
| N(+0.98)NGGYGLRRASLGK | 1462.8 | 732.39 | 3.37 | 7.30E+05 | not detected |

**Figure S14. Mass spectrometry analysis of phosphorylation in MBP$^{PKA1}$.** The table shows the detected peptides of the phosphorylation site in MBP$^{PKA1}$ for the unmodified sample (MBP1-PKA_unmod) and the phosphorylated sample (MBP-PKA1_phos).

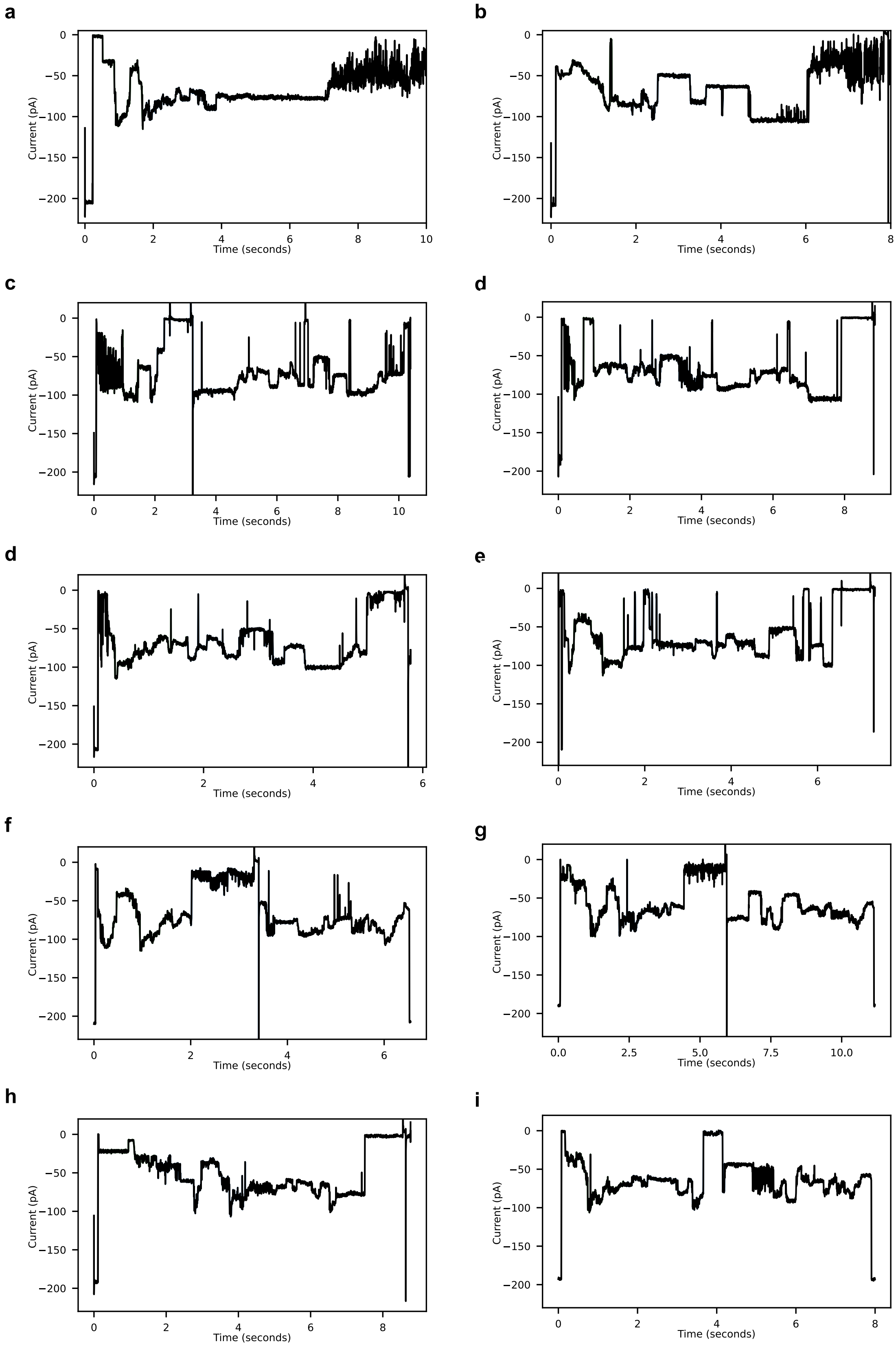


**Figure S15. Example of blocked squiggles.** (a, i) Representative traces excluded from data analysis showing significant signal blockage covering a substantial portion of the protein sequence. All traces were obtained using MspA in the presence of 100 nM of ClpX, 5 nM of POI, with a voltage of -65 mV applied at 37°C in 1 M K-gluconate, 50 mM HEPES, 10 mM $MgCl_2$, pH 7.4, 2 mM of ATP, 1.6 mM creatine phosphate, 0.4 µM creatine kinase, 1 mM DTT, and 0.5 mM EDTA. Traces were filtered using a low-pass Bessel filter (500Hz) for enhanced visualization.

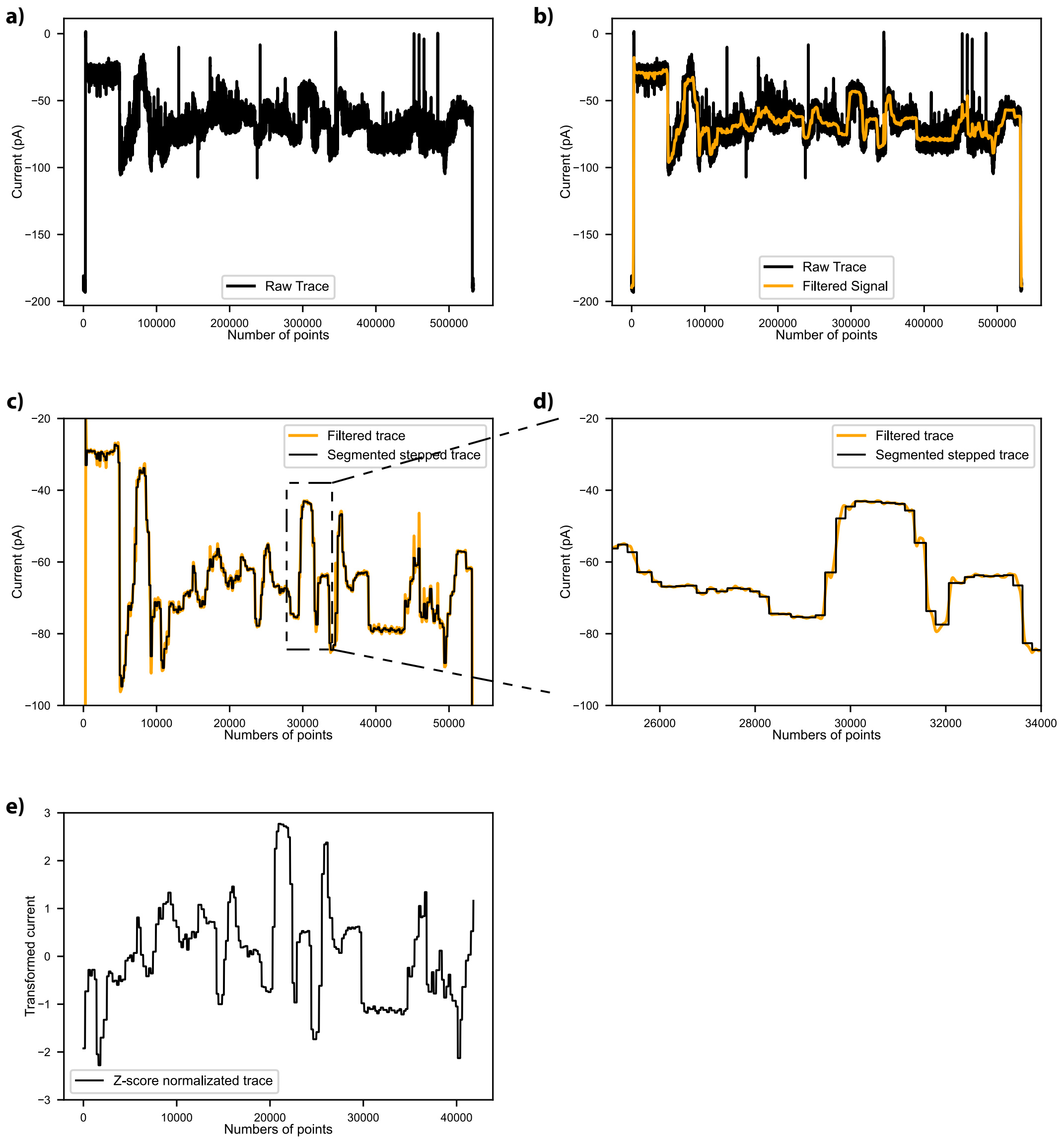


**Figure S16. Data analysis processing workflow for a representative nanopore trace of MBP-SQtag**. (a) Raw trace segmented by continuous measurement intervals. (b) Trace filtered using a low-pass Bessel filter (50 Hz) to remove the noise. (c) Transformation of the continuous signal into a step-shaped signal through change point detection (PELT algorithm), with current values averaged between detected change points. (d) Extraction of specific portions to better visualize the step-shaped signal. (e) Normalization of the trace using Z-score normalization to highlight current signal variations relative to the average.

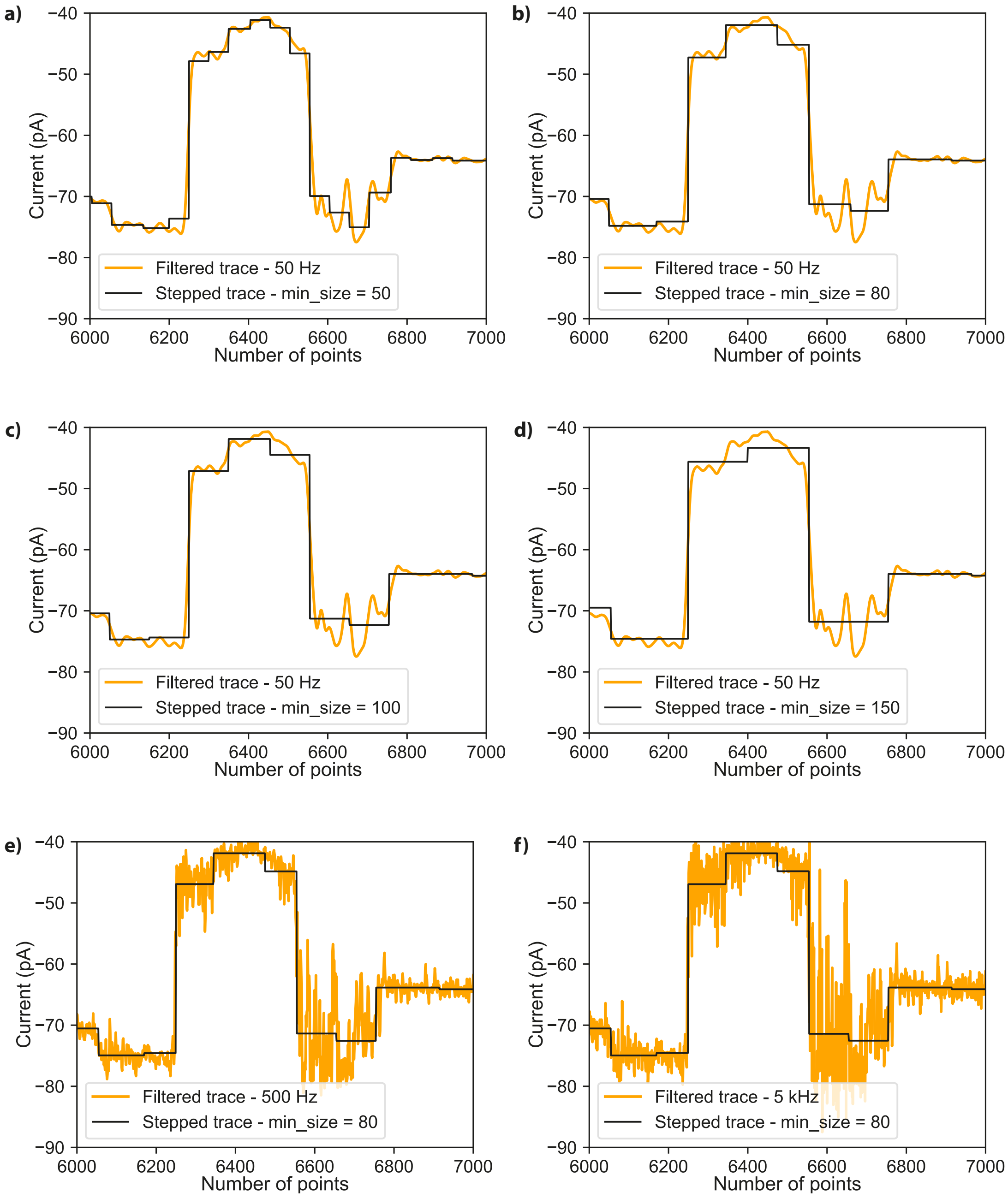


**Figure S17. Zoomed examples of a portion of MBP traces before and after applying the change-point detection PELT algorithm.** (a–d) Traces filtered with a 50 Hz low-pass Bessel filter and processed using the PELT *min_size* parameter varied between 50 and 150 ms. (e, f) Traces filtered at 500 Hz and 5 kHz and processed with *min_size* fixed at 80 ms.

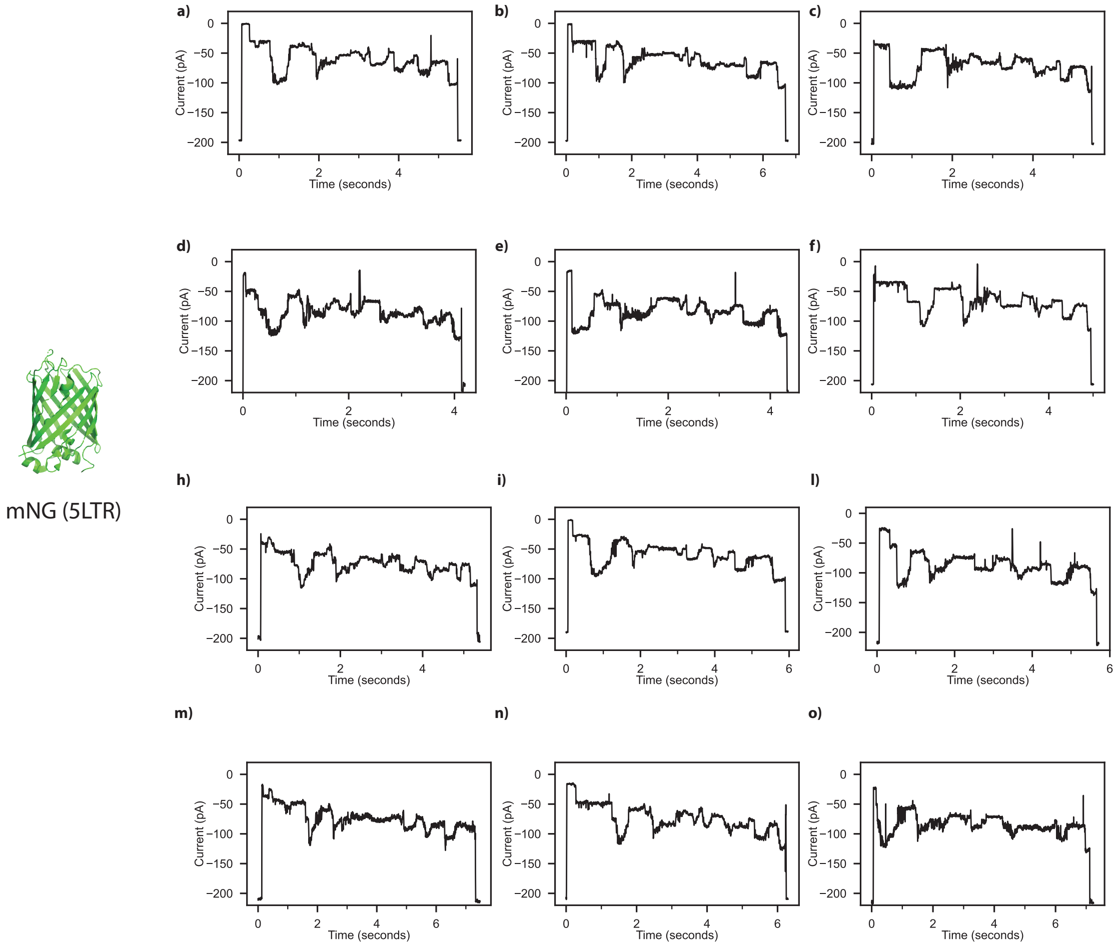


**Figure S18**. **Representative squiggles of the mNG-SQtag substrate.** Raw traces were acquired using MspA in the presence of 100 nM of ClpX, 5 nM of POI, with a voltage of -65 mV applied at 37°C in 1 M K-gluconate, 50 mM HEPES, 10 mM $MgCl_2$, pH 7.4, 2 mM of ATP, 1.6 mM creatine phosphate, 0.4 µM creatine kinase, 1 mM DTT, and 0.5 mM EDTA. (a-o) Traces were filtered using a low-pass Bessel filter (500Hz) for enhanced visualization.

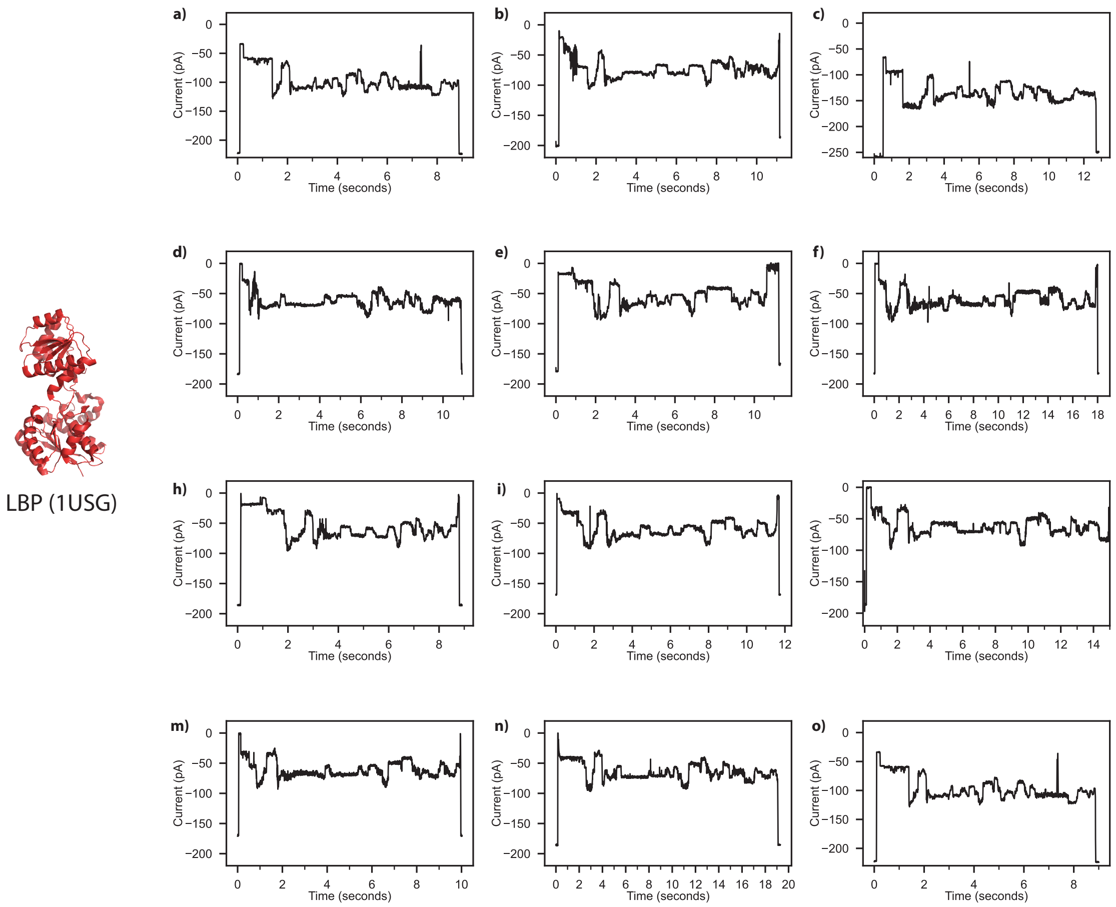


**Figure19. Representative squiggles of the LBP-SQtag substrate**. Raw traces were acquired using MspA in the presence of 100 nM of ClpX, 5 nM of POI, with a voltage of -65 mV applied at 37°C in 1 M K-gluconate, 50 mM HEPES, 10 mM $MgCl_2$, pH 7.4, 2 mM of ATP, 1.6 mM creatine phosphate, 0.4 µM creatine kinase, 1 mM DTT, and 0.5 mM EDTA. (a-o) Traces were filtered using a low-pass Bessel filter (500Hz) for enhanced visualization.

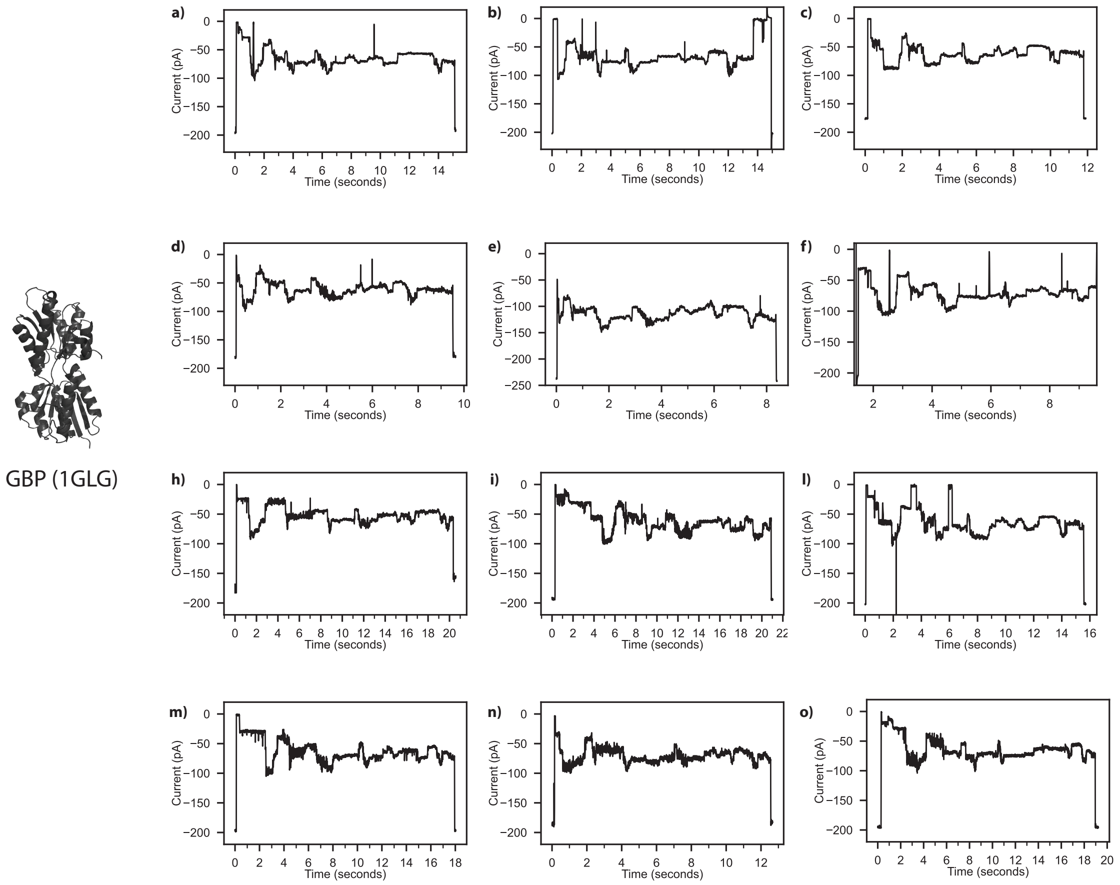


**Figure20. Representative squiggles of the GBP-SQtag substrate**. Raw traces were acquired using MspA in the presence of 100 nM of ClpX, 5 nM of POI, with a voltage of -65 mV applied at 37°C in 1 M K-gluconate, 50 mM HEPES, 10 mM $MgCl_2$, pH 7.4, 2 mM of ATP, 1.6 mM creatine phosphate, 0.4 µM creatine kinase, 1 mM DTT, and 0.5 mM EDTA. (a-o) Traces were filtered using a low-pass Bessel filter (500Hz) for enhanced visualization.

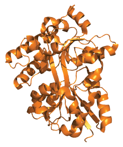

SpuD (3TTM)

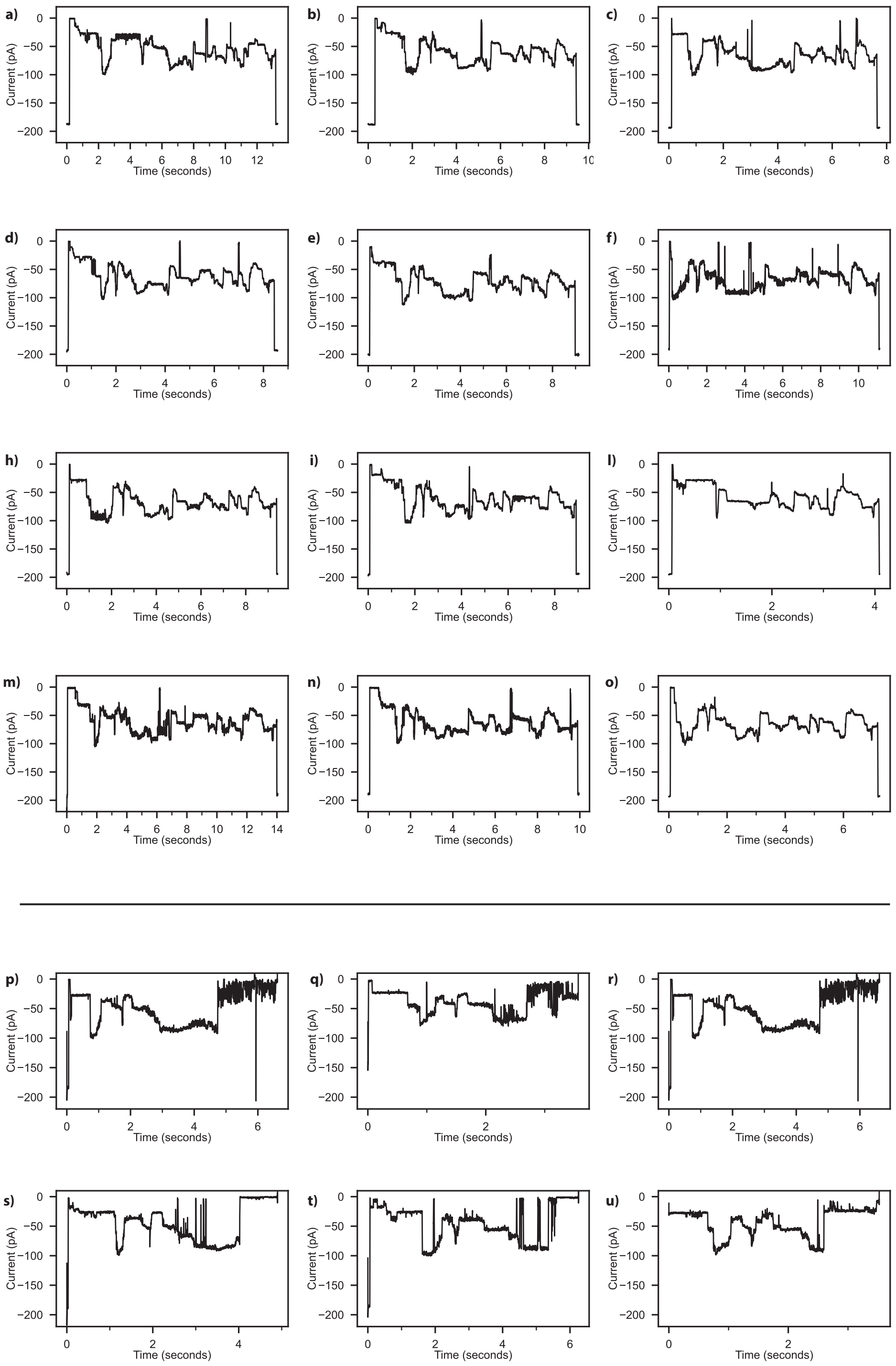


**Figure21. Representative squiggles of the SpuD-SQtag substrate**. Raw traces were acquired using MspA in the presence of 100 nM of ClpX, 5 nM of POI, with a voltage of -65 mV applied at 37°C in 1 M K-gluconate, 50 mM HEPES, 10 mM $MgCl_2$, pH 7.4, 2 mM of ATP, 1.6 mM creatine phosphate, 0.4 µM creatine kinase, 0.5 mM EDTA either with 1 mM DTT (a–o) or without DTT (p–u). (a-u) Traces were filtered using a low-pass Bessel filter (500Hz) for enhanced visualization.

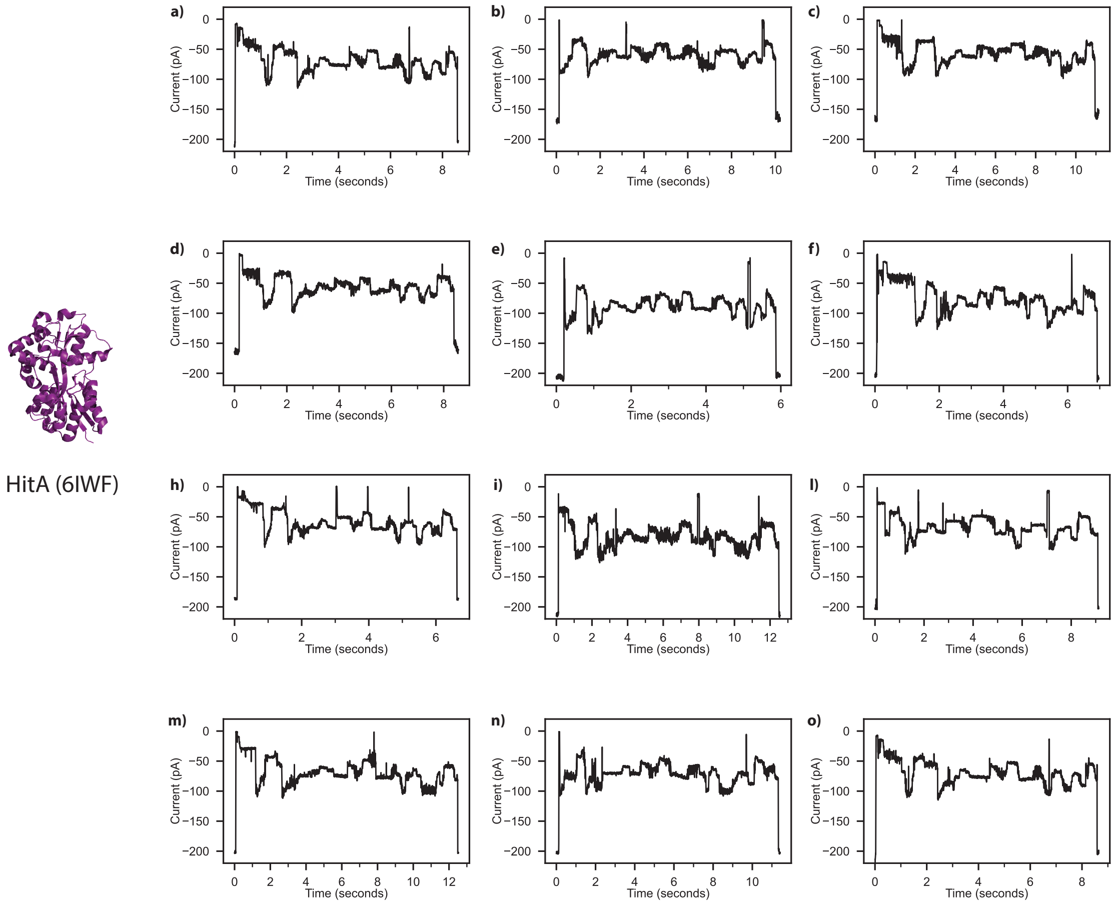


**Figure22. Representative squiggles of the HitA-SQtag substrate.** Raw traces were acquired using MspA in the presence of 100 nM of ClpX, 5 nM of POI, with a voltage of -65 mV applied at 37°C in 1 M K-gluconate, 50 mM HEPES, 10 mM $MgCl_2$, pH 7.4, 2 mM of ATP, 1.6 mM creatine phosphate, 0.4 µM creatine kinase, 1 mM DTT, and 0.5 mM EDTA. (a-o) Traces were filtered using a low-pass Bessel filter (500Hz) for enhanced visualization.

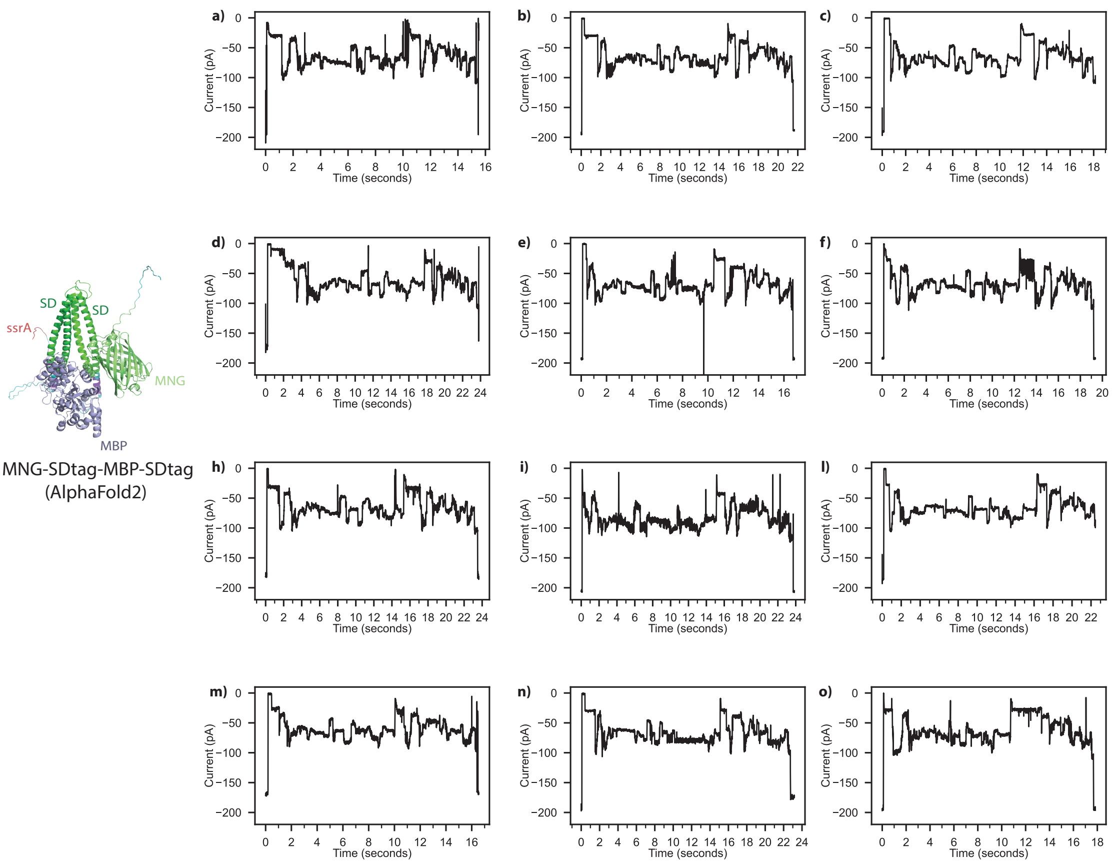


**Figure S23. Representative squiggles of the MNG-SD-MBP-SQtag substrate.** Raw traces were acquired using MspA in the presence of 100 nM of ClpX, 5 nM of POI, with a voltage of -65 mV applied at 37°C in 1 M K-gluconate, 50 mM HEPES, 10 mM $MgCl_2$, pH 7.4, 2 mM of ATP, 1.6 mM creatine phosphate, 0.4 µM creatine kinase, 1 mM DTT, and 0.5 mM EDTA. (a-o) Traces were filtered using a low-pass Bessel filter (500Hz) for enhanced visualization.

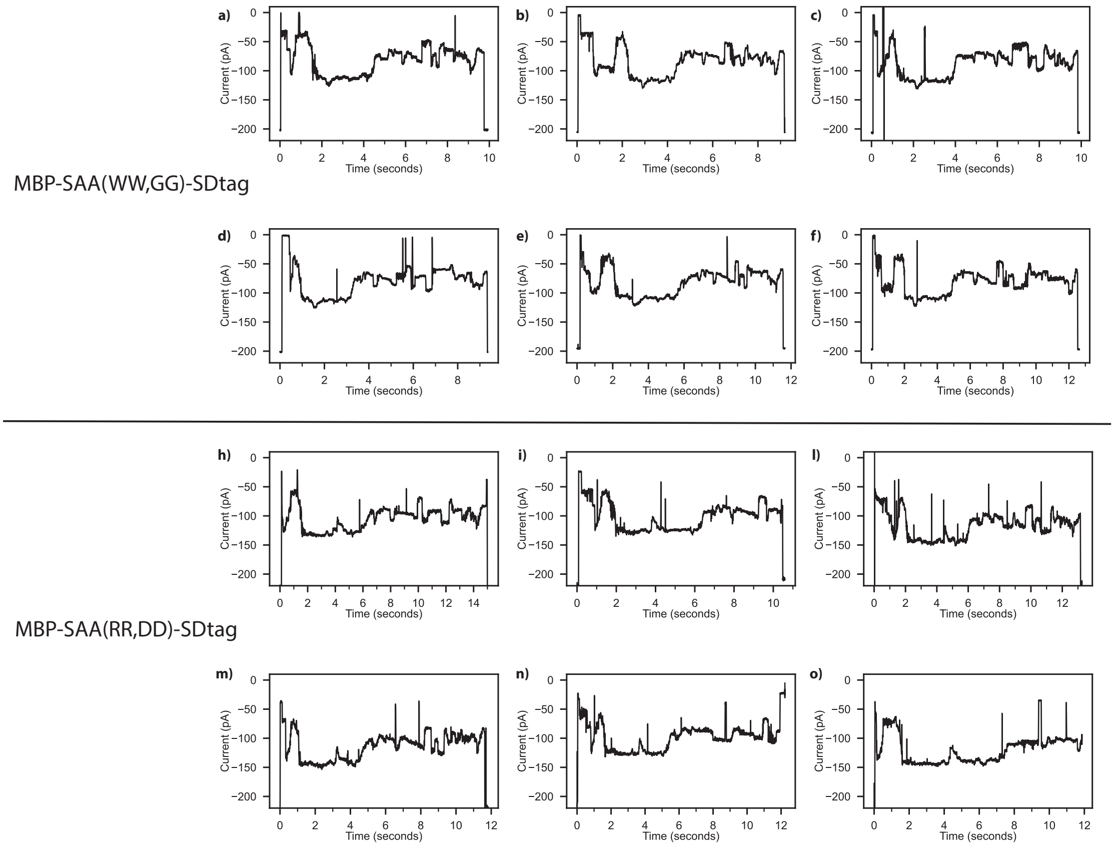


**Figure S24. Representative squiggles of the MBP-SAA-SQtag substrates.** (a-f) MBP-SAA(WW,GG)-SQtag and (h-o) MBP-SAA(WW,GG)-SQtag. Raw traces were acquired using MspA in the presence of 100 nM of ClpX, 5 nM of POI, with a voltage of -65 mV applied at 37°C in 1 M K-gluconate, 50 mM HEPES, 10 mM $MgCl_2$, pH 7.4, 2 mM of ATP, 1.6 mM creatine phosphate, 0.4 µM creatine kinase, 1 mM DTT, and 0.5 mM EDTA. (a-o) Traces were filtered using a low-pass Bessel filter (500Hz) for enhanced visualization.

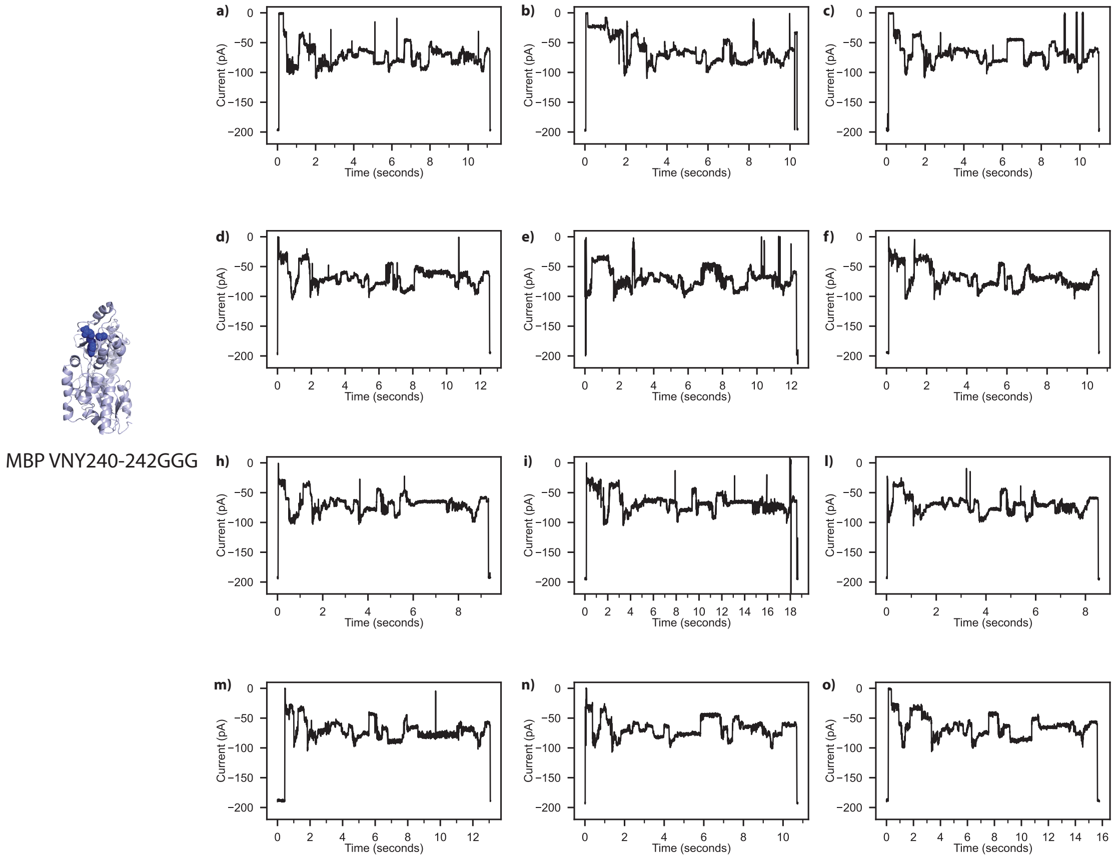


**Figure S25. Representative squiggles of the MBP-GGG-SQtag substrate.** Raw traces were acquired using MspA in the presence of 100 nM of ClpX, 5 nM of POI, with a voltage of -65 mV applied at 37°C in 1 M K-gluconate, 50 mM HEPES, 10 mM $MgCl_2$, pH 7.4, 2 mM of ATP, 1.6 mM creatine phosphate, 0.4 µM creatine kinase, 1 mM DTT, and 0.5 mM EDTA. (a-o) Traces were filtered using a low-pass Bessel filter (500Hz) for enhanced visualization.

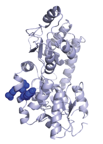

MBP YEE307-309RRR

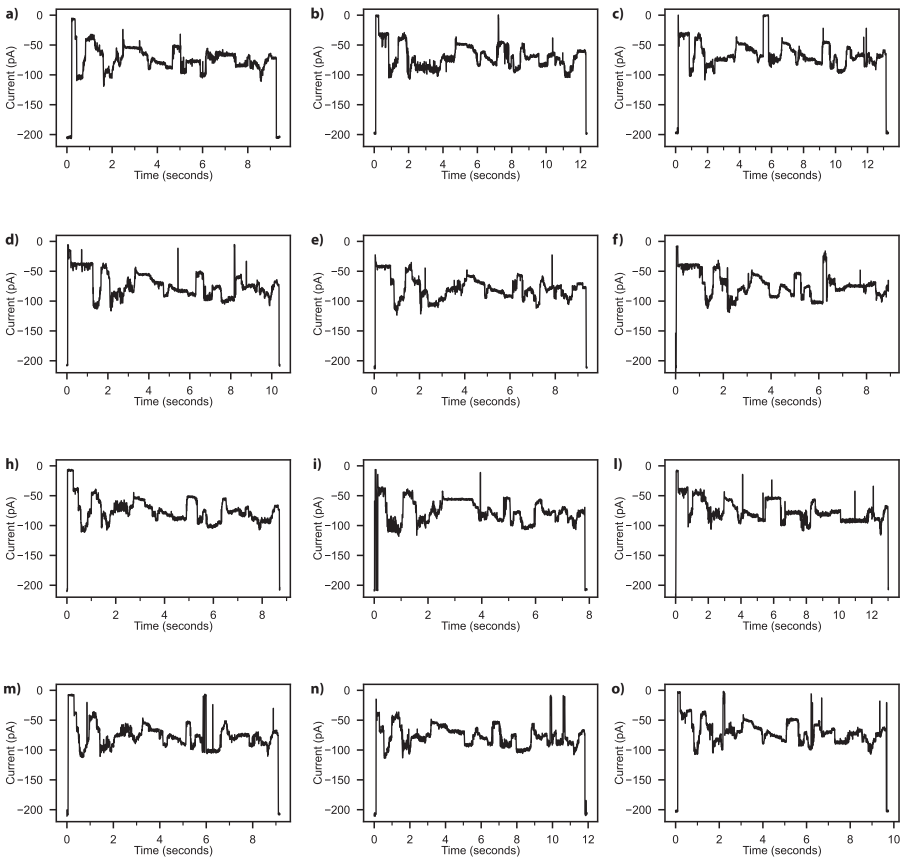


**Figure S26. Representative squiggles of the MBP-RRR-SQtag substrate.** Raw traces were acquired using MspA in the presence of 100 nM of ClpX, 5 nM of POI, with a voltage of -65 mV applied at 37°C in 1 M K-gluconate, 50 mM HEPES, 10 mM $MgCl_2$, pH 7.4, 2 mM of ATP, 1.6 mM creatine phosphate, 0.4 µM creatine kinase, 1 mM DTT, and 0.5 mM EDTA. (a-o) Traces were filtered using a low-pass Bessel filter (500Hz) for enhanced visualization.

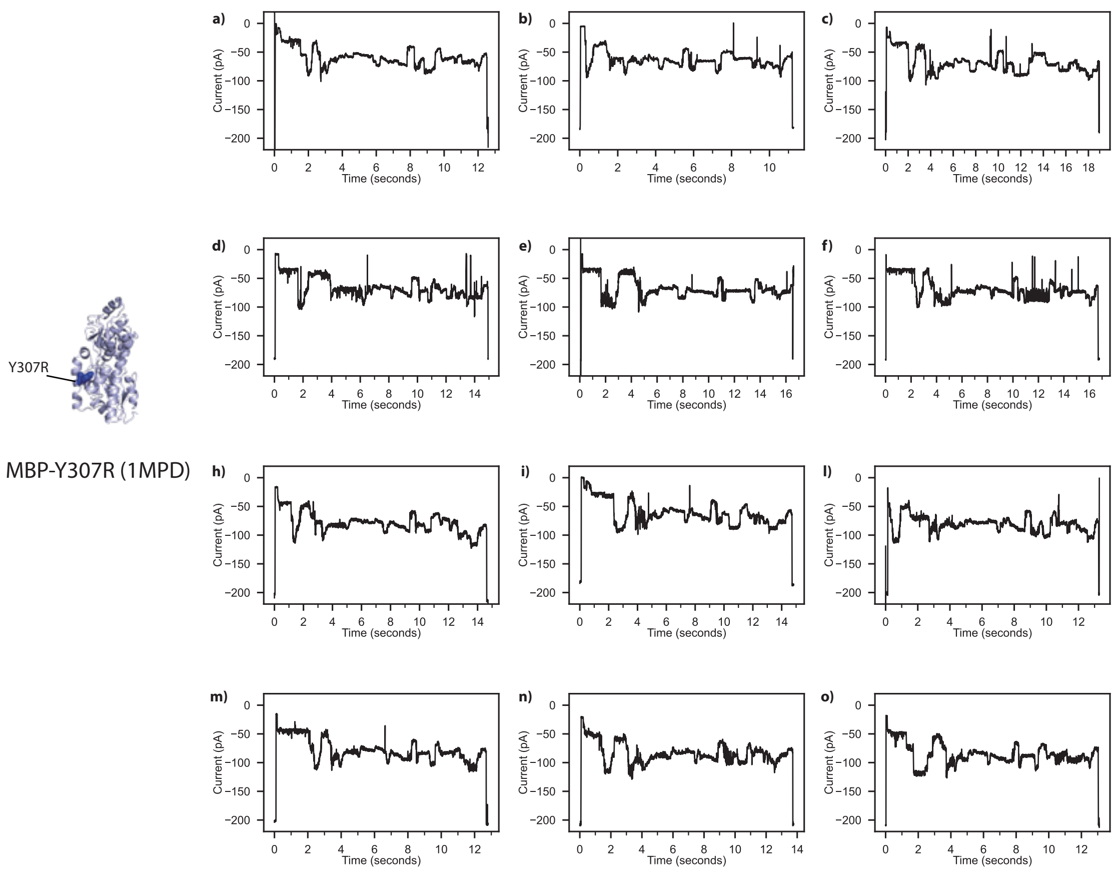


**Figure S27. Representative squiggles of the MBP-Y307R-SQtag substrate**. Raw traces were acquired using MspA in the presence of 100 nM of ClpX, 5 nM of POI, with a voltage of -65 mV applied at 37°C in 1 M K-gluconate, 50 mM HEPES, 10 mM $MgCl_2$, pH 7.4, 2 mM of ATP, 1.6 mM creatine phosphate, 0.4 µM creatine kinase, 1 mM DTT, and 0.5 mM EDTA. (a-o) Traces were filtered using a low-pass Bessel filter (500Hz) for enhanced visualization.

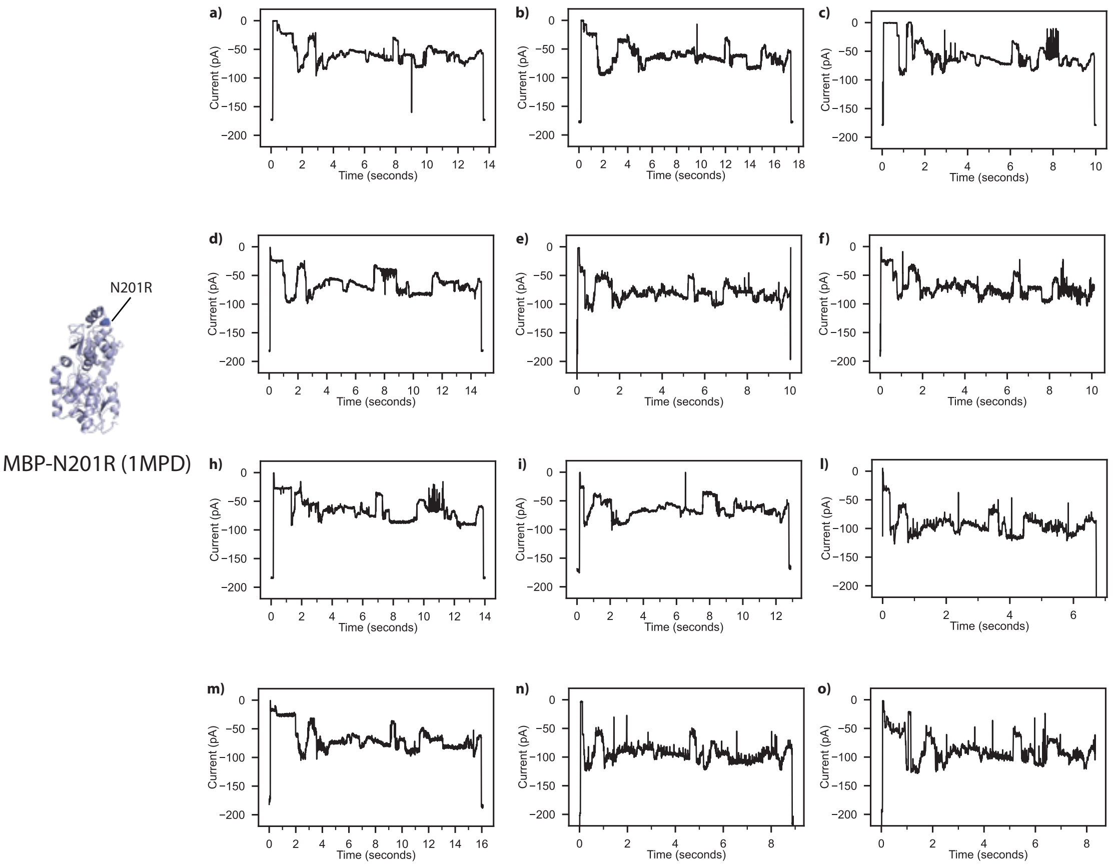


**Figure S28. Representative squiggles of the MBP-N201R-SQtag substrate**. Raw traces were acquired using MspA in the presence of 100 nM of ClpX, 5 nM of POI, with a voltage of -65 mV applied at 37°C in 1 M K-gluconate, 50 mM HEPES, 10 mM $MgCl_2$, pH 7.4, 2 mM of ATP, 1.6 mM creatine phosphate, 0.4 µM creatine kinase, 1 mM DTT, and 0.5 mM EDTA. (a-o) Traces were filtered using a low-pass Bessel filter (500Hz) for enhanced visualization.

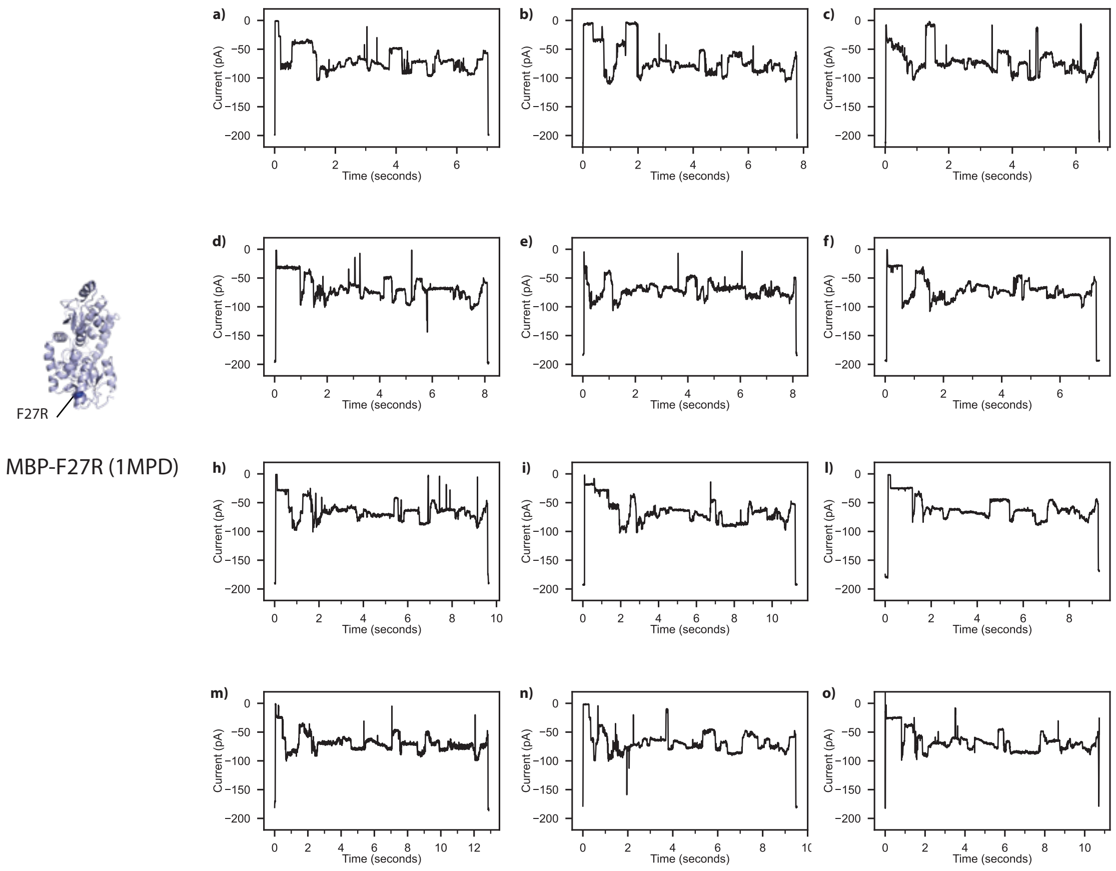


**Figure S29. Representative squiggles of the MBP-F27R-SQtag substrate**. Raw traces were acquired using MspA in the presence of 100 nM of ClpX, 5 nM of POI, with a voltage of -65 mV applied at 37°C in 1 M K-gluconate, 50 mM HEPES, 10 mM $MgCl_2$, pH 7.4, 2 mM of ATP, 1.6 mM creatine phosphate, 0.4 µM creatine kinase, 1 mM DTT, and 0.5 mM EDTA. (a-o) Traces were filtered using a low-pass Bessel filter (500Hz) for enhanced visualization.

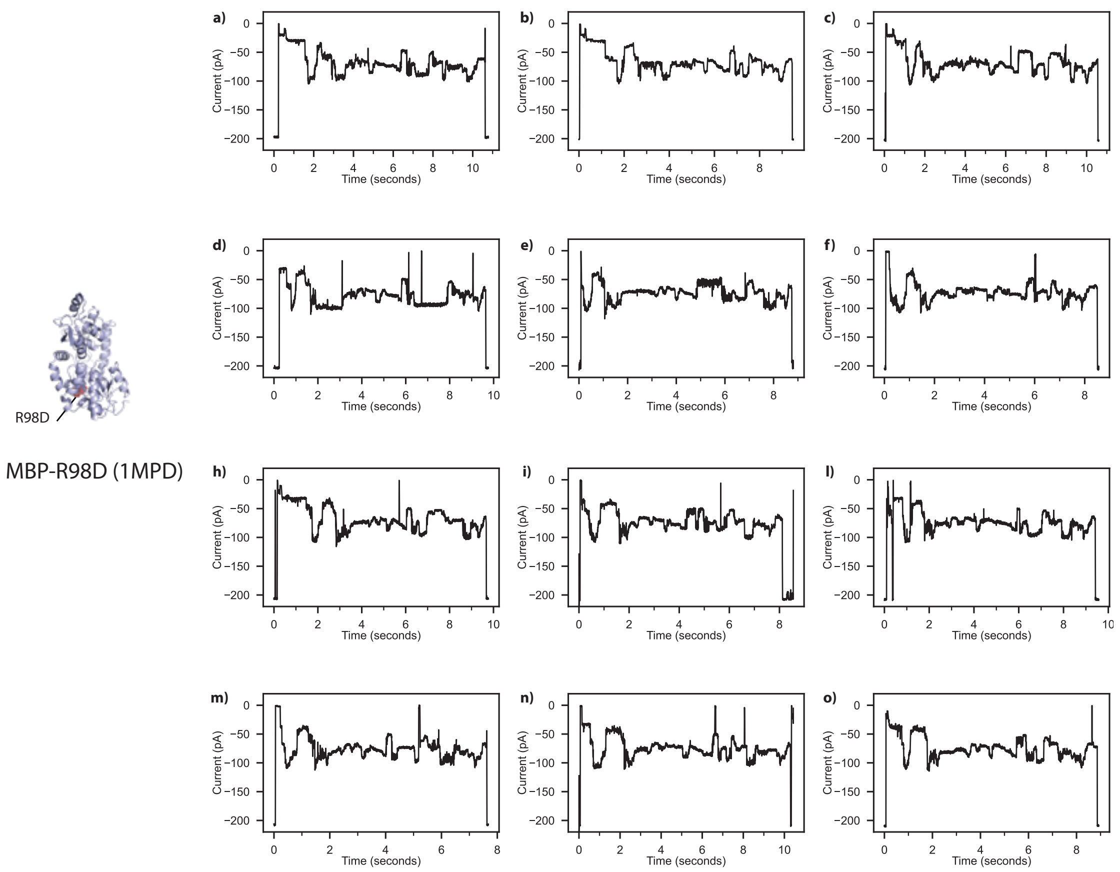


**Figure S30. Representative squiggles of the MBP-R98D-SQtag substrate**. Raw traces were acquired using MspA in the presence of 100 nM of ClpX, 5 nM of POI, with a voltage of -65 mV applied at 37°C in 1 M K-gluconate, 50 mM HEPES, 10 mM $MgCl_2$, pH 7.4, 2 mM of ATP, 1.6 mM creatine phosphate, 0.4 µM creatine kinase, 1 mM DTT, and 0.5 mM EDTA. (a-o) Traces were filtered using a low-pass Bessel filter (500Hz) for enhanced visualization.

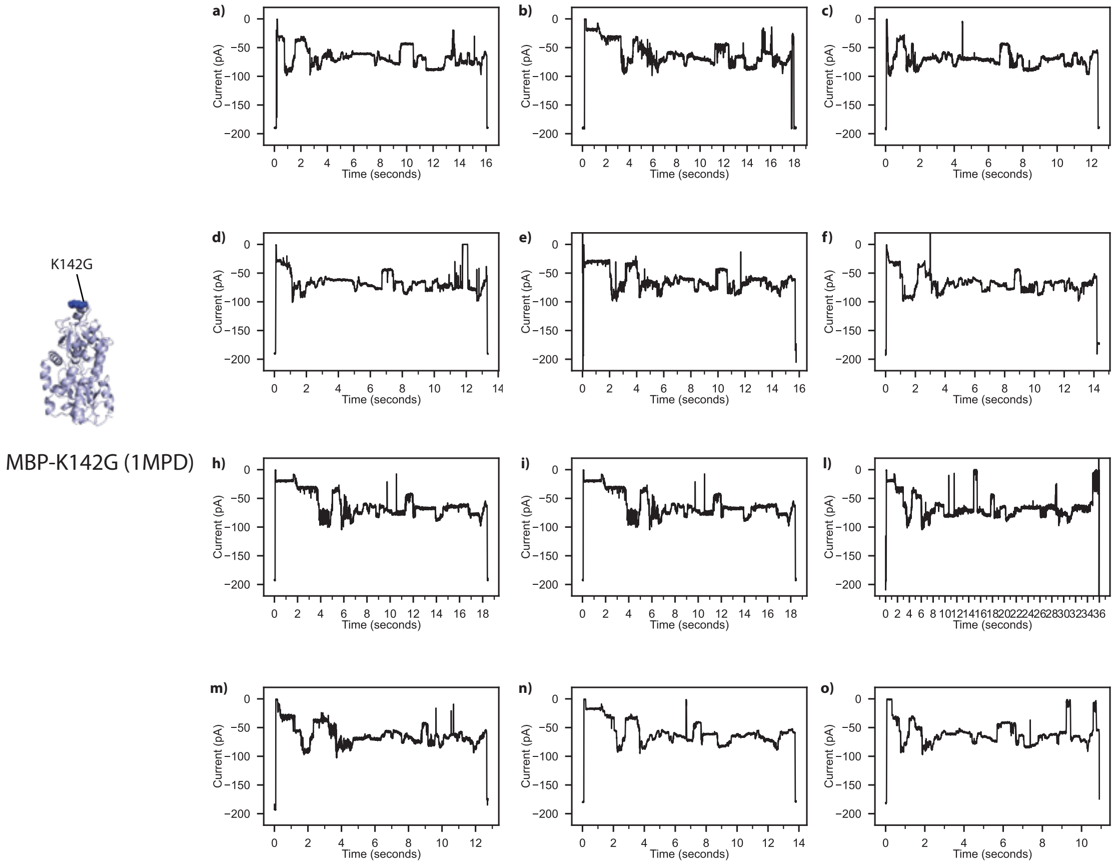


**Figure S31. Representative squiggles of the MBP-K142G-SQtag substrate**. Raw traces were acquired using MspA in the presence of 100 nM of ClpX, 5 nM of POI, with a voltage of -65 mV applied at 37°C in 1 M K-gluconate, 50 mM HEPES, 10 mM $MgCl_2$, pH 7.4, 2 mM of ATP, 1.6 mM creatine phosphate, 0.4 µM creatine kinase, 1 mM DTT, and 0.5 mM EDTA. (a-o) Traces were filtered using a low-pass Bessel filter (500Hz) for enhanced visualization.

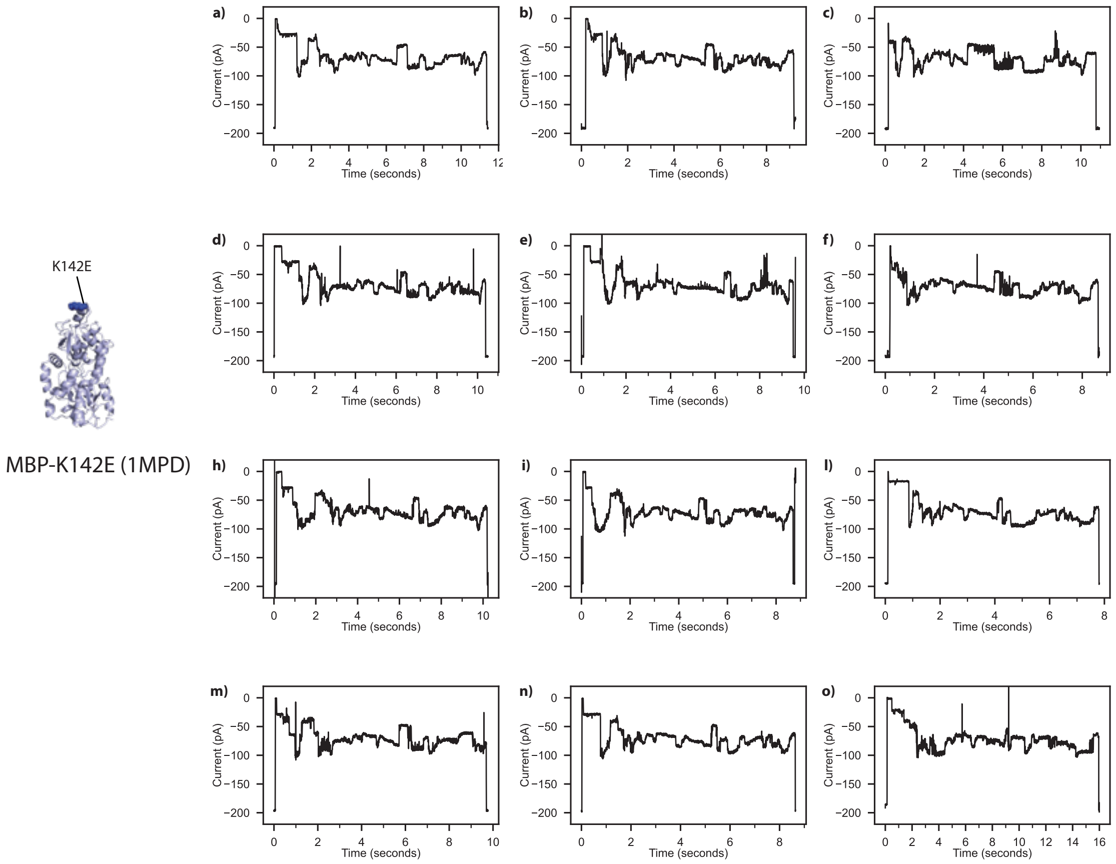


**Figure S32. Representative squiggles of the MBP-K142E-SQtag substrate**. Raw traces were acquired using MspA in the presence of 100 nM of ClpX, 5 nM of POI, with a voltage of -65 mV applied at 37°C in 1 M K-gluconate, 50 mM HEPES, 10 mM $MgCl_2$, pH 7.4, 2 mM of ATP, 1.6 mM creatine phosphate, 0.4 µM creatine kinase, 1 mM DTT, and 0.5 mM EDTA. (a-o) Traces were filtered using a low-pass Bessel filter (500Hz) for enhanced visualization.

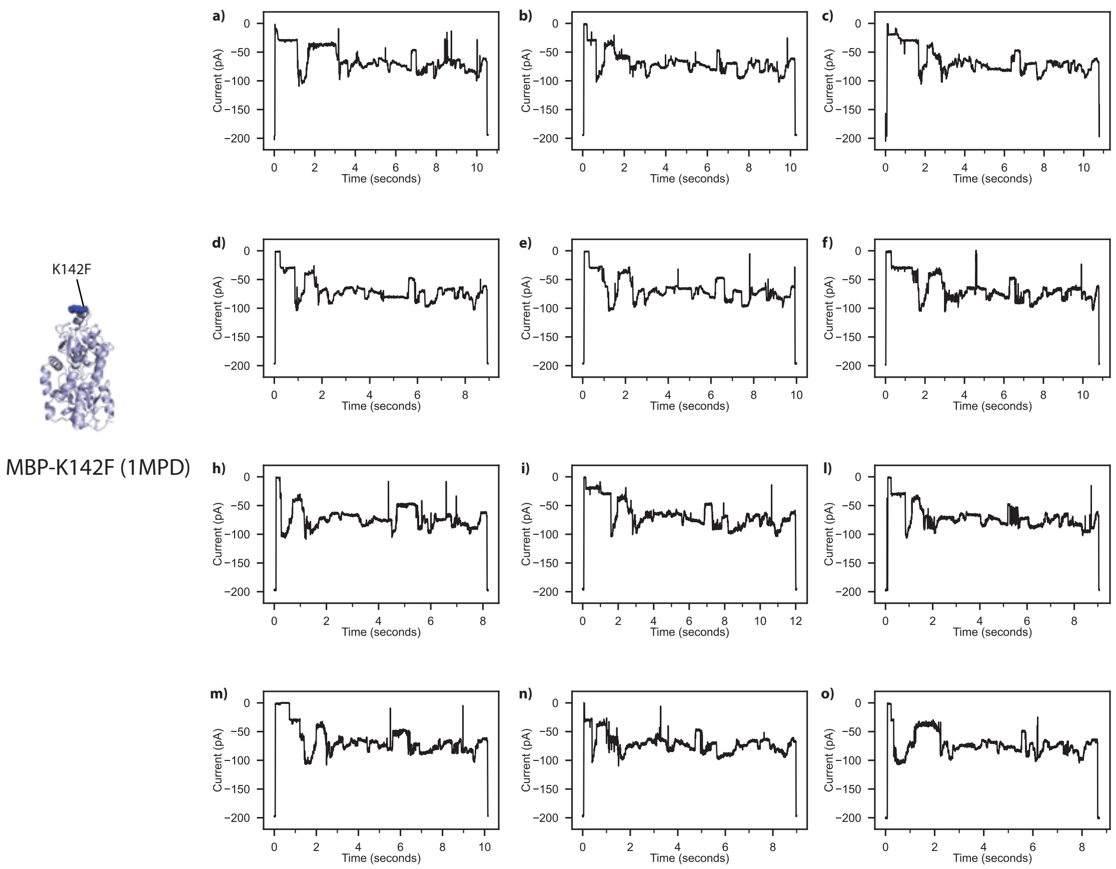


**Figure S33. Representative squiggles of the MBP-K142F-SQtag substrate**. Raw traces were acquired using MspA in the presence of 100 nM of ClpX, 5 nM of POI, with a voltage of -65 mV applied at 37°C in 1 M K-gluconate, 50 mM HEPES, 10 mM $MgCl_2$, pH 7.4, 2 mM of ATP, 1.6 mM creatine phosphate, 0.4 µM creatine kinase, 1 mM DTT, and 0.5 mM EDTA. (a-o) Traces were filtered using a low-pass Bessel filter (500Hz) for enhanced visualization.

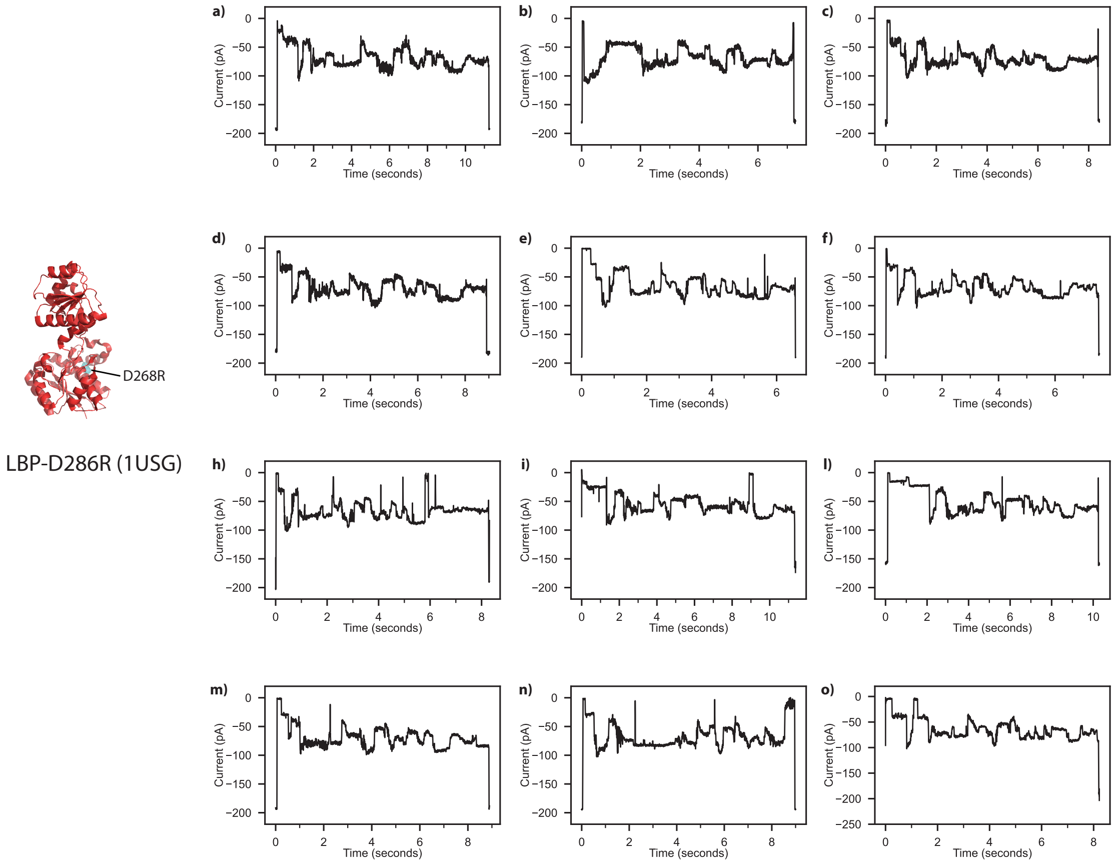


**Figure S34. Representative squiggles of the LBP-D286R-SQtag substrate**. Raw traces were acquired using MspA in the presence of 100 nM of ClpX, 5 nM of POI, with a voltage of -65 mV applied at 37°C in 1 M K-gluconate, 50 mM HEPES, 10 mM $MgCl_2$, pH 7.4, 2 mM of ATP, 1.6 mM creatine phosphate, 0.4 µM creatine kinase, 1 mM DTT, and 0.5 mM EDTA. (a-o) Traces were filtered using a low-pass Bessel filter (500Hz) for enhanced visualization.

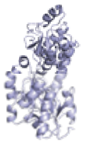

MBP (1MPD)

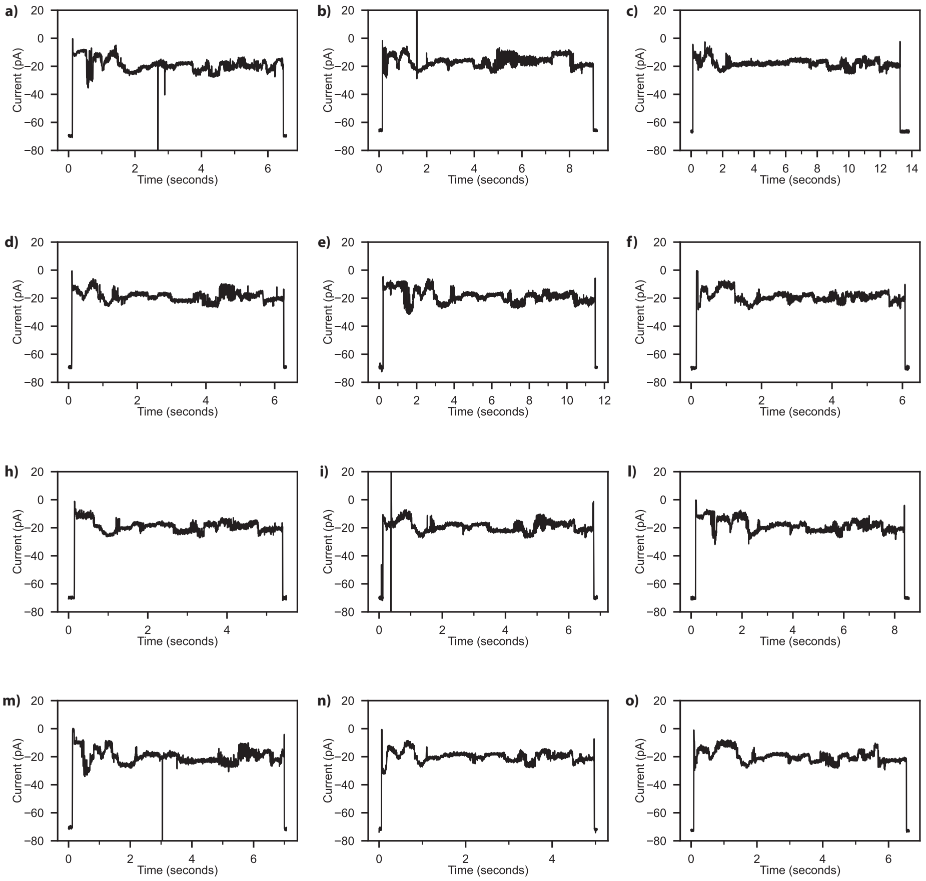


**Figure S35. Representative squiggles of the MBP-SQtag substrate**. Raw traces were acquired using CytK-4D in the presence of 100 nM of ClpX, 5 nM of POI, with a voltage of -65 mV applied at 37°C in 1 M K-gluconate, 50 mM HEPES, 10 mM $MgCl_2$, pH 7.4, 2 mM of ATP, 1.6 mM creatine phosphate, 0.4 µM creatine kinase, 1 mM DTT, and 0.5 mM EDTA. (a-o) Traces were filtered using a low-pass Bessel filter (500Hz) for enhanced visualization.

References.


1. Motone, K. *et al.* Multi-pass, single-molecule nanopore reading of long protein strands. *Nature* **633**, (2024).
2. Yan, Shuanghong, et al. Single molecule ratcheting motion of peptides in a Mycobacterium smegmatis porin A (MspA) nanopore. Nano letters 21.15, (2021).
3. Fei, Xue, et al. "Structures of the ATP-fueled ClpXP proteolytic machine bound to protein substrate." *elife* 9 (2020): e52774.
